# Architecture of the vertebrate egg coat and structural basis of the ZP2 block to polyspermy

**DOI:** 10.1101/2023.06.21.544075

**Authors:** Shunsuke Nishio, Chihiro Emori, Benjamin Wiseman, Dirk Fahrenkamp, Elisa Dioguardi, Sara Zamora-Caballero, Marcel Bokhove, Ling Han, Alena Stsiapanava, Yonggang Lu, Mayo Kodani, Rachel E. Bainbridge, Kayla M. Komondor, Anne E. Carlson, Michael Landreh, Daniele de Sanctis, Shigeki Yasumasu, Masahito Ikawa, Luca Jovine

**Affiliations:** Department of Biosciences and Nutrition, Karolinska Institutet, Huddinge, Sweden; Department of Experimental Genome Research, Research Institute for Microbial Diseases, Osaka University, Suita, Osaka, Japan; Immunology Frontier Research Center, Osaka University, Suita, Osaka, Japan; Graduate School of Pharmaceutical Sciences, Osaka University, Suita, Osaka, Japan; Department of Biological Sciences, University of Pittsburgh, Pittsburgh, PA, U.S.A.; Department of Microbiology, Tumor and Cell Biology, Karolinska Institutet, Stockholm, Sweden; Department of Cell and Molecular Biology, Uppsala University, S-75124 Uppsala, Sweden; ESRF - The European Synchrotron, Grenoble, France; Department of Materials and Life Sciences, Faculty of Science and Technology, Sophia University, Tokyo, Japan; Center for Infectious Disease Education and Research (CiDER), Osaka University, Suita, Osaka, Japan; Institute of Fermentation Sciences (IFeS), Faculty of Food and Agricultural Sciences, Fukushima University, Fukushima, Japan; Department of Biochemistry and Biophysics, Stockholm University, Stockholm, Sweden; Crelux GmbH, Martinsried, Germany; Chiesi Pharma AB, Stockholm, Sweden; Instituto de Biomedicina de Valencia (IBV-CSIC), Valencia, Spain; Japan Synchrotron Radiation Research Institute (JASRI), Hyogo, Japan

**Keywords:** Fertilization, egg-sperm interaction, block to polyspermy, egg coat hardening, zona pellucida, ZP module, ZP2 cleavage, ovastacin, protein filament, integrative structural biology

## Abstract

Post-fertilization cleavage of glycoprotein ZP2, a major subunit of egg zona pellucida (ZP) filaments, is crucial for mammalian reproduction by irreversibly blocking polyspermy. ZP2 processing is thought to inactivate a sperm-binding activity located upstream of the protein’s cleavage site; however, its molecular consequences and connection with ZP hardening are unknown. Here we report X-ray crystallographic, cryo-EM and biochemical studies showing that cleavage of ZP2 triggers its oligomerization. Deletion of the ZP-N1 domain that precedes the cleavage site of mouse ZP2 allows it to homodimerize even without processing, and animals homozygous for this variant are subfertile by having a semi-hardened ZP that allows sperm attachment but hinders penetration. Combined with the structure of a native egg coat filament, which reveals the molecular basis of heteromeric ZP subunit interaction, this suggests that oligomerization of cleaved ZP2 cross-links the ZP, rigidifying it and making it physically impenetrable to sperm.

## INTRODUCTION

By interfering with the generation of a diploid zygote, fertilization of an egg by more than one sperm is lethal for most animals, including humans. Whereas prevention of the fusion of multiple sperm with the egg plasma membrane is achieved using different mechanisms and is often only transient, an evolutionarily conserved and permanent block to polyspermy is established by the irreversible modification of the egg coat (called ZP in mammals and vitelline envelope (VE) in non-mammals)^1,2^. In mammals and anuran amphibians, this depends on site-specific cleavage of ZP/VE subunit ZP2 by a cortical granule (CG) activity released from the egg after fertilization^3,4^, which was identified as zinc metalloprotease ovastacin (Ovst) in the mouse^5^.

ZP2 is a multidomain glycoprotein consisting of a large N-terminal region (NTR) and a C-terminal ZP “domain” (Figure 1A). While the latter is a polymerization module composed of two distinct immunoglobulin-like domains (ZP-N and ZP-C) connected by an interdomain linker (IDL)^6^, the ZP2 NTR plays a key role in the regulation of gamete interaction and was predicted to contain three additional tandem copies of the ZP-N domain (ZP2-N1-3)^7,8^. This hypothesis was corroborated by the crystal structure of mouse ZP2-N1^9^, a domain that is present in mature mammalian ZP2 but proteolytically removed during maturation of anuran ZP2^4^. Despite this difference, in both mammals and amphibians the following ZP2-N2 domain harbors a conserved recognition site for the CG protease^4,10^, whose action generates two ZP2 protein fragments that remain covalently attached through an intramolecular disulfide bond^3,10,11^.

**Figure 1.**
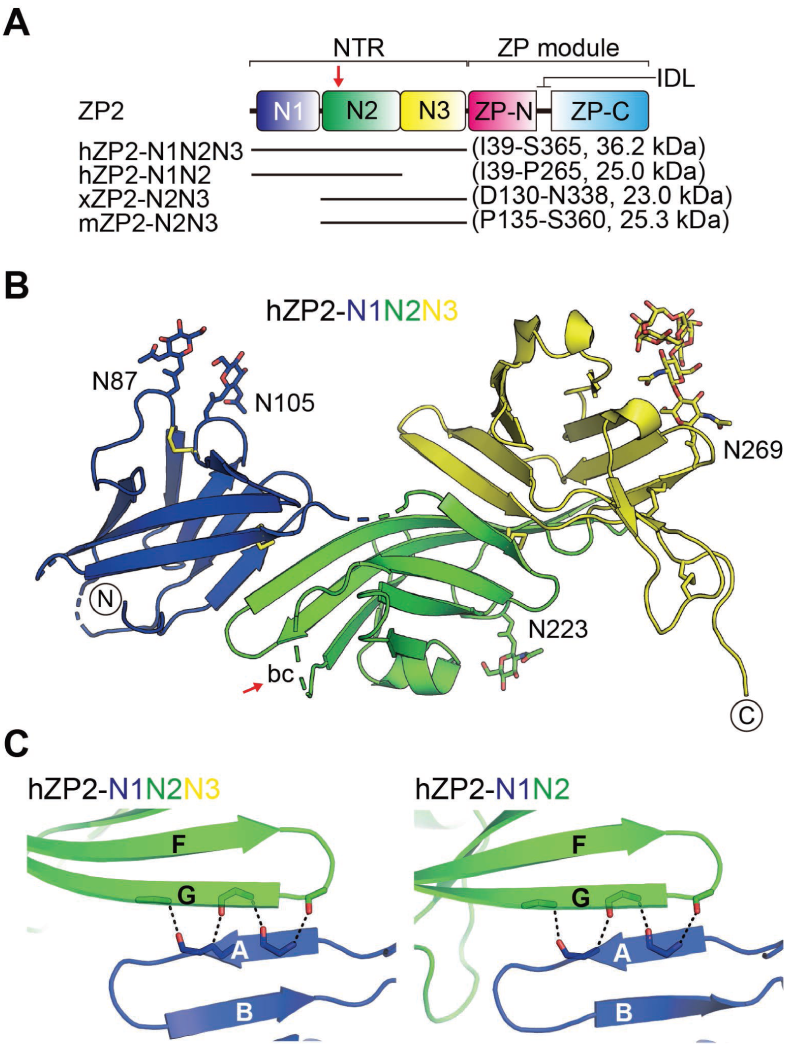
Crystal structures of the functional N-terminal region of human ZP2 in its uncleaved, monomeric state. (A) ZP2 domain architecture. Red arrow, post-fertilization cleavage site within the bc loop of ZP2-N2; horizontal lines, protein regions whose crystallographic structure was determined in this work. hZP2, human ZP2; xZP2, *Xenopus* ZP2; mZP2, mouse ZP2. (B) Crystal structure of hZP2-N1N2N3. Domains are colored as in panel A, with N-linked carbohydrates shown as sticks labeled according to the Asn residues to which they are attached. Protein termini are marked and disordered loops are represented by dashed lines, with the ZP2-N2 bc loop targeted by ovastacin indicated by a red arrow. Disulfide bonds are depicted as yellow sticks. (C) The crystal structures of hZP2-N1N2N3 and hZP2-N1N2 show a conserved intramolecular ZP2-N1/ZP2-N2 domain interface. β-strands are indicated by letters following standard ZP-N nomenclature^40^; dashed lines indicate hydrogen bonds. See also Figures S1 and S2 and Table S1.

Tandem ZP-N repeats are also found at the N-terminus of other egg coat components that are absent in mammals, including the ZPAX subunits of the avian, amphibian and fish VE as well as vitelline envelope receptor for lysin (VERL), a sperm-binding VE component of the mollusk abalone^9,12^. On the other hand, due to its key role in protein polymerization^13^, the ZP module is conserved in all the subunits of the egg coat of vertebrates (such as mouse ZP1-3 and human ZP1-4), the often much more numerous components of the VE of many invertebrates (including abalone VERL and VEZP2-30), and hundreds of other extracellular molecules from multicellular eukaryotes with highly different biological functions (for example urinary glycoprotein uromodulin (UMOD))^14–16^. Studies on ZP subunits and UMOD showed that incorporation into the respective nascent filaments depends on processing of their membrane-bound precursors at a cleavage site (referred to as consensus furin cleavage site (CFCS) in ZP proteins) located after the ZP module^17,18^. This triggers the dissociation of a polymerization-blocking external hydrophobic patch (EHP) propeptide, which in the precursors of ZP3, ZP2 and UMOD acts as β-strand G of the ZP-C fold^6,19,20^. Recent cryo-EM studies showed that the homomeric filament of human UMOD adopts a unique architecture that results from major conformational changes of the ZP module during polymerization^21,22^. However, there is no detailed structural information on any heteromeric ZP module protein filament, and the molecular basis of how ZP subunits interact with each other to form the egg coat remains unknown.

Post-fertilization cleavage of ZP2 was initially hypothesized to abolish sperm binding by either altering the conformation of ZP proteins or somehow modifying the overall architecture of the egg coat^10,11^, whereas in the current model of mammalian gamete recognition it is postulated to act by inactivating a direct sperm-binding site on ZP2-N1^23,24^. At the same time, processing of ZP2 has long been connected with ZP hardening, a global physicochemical change of the egg coat matrix associated with significant increases in stiffness, resistance to proteolytic digestion and filament density^11,25^. Considering that gamete recognition and ZP hardening are intimately connected with the NTR and ZP module moieties of ZP2, respectively, we combined studies of the protein’s ZP-N repeats and egg coat architecture to bridge the aforementioned observations and understand how ZP2 cleavage regulates sperm binding at the molecular level.

## RESULTS

### Structure of intact, monomeric ZP2 NTR

To investigate the structure of ZP2 NTR in its intact (pre-fertilization) and cleaved (post-fertilization) states, we first expressed the complete NTR of human ZP2 (hZP2) in mammalian cells and determined its crystal structure at 2.7 Å resolution by experimental phasing (Table S1). This showed that, rather than being arranged like beads on a string, the three NTR domains generate a V-shaped architecture where ZP2-N1 and ZP2-N3 extend the sheets of the ZP2-N2 β-sandwich (Figure 1B). This is facilitated by the second β-sheet of ZP2-N2, whose F and G strands are significantly longer than those of ZP2-N1/N3. Notably, although ZP2-N2 and ZP2-N3 share the same basic fold as ZP2-N1^9^, their cd loops are not engaged in the typical disulfide bond between conserved Cys 2 and 3 (C_2_-C_3_) found in other ZP-N domains^6,8,9,20^, but instead harbor an additional C’ β-strand (as previously observed in ZP1-N1^26^) and a helical region (Figure S1). Whereas ZP2-N2 and -N3 share an interface of ∼770 Å^2^, the interaction between ZP2-N1 and -N2 is less extensive (∼475 Å^2^) but also observed in a 3.2 Å resolution structure of hZP2-N1N2 that we subsequently solved by molecular replacement (MR) (Figure 1C, Figure S2 and Table S1). In both crystals, however, there is no density for the ZP-N2 bc loop containing the CG protease cleavage site (171-LA|DD-174), suggesting that this is highly flexible in uncleaved ZP2.

### Cleavage triggers ZP2 NTR oligomerization

Although the anuran CG protease remains to be identified, collagenase can closely mimic its action by both specifically cleaving the bc loop of ZP2-N2 and triggering a block to sperm binding^4,27^. Based on this observation, we treated preparative amounts of recombinant *Xenopus* ZP2 (xZP2)-N2N3 with collagenase and discovered that cleavage induces almost complete tetramerization of the protein (Figure 2A-C). The same phenomenon was observed in parallel small-scale experiments performed using a crude exudate from activated *Xenopus* eggs, which however also yielded a roughly equal amount of dimers (Figure 2D-F). Crystallography and cryo-electron microscopy (EM) (Tables S1 and S2) reveal that cleavage of xZP2-N2N3, which is highly similar to the same region of hZP2 (average RMSD 1.9 Å over 153 Cα), allows it to homodimerize through the reorientation and self-interaction of the severed bc loops of two molecules (Figure 2G). These pack on top of paired βG strands (Figure 2H), generating an interface of ∼1630 Å^2^ that — combined with a second type of interface mediated by intermolecular pairing between ZP2-N2 C’ and E’ strands (∼460 Å^2^; Figure 2I) — results in the formation of a highly stable dimer of homodimers (Figures 2J, K and S4). To extend these observations to mammals, we developed a fully recombinant system for *in vitro* cleavage of hZP2 NTR with hOvst (Figure 3A-C). This demonstrated that hOvst directly cleaves hZP2 in a metal-dependent way and showed that hZP2 also homodimerizes upon processing (Figure 3D).

**Figure 2.**
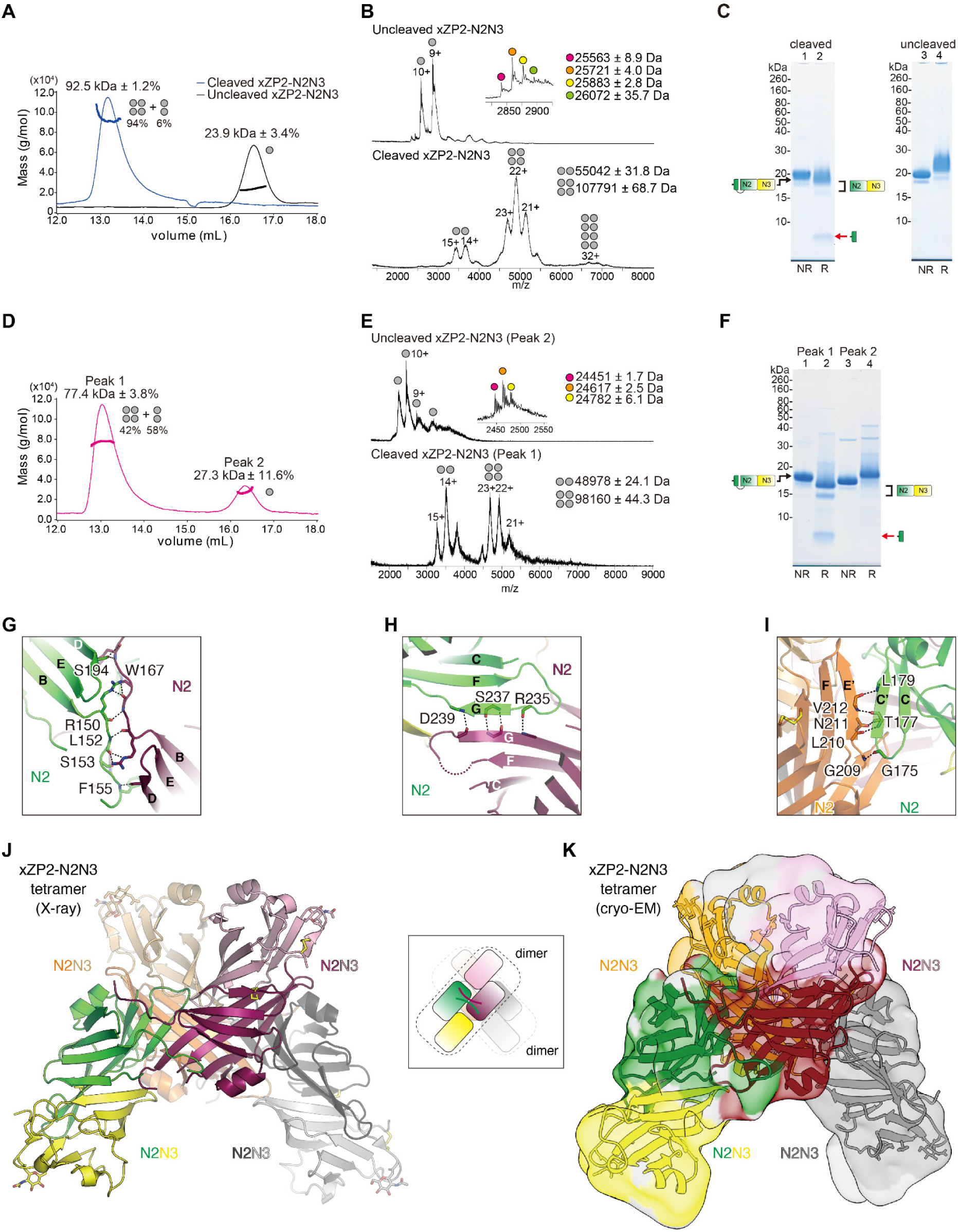
Biochemical and structural analysis of the N-terminal region of *Xenopus* ZP2 in its cleaved, homotetrameric state. (A) Size exclusion chromatography-multi angle light scattering (SEC-MALS) profile of uncleaved and collagenase-cleaved xZP2-N2N3. Gray dots indicate the oligomerization state of the protein. Note that the peaks of protein tetramers and dimers overlap, due to their very similar radiuses of gyration (deduced from the structure shown in panels J, K). (B) Native mass spectrometry (MS) of the samples from panel A. (C) SDS-PAGE analysis of the samples from panel A. The red arrow in this panel and panel F indicates the N-terminal fragment of cleaved xZP2, which separates from the rest of the protein upon reduction of the C_1_139-C_4_244 disulfide. (D) SEC-MALS profile of *Xenopus* egg exudate-treated xZP2-N2N3. (E) Native MS analysis of the material from panel D shows that peak 2 corresponds to uncleaved monomeric xZP2-N2N3 (top panel) whereas peak 1 consists of dimeric and tetrameric forms of the cleaved protein (bottom panel). (F) SDS-PAGE analysis shows that peaks 2 and 1 from panel D correspond to cleaved and uncleaved xZP2-N2N3, respectively. (G-I) Details of ZP2 oligomeric interactions mediated by cleaved loops (G), G β-strands (H) and C’, E’ β-strands (I). (J) Crystal structure of cleaved xZP2-N2N3. Right inset, xZP2-N2N3 tetramer scheme. (K) Cryo-EM map of the xZP2-N2N3 tetramer. See also Figures S3 and S4 and Table S2.

**Figure 3.**
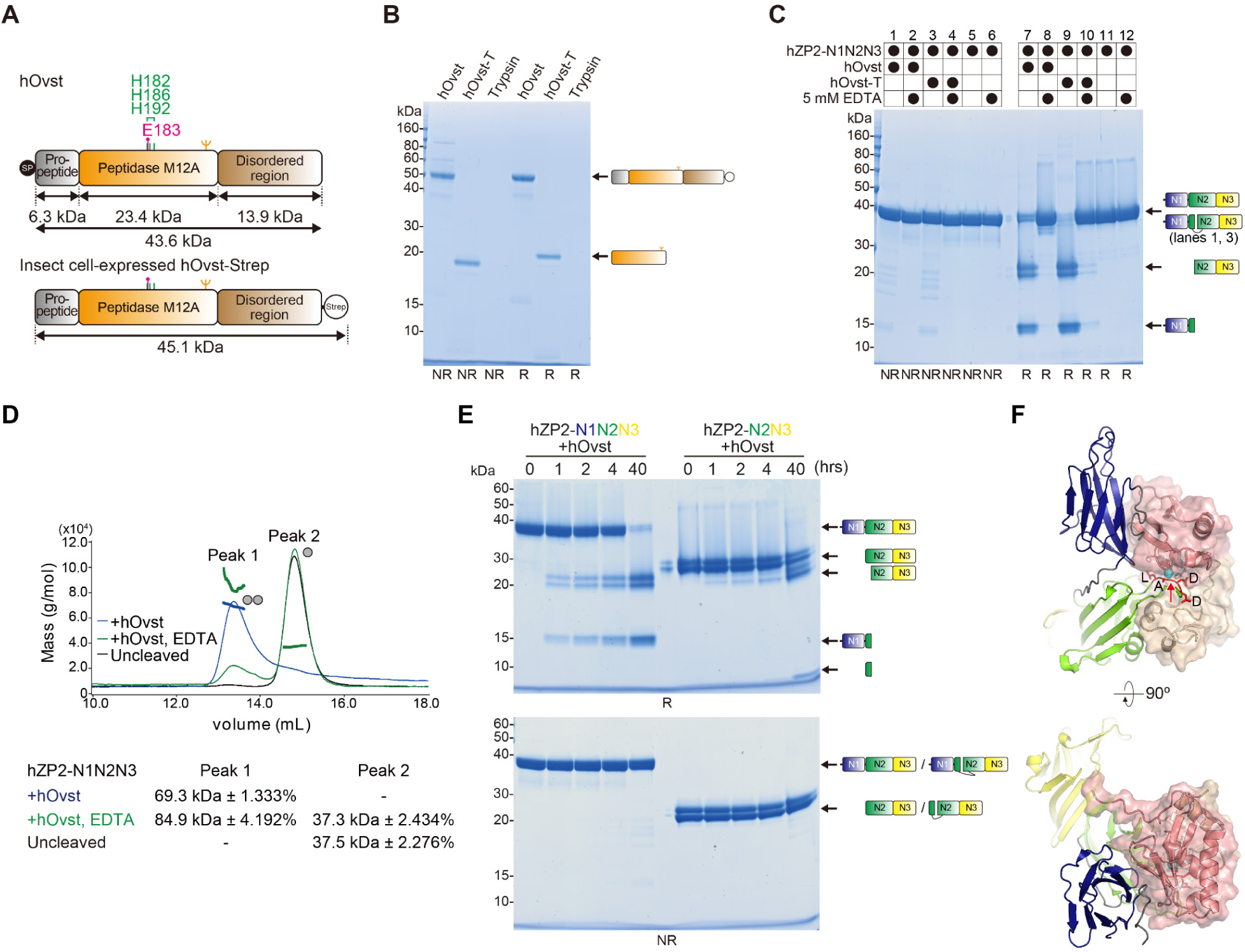
Ovastacin cleavage of the N-terminal region of human ZP2 triggers its homodimerization and is facilitated by the ZP2-N1 domain. (A) Schematic diagram of hOvst. Theoretical molecular weights of domains/regions and positions of residues constituting the zinc-binding HExxHxxGxxH active site motif are indicated. SP, signal peptide. (B) Trypsinization of purified hOvst expressed in S2 insect cells. hOvst-T, trypsinized hOvst. NR, non-reducing conditions; R, reducing conditions. (C) Cleavage of hZP2-N1N2N3 with untrypsinized or trypsinized hOvst. Note how EDTA inhibits the Zn^2+^-dependent proteolytic activity of hOvst. (D) SEC-MALS profile of uncleaved and hOvst-cleaved hZP2-N1N2N3. (E) Time course of hZP1-N1N2N3 or hZP2-N2N3 cleavage with hOvst. (F) Orthogonal views of the top-ranked AlphaFold-Multimer model of the hZP2 NTR/hOvst complex. hZP2 NTR domains are depicted as in Figure 1, with the residues flanking the ovastacin cleavage site (black arrow) shown as sticks and colored red. hOvst is in mixed surface/cartoon representation, with the N- and C-terminal lobes of the astacin-like peptidase domain colored salmon and wheat, respectively; C-terminal residues P283-D431, predicted with low confidence and largely adopting the typical ribbon-like appearance of disordered protein regions^73^, do not make contacts with the hZP2 NTR and are not shown. HExxHxxGxxH motif residues are represented as sticks, with the Zn^2+^ ion depicted as a cyan sphere.

### The ZP-N1 domain regulates cleavage and dimerization of mammalian ZP2

Comparison of cleavage experiments carried out using the complete NTR of hZP2 or a fragment consisting of the ZP2-N2 and ZP2-N3 domains shows that the catalytic efficiency of hOvst is strongly dependent on the presence of the ZP2-N1 domain (Figure 3E), suggesting that the latter facilitates protease/substrate interaction. In agreement with such a function, prediction of the structure of the hZP2 NTR/hOvst complex with AlphaFold-Multimer consistently results in models where ZP2-N1 binds to the N-terminal lobe of hOvst and is displaced from ZP2-N2, which interacts with the C-terminal lobe of the protease and inserts the bc loop LADD sequence into the enzyme’s active site (Figure 3F).

Notably, the ZP2-N2 βG strand that mediates the intermolecular pairing of cleaved xZP2 (Figure 2H) pairs intramolecularly with ZP2-N1 βA in the context of uncleaved hZP2 (Figure 1C). This raises the question of whether, in addition to its suggested role as a sperm receptor, the ZP2-N1 domain influences the post-fertilization cleavage of mammalian ZP2 in ways other than facilitating the action of Ovst. To address this point, we generated mice expressing ZP2 lacking ZP2-N1 instead of the wild-type protein (Figure S5A-C). These animals are severely subfertile *in vivo* (Figure 4A) and sterile *in vitro* (Figure 4B). Unexpectedly, sperm still attach to the ZPs of animals homozygous for this ΔN1 mutation (Figure 4C); however, very few sperm manage to penetrate the mutant ZP (Figure 4D-G), which — despite detectable lack of mZP2 cleavage by mOvst (Figure S5D-H) — is as resistant to collagenase digestion (used as as an indicator for hardening) as that of fertilized wild-type eggs (Figure 4H). In agreement with the latter observation, crystallography of a mouse NTR ΔN1 construct (mZP2-N2N3) revealed that this protein is able to establish a homodimeric interface similar to that of cleaved xZP2-N2N3, even though its N2 bc loop is intact (Figures 4I, J and S5D-H; Table S1).

**Figure 4.**
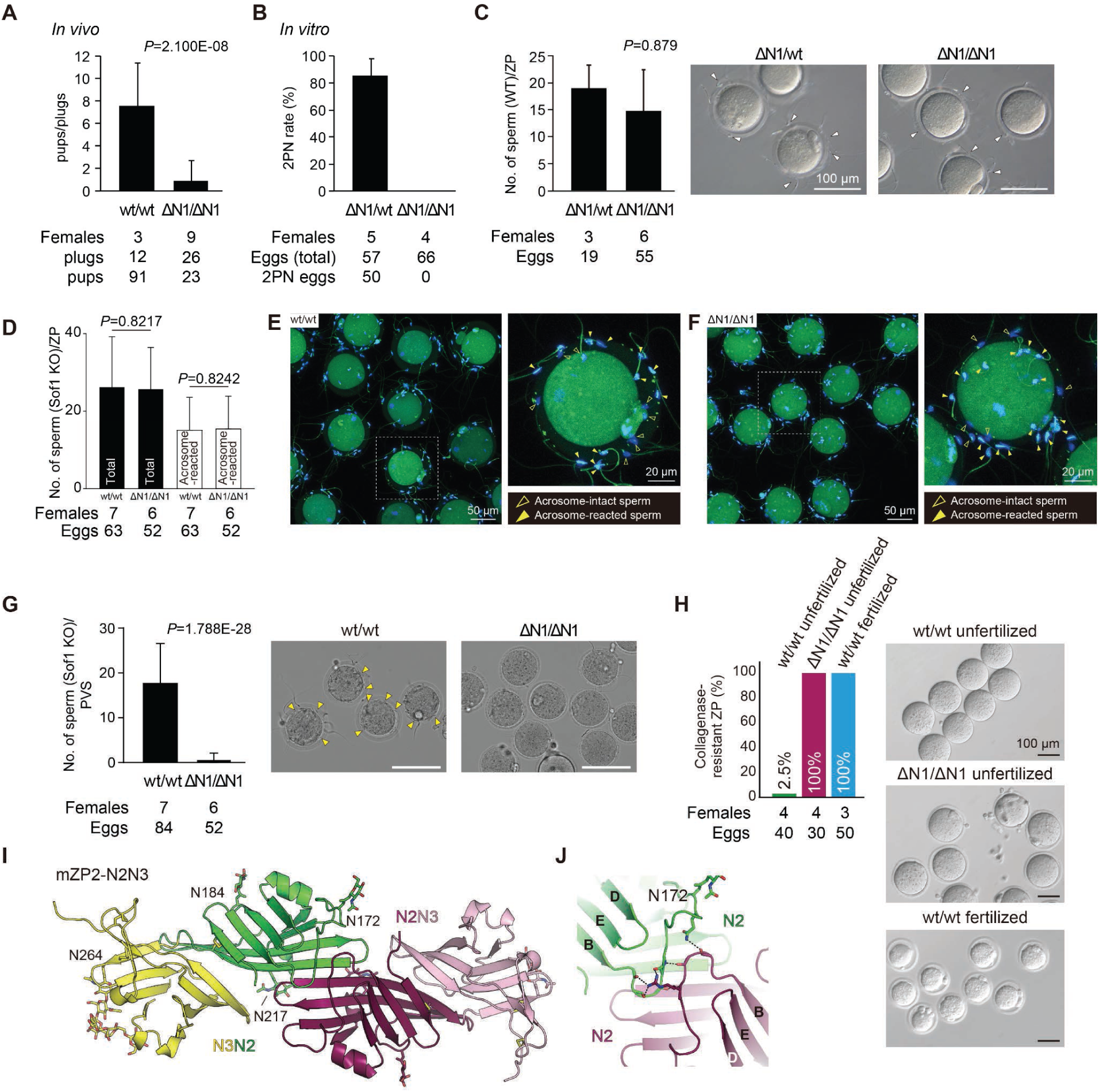
Deletion of ZP2-N1 does not affect sperm binding, but leads to a semi-hardened ZP and subfertility in mice. (A) Number of pups over plugs. (B) *In vitro* fertilization (IVF) percentage. (C) Sperm binding assay. White arrowheads indicate ZP-bound sperm. (D-F) *Sof1* KO sperm bind equally well to the ZP of wt and ZP2 ΔN1 homozygous eggs. (D) Number of *Sof1* KO sperm bound to wt/wt or ΔN1/ΔN1 eggs. *Sof1* KO sperm show comparable binding to the ZP of both kinds of oocytes and their acrosome status does not influence this activity. (E, F) Fluorescence images of *Sof1* KO sperm bound to wt or ΔN1/ΔN1 eggs. To monitor the acrosome status, sperm were pre-stained with anti-Izumo1 antibody. Open and close yellow arrowheads indicate acrosome-intact and -reacted sperm, respectively. (G) Sperm penetration assay. PVS, perivitelline space. Yellow arrowheads indicate sperm in the PVS. (H) Collagenase resistance of eggs from wt or ΔN1/ΔN1 mice. (I) The crystal structure of mZP2-N2N3 shows a homodimer. (J) Detail of the interaction between uncleaved mZP2-N2 bc loops. See also Figure S5.

Taken together, these results indicate that, unlike previously thought, ZP2-N1 is not required for sperm attachment to the mouse ZP; however, this domain is crucial for fertility by physically counteracting the association of uncleaved ZP2 molecules before fertilization, as well as facilitating their cleavage by Ovst after fertilization.

### Three-dimensional architecture of a vertebrate egg coat filament

Because ZP2 is a multi-domain protein whose NTR and ZP module halves are involved in the regulation of gamete interaction and polymerization, respectively (Figure 1A), the biological significance of our NTR-related findings is intimately connected with the essential role of this molecule as a building block of the egg coat^16,28^. As mentioned above, cryo-EM recently unveiled the homopolymeric architecture of urinary ZP module protein uromodulin (UMOD)^21^. However, comparable studies of native heteromeric ZP filaments have been hindered by their paucity, covalent cross-linking and extreme biochemical heterogeneity; moreover, there is currently no system to reconstitute these polymers *in vitro* from recombinant subunits. To overcome this impasse, we exploited the fact that relatively homogeneous native egg coat filament fragments can be obtained upon specific digestion of medaka fish VE with hatching enzymes^21,29^. Using this material, we crystallized a ∼74 kDa native heteromeric complex consisting of an intact ZP module from the fish homolog of ZP3 (fZP3) and single copies of the ZP-N and ZP-C domains of fish ZP1 (fZP1) — a trefoil domain-containing VE subunit whose ZP module closely resembles those of ZP1, ZP2 and ZP4 proteins from other vertebrates, and whose IDL is specifically cleaved during hatching. Although the ∼4 Å resolution diffraction of the crystals hampered experimental phasing, we could solve the structure by MR using a partial AlphaFold model of the fZP3/fZP1 complex, generated using a procedure developed to investigate UMOD polymerization *in silico*^30^ (crystal form I, Figure S6A, B and Table S3). The resulting refined model was then used for MR phasing of data collected from a different crystal form of the same material, which diffracted highly anisotropically beyond 2.7 Å resolution (crystal form II, Figure S6D-H and Table S3).

The maps provide the first detailed information on the polymeric structure of egg coat proteins by revealing a common rod-shaped assembly where fZP3 adopts a fully extended conformation, with its IDL embracing the ZP-C and ZP-N domains of preceding and subsequent fZP1 subunits, respectively (Figure 5A-C). However, whereas crystal form I contains dimers of fZP3/fZP1 complexes held together by weak head-to-head contacts (Figure S6B, C), crystal form II consists of pseudo-infinite filaments where complexes are engaged in extensive head-to-tail interfaces (Figure 5B and Figure S6H). Crucially, the latter position the hatching enzyme-cleaved termini of fZP1-C and fZP1-N moieties at a distance that is compatible with the covalent connectivity of intact fZP1 subunits (Figure 5B, red arrows). This strongly suggests that the packing of crystal form II — which will be henceforth discussed — is biologically relevant and recapitulates a filament architecture where two protofilaments made up of fZP3 and fZP1 subunits, respectively, wrap around each other into a left-handed double helix (Figure 5C).

**Figure 5.**
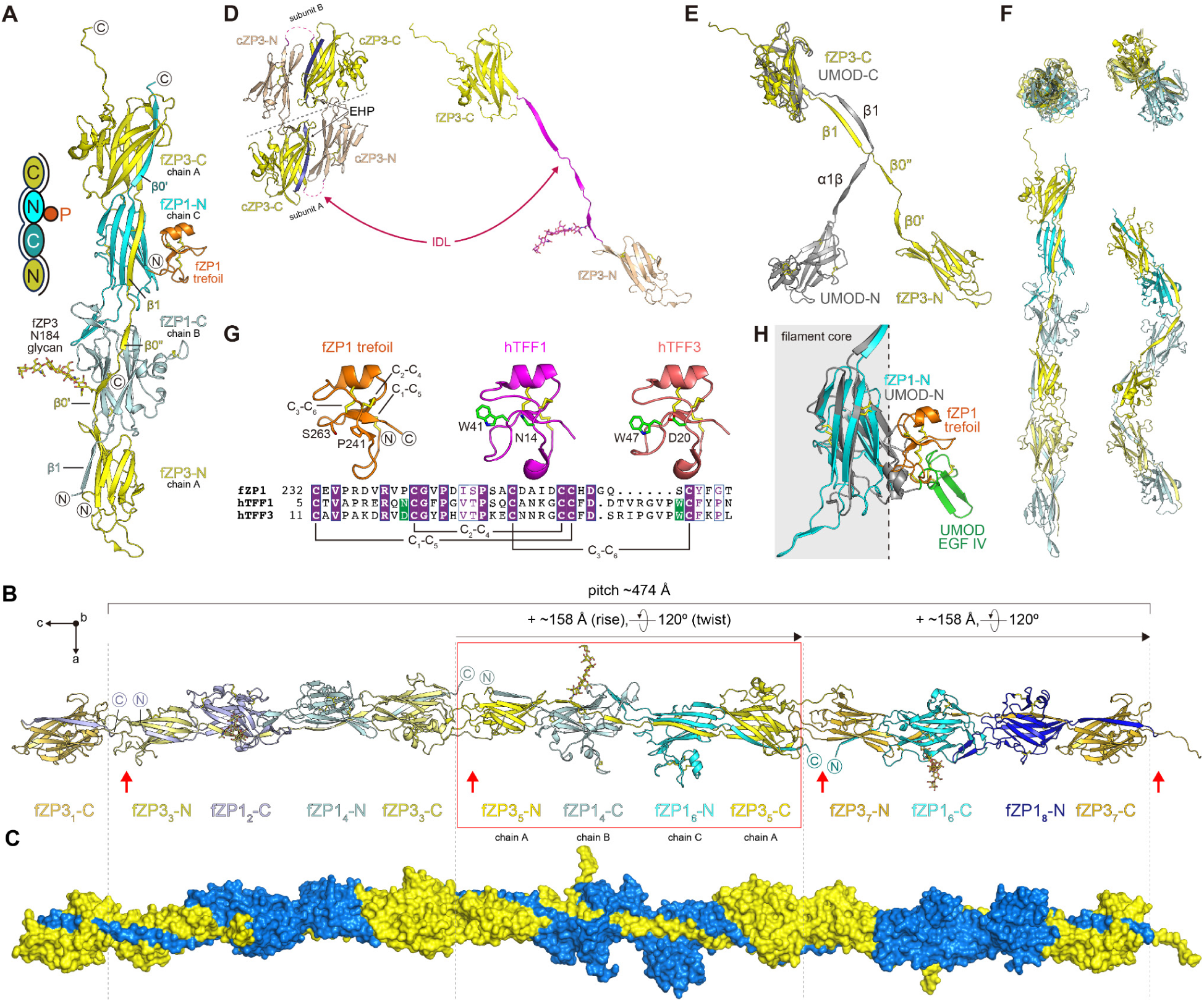
Structure of a native fish VE fragment crystallized as a filament. (A) Overall structure of the fZP3/fZP1 complex (crystal form II). N, C and P in the scheme on the left indicate ZP-N, ZP-C and trefoil domains, respectively. (B) ac plane view of crystal form II. The fZP3-N and fZP3-C domains of complexes contained in adjacent asymmetric units interact head-to-tail, bringing the C- and N-terminal ends of cleaved fZP1 IDLs in close proximity to one another (red arrows). This regenerates pseudo-infinite helical egg coat filaments, only one of which is depicted here, that run parallel along the c axis. Axes directions are indicated on the top left. Asymmetric unit boundaries are indicated by vertical dashed lines; the second of the three complexes that make up a complete helical turn, according to the parameters reported on top, is enclosed by a red rectangle and colored as in panel A. Protein subunits are numbered according to their order in the filament. (C) Surface representation of the egg coat filament shown in panel B, with fZP3 and fZP1 subunits colored yellow and blue, respectively. To highlight how the IDLs of each subunits wrap around the ZP-C and ZP-N domains of the other, cleaved fZP1 linker moieties have been reconnected *in silico*. (D) Structural comparison of the homodimeric precursor of cZP3^6^ and polymeric fZP3. The ZP-N/ZP-C linker is largely disordered before polymerization, which requires cleavage and dissociation of the C-terminal EHP^19,74^. (E) Superposition of the fZP3 subunit from the heterocomplex and a subunit of the UMOD homopolymer. (F) Comparison of the fish VE and UMOD filament cores in section (top) and side (bottom) views. VE and corresponding UMOD subunits are depicted using the same color scheme as panel B; non-ZP module domains are omitted. (G) Structure of the trefoil domain of fZP1 and its comparison with hTFF1 and hTFF3^75^ (RMSD over 36 Cα 1.05 Å and 1.06 Å, respectively). While the fZP1 trefoil domain shares the same disulfide pattern as hTFF1 and hTFF3, it lacks both of their conserved sugar-binding residues (green).Z (H) The NTRs of polymeric fZP1 and UMOD protrude similarly from the respective filaments. See also Figure S6 and Table S3.

Although the conformation of the ZP module of the precursor forms of ZP1/2/4 is unknown, such information is available for ZP3, whose precursors can be secreted as either obligatory or facultative homodimers^6,21,31^. Comparison of the structure of the homodimeric precursor of chicken ZP3 (cZP3) with that of polymeric fZP3 indicates that, although its precursor is very different from that of UMOD^6,20^, ZP3 also undergoes a major conformational rearrangement during polymerization which stretches its ZP-N/ZP-C linker for interaction with other subunits (Figure 5D). As a result, although the heteromeric egg coat polymer has significantly different helical parameters than homopolymeric UMOD^21^ and lacks its characteristic zig-zag pattern (Figure 5E, F), the two types of filaments share the same overall architecture. Accordingly, as observed in the case of UMOD epidermal growth factor (EGF) domain IV^21^, the trefoil domain of fZP1 (Figure 5G) — which is conserved in all ZP1/ZP4 homologs and corresponds to ZP2 NTR by immediately preceding the ZP-N domain — protrudes from the core of the filament (Figure 5H).

### Molecular basis of egg coat subunit heteropolymerization

Consistent with the suggested mechanism of UMOD assembly^21^, the egg coat heterofilament has a polarity dictated by the interaction of the ZP-N domain of an incoming subunit (n) with the ZP-C domain of the second to last one (n-2) component of a growing polymer. Combined with the two-protofilament architecture (Figure 5C), this gives rise to a linear alternation of homomeric (ZP1/ZP1 or ZP3/ZP3) ZP-C/ZP-N interfaces and heteromeric (ZP3/ZP1 or ZP1/ZP3) ZP-N/ZP-C interfaces (Figure 6A).

**Figure 6.**
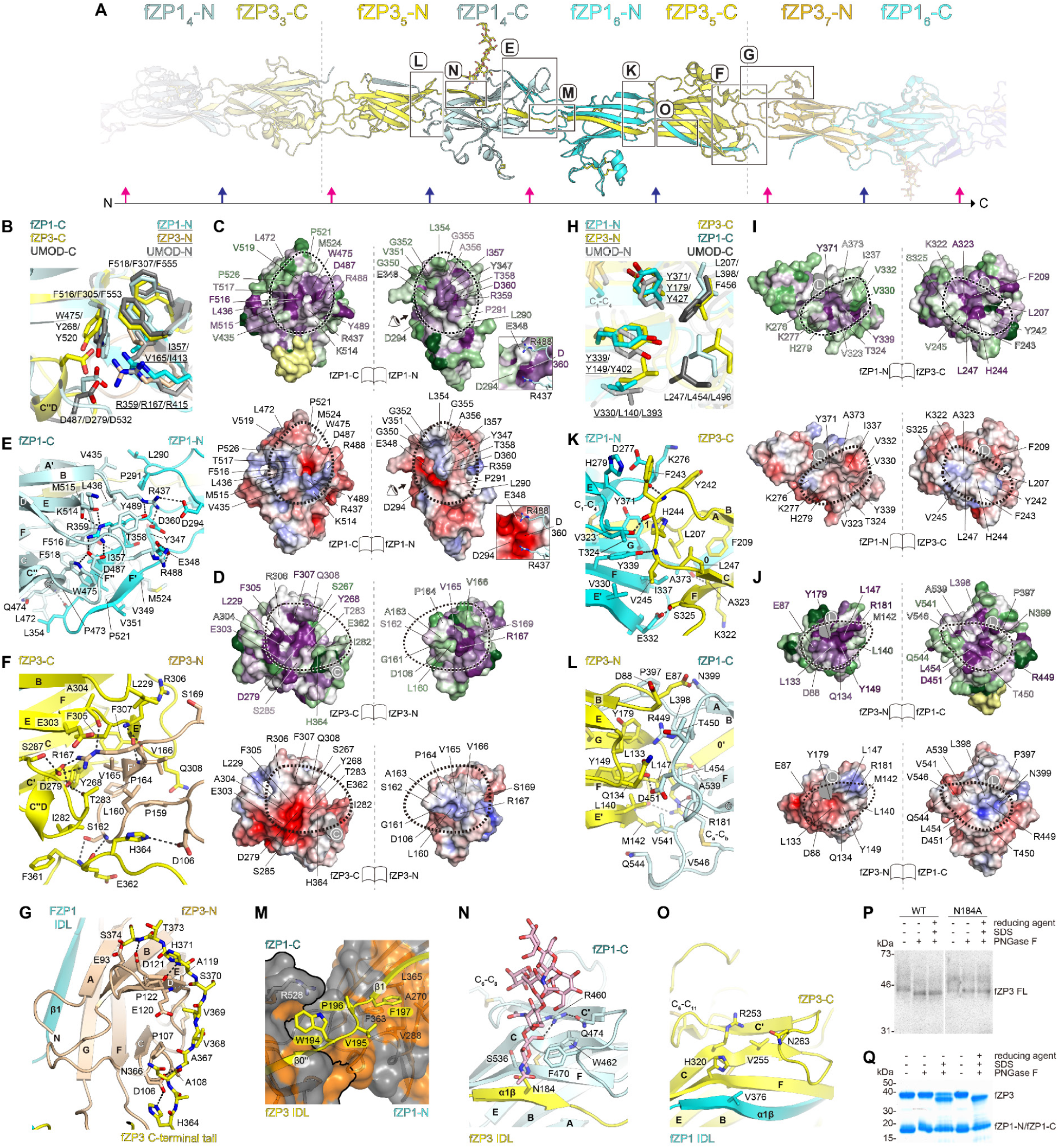
Analysis of protein-protein interactions regulating the assembly of the fZP1/fZP3 heteropolymer. (A) Enlarged view of the central section of the egg coat filament shown in Figure 5B. Different contact areas between ZP subunits are enclosed by boxes, whose labels refer to the following panels. At the bottom, a black horizontal arrow indicates the filament polarity, with vertical magenta and dark blue arrows highlighting the alternation of homomeric and heteromeric interfaces, respectively. (B) Five highly conserved residues mediate homomeric ZP-C/ZP-N domain interactions in both the fZP1 and fZP3 protofilaments, as well as the UMOD polymer. ZP-N residues are underlined. (C) Surface representations of the fZP1-C/fZP1-N homomeric interface colored by evolutionary conservation of amino acid positions in ZP1 and the related ZP modules of ZP2 and ZP4 (using a spectrum that ranges from deep magenta (highest conservation) to light green (lowest conservation), with yellow indicating residues with a low-confidence conservation level; top), or by electrostatic potential (using a spectrum that ranges from red (2.5 kT/e) to blue (+2.5 kT/e) through white (0 kT/e); bottom). Panels are shown in an open book representation, with black dashed ovals indicating the moieties of the interface. The inset is a close-up view of the fZP-N1 surface from the direction of the eye symbol, highlighting charge-related interactions between R437 and R488 of fZP1-C and D294, E348 and D360 of fZP1-N. (D) Amino acid conservation in ZP3 homologs and electrostatic potential surface of the homomeric fZP3-C/fZP3-N interface. The gray area is a cross-section of the fZP3 C-terminal tail, indicated by the symbol “C”. (E, F) fZP1-C/fZP1-N (E) and fZP3-C/fZP3-N (F) homomeric interface contacts. In these and subsequent panels, black dashed lines indicate hydrogen bonds and salt bridges. (G) The C-terminal tail of fZP3 interacts with the ZP-N domain of the following subunit within the fZP3 protofilament. Notably, the same C-terminal residues are completely disordered in crystal form I, consistent with the idea that its packing is non-biologically-relevant. (H) Heteromeric ZP-N/ZP-C interactions are mediated by five highly conserved residues sharing similar relative positions in the fZP1-N/fZP3-C, fZP3-N/fZP1-C and UMOD-N/UMOD-C interfaces. (I, J) Amino acid conservation and electrostatic potential surface of the heteromeric fZP1-N/fZP3-C (I) and fZP3-N/fZP1-C (J) interfaces. IDL cross-sections are in gray and indicated by the symbol “L”. (K, L) fZP1-N/fZP3-C (K) and fZP3-N/fZP1-C (L) homomeric interface contacts. (M) Detail of part of the fZP3 IDL wrapping around the fZP1-C/fZP1-N homomeric interface. The fZP1-C/fZP1-N boundary is marked by a thick black outline; the surface of hydrophobic and non-hydrophobic fZP1 residues is colored orange and gray, respectively. (N) The N-glycan attached to the invariant Asn residue of fZP3 IDL’s α1β interacts with R460, W462 and F470 of the ZP-C domain of the preceding fZP1 subunit. (O) View of the fZP1 α1β/fZP3-C interface, inverse of the one shown in panel N. (P) Immunoblot analysis of secreted recombinant wild-type fZP3 and N-glycosylation site mutant N184A, treated with PNGase F under native or fully denaturing conditions. (Q) PNGase F treatment of the fZP1/fZP3 complex under native, partially denaturing or fully denaturing conditions. See also Figure S6.

The homomeric interfaces share a set of six highly conserved residues (Figure 6B-D), which are also present in UMOD despite a much more kinked ZP-C/ZP-N orientation (Figure 5F) stabilized by additional contacts (ZP-C ef loop/ZP-N cd loop and Y407) not found in the straight egg coat filament. Notably, the set includes the signature FXF motif of the ZP-C domain, as well as the invariant Arg of the ZP-N fg loop and its counterpart acidic residues on ZP-C. The interactions mediated by these elements are not only essential for UMOD polymerization^21^, but also required for the homodimerization and secretion of the cZP3 precursor^6^. This suggests how, following EHP release and the stochastic dissociation of one of their two ZP-C/ZP-N interfaces, both subunits of a ZP3 homodimer may be incorporated head-to-tail into a growing ZP3 protofilament.

Outside the six-residue interface core, fZP1 amino acids that are largely conserved in ZP1/2/4 homologs form a complex network of hydrogen bonds, salt bridges and hydrophobic interactions. These include loops a’b, cc’ and c’’d of ZP-C, loop bc of ZP-N and an extensive cross-hairpin interaction involving two pairs of β-strands belonging to the ef and fg loops of ZP-C and ZP-N, respectively (Figure 6C, E). These contacts make the fZP1-C/fZP1-N interface structurally distinct and significantly larger (880 vs 627 Å^2^) than fZP3-C/fZP3-N. Together with differences in the latter at the level of the core (where the invariant Asp is supplied by ZP3-specific 3_10_ helix c’’d), the ef/fg loop interaction (where a much shorter ZP-N ef loop can only contribute a minimal β-strand) and the C-terminal tail of fZP3-C (where a stretch of 11 residues interacts with the subsequent subunit within the ZP3 protofilament) (Figure 6B, F, G), these features explain the homomeric nature of ZP-C/ZP-N interactions.

The heteromeric ZP-N/ZP-C interfaces are smaller than homomeric ones (fZP1-N/fZP3-C: 494 Å; fZP3-N/fZP1-C: 571 Å) and less polar, as reflected by a distinct common core of five conserved hydrophobic residues (Figure 6H-J). These include the signature Tyr within the ZP-N domain E’-F-G extension^32^ (fZP1 Y339, fZP3 Y149), hydrogen-bonded to the main chain of ZP3 V245 or the side chain of invariant ZP1 D451, and a second near-invariant Tyr at its C-terminus (fZP1 Y371, fZP3 Y179), involved in highly conserved intermolecular hydrogen/cation-π and stacking interactions with fZP3 H244, L207 or fZP1 R449, L398 (Figure 6K, L). Mutation of the signature Tyr in human inner ear protein α-tectorin disrupts the tectorial membrane and causes non-syndromic deafness^33^; moreover, both Tyr are implicated in UMOD homopolymerization^21^ and replaced by non-hydrophobic amino acids in endoglin, a non-polymeric ZP module-containing protein^32^. Overall, the heteromeric interfaces appear to be less discriminating than the homomeric ones; however, some interactions outside their conserved core clearly favor the observed subunit combinations. These include contacts made by near-invariant fZP3 R181 (substituted by A373 in fZP1), as well as hydrophobic interactions between fZP3 M142 (E332 in fZP1) and the loop stapled by the conserved C_a_-C_b_ disulfide of fZP1-C (Figure 6K, L).

Elements outside ZP-N and ZP-C also contribute to the specificity of subunit interaction. Rationalizing previous biochemical studies of the bovine (b) ZP3/ZP4 complex^34^, the near-invariant WVPF motif at the center of the fZP3 IDL engages in yet another cation-π interaction with fZP1-C (fZP3 W194/fZP1 R528, corresponding to bZP3 W156/bZP4 K434) and hydrophobic interactions between its Phe (fZP3 F197, bZP3 F159) and a large patch of fZP1-N residues highly conserved in ZP1/4 homologs, including bZP4 (Figure 6M). Interestingly, in animal classes other than fish (where sperm penetrates the egg coat through the micropyle and VE proteins have only a structural role), the residue that immediately precedes the WVPF motif is almost always a Thr. O-glycosylation at this position has been implicated in gamete recognition in bird and mammals^6,35^ and the Thr is closely preceded by the only highly conserved N-glycosylation site of ZP3 (N184 in fZP3), important for sperm binding in bovine^34^. The map shows clear density for a corresponding N-glycan that contributes to the ZP3/ZP1 interface by packing against the βC-βC’ region of fZP1 ZP-C, including the invariant Trp (W462) that immediately follows conserved Cys 5 of ZP1/2/4 (Figures 6N and S6D). Substitution of the Trp with Val in ZP3 homologues and additional differences, such as fZP1 S536 being replaced by bulky H320 in fZP3, indicate that a comparable interaction between the ZP3 N-glycan and another ZP3 subunit would be less favored (Figure 6O). Notably, the N184 glycan can be enzymatically cleaved from the fZP3 precursor and is not required for its secretion (Figure 6P), whereas — consistent with its interaction with fZP1 — it is protected from digestion in the context of the native fZP1/fZP3 complex (Figure 6Q). These observations suggest that the conserved N-glycan of ZP3 contributes to egg coat filament architecture. Moreover, the packing of crystal form II (Figure S6H) hints at the possibility that, in the case of the compact VE of fish, the same glycan also facilitates the close juxtaposition of filaments observed by EM^21^.

### AlphaFold-based modeling of the human ZP

Although AlphaFold models of monomeric proteins can generally facilitate crystallographic structure determination by MR^36^, this approach did not succeed in the case of the fZP3/fZP1 complex due to the major conformational changes involved in ZP subunit polymerization (Figures 5 and 6). However, the fact that we could use a multi-chain prediction to successfully phase the challenging fZP3/fZP1 X-ray diffraction data indicates that the multimer version of the neural network could accurately predict the structure of the corresponding ZP module complex. Based on this consideration, we extended our experimental studies by generating a comprehensive prediction matrix for possible human ZP subunit interactions (Figure S7). This computational analysis was first carried out for three-component systems consisting of one ZP module and two ZP-C and ZP-N moieties (based on the fZP3/fZP1 complex), and subsequently extended to systems encompassing four complete ZP modules. Because AlphaFold has no explicit knowledge of the concept of filament, top-scoring models from the latter set of predictions consisted of either circular or linear, but artificially stretched, arrangements of subunits. Nonetheless, the network’s predictions consistently replicated the basic features of the experimentally observed filament architecture whenever copies of the ZP3 ZP module were combined with either partial or full ZP modules from ZP1, ZP2 or ZP4, or vice versa. By integrating this information with the crystallographic knowledge on fZP3/fZP1 (Figures 5 and 6), the structural data on ZP2 NTR (Figures 1, 2 and 4) and previous investigations of the ZP filament cross-link^26^, we were able to assemble molecular models of human ZP filaments in their pre- and post-fertilization state (Figure 7A-C). As discussed in the following sections, this complemented our empirical findings to yield major insights into egg coat architecture, suggest how ZP2 cleavage blocks polyspermy at the molecular level, and rationalize *ZP* gene mutations linked to human infertility.

**Figure 7.**
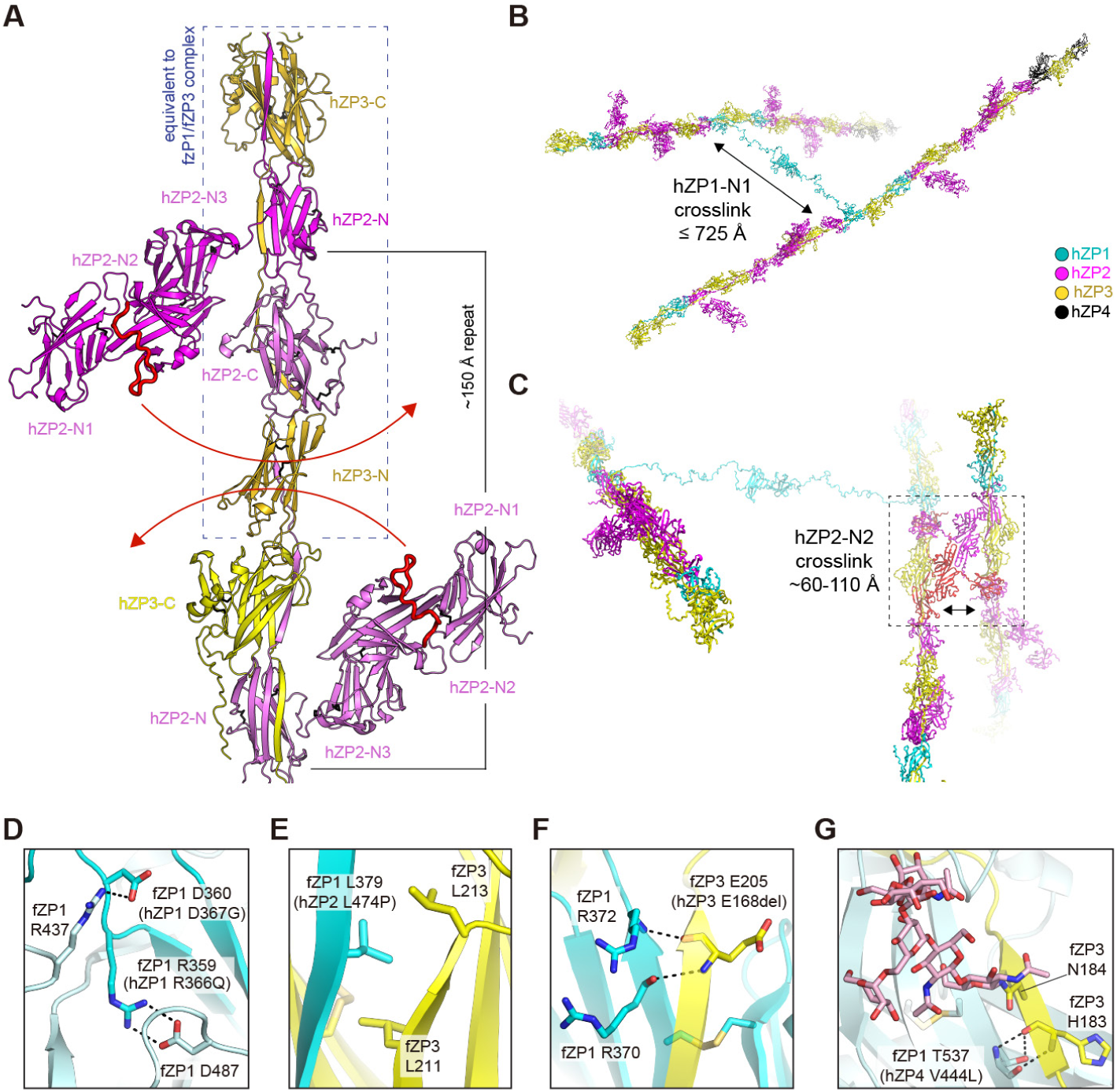
Modeling of human ZP filaments, before and after post-fertilization cleavage of ZP2. (A) Section of a hZP filament model encompassing two complete hZP3 and hZP2 subunits (gold and pink, respectively), as well as the NTR+ZP-N of another ZP2 subunit (magenta) and the ZP-C of another ZP3 subunit (yellow) plus part of the respective IDLs. The red arrows show that, although ZP2 NTR is predicted to adopt variable orientations relative to the core of the filament (Figure S7D), it cannot form an intramolecular cross-link with an adjacent copy of ZP2 upon post-fertilization cleavage of its ZP-N2 bc loop (red) by ovastacin. (B) Before fertilization, hZP filaments are only loosely cross-linked via covalent homodimerization of the ZP-N1 domain of hZP1, which is connected to the filament core by a long flexible linker. (C) Post-fertilization cleavage triggers the homodimerization of hZP2 NTRs that protrude from different filaments, non-covalently cross-linking the ZP into a tight mesh. (D-G) Close-up views of residues corresponding to positions affected by human *ZP* gene mutations associated with infertility. See also Figure S7.

## DISCUSSION

### A conserved framework for egg coat assembly

Underlining the conservation of the ZP module as the building block of egg coats from mollusk to human^14^, the 158 Å helical rise of the fish VE filament (Figure 5B, C) is remarkably consistent with the structural repeat of ∼150 Å suggested by early EM studies of the mouse ZP^16,37,38^. However, due to the unique architecture of the ZP module polymer, this conserved repeat does not correspond to a subunit heterodimer (as previously suggested for mZP2 and mZP3), but rather to a complete ZP module from one protofilament embracing two half-ZP modules from the other (Figure 5A-C). A key feature of the repeat is the IDL of the ZP module, whose contact area with the interacting ZP-C/ZP-N unit is more than double the ZP-C/ZP-N interface itself. Because of this, the IDL acts as a molecular tape that strongly stabilizes the interaction between adjacent domains within the opposite protofilament (Figure 6M-O). Combined with the fact that the ZP module of ZP1 is an integral component of the egg coat filament (as opposed to being peripherally attached, as previously hypothesized), this explains how the fish hatching enzyme disrupts VE filaments by targeting an exposed site in the middle of fZP1 IDL^29^ (Figure 5B, C).

In the case of the crystallized fish VE complex, the presence of two protofilaments is a direct product of the homomeric ZP-C/ZP-N interfaces that mediate the head-to-tail interaction between fZP3 or fZP1 subunits, respectively (Figures 5 and 6). However, AlphaFold modeling suggests that the same two-protofilament architecture is conserved across vertebrates, regardless of how many different subunits make up the respective VEs/ZPs. This is because, as a result of their ability to interact head-to-tail with each other, ZP1, ZP2 and ZP4 can coassemble into a common protofilament, which pairs with a ZP3 protofilament like the two sides of a zipper (Figure 7). Such an organization is consistent with the notion that — based on structural differences in their ZP-C domains — egg coat proteins can be divided into type I (ZP3) and type II (ZP1/2/4) subunits (with the latter type also including non-mammalian ZPAX and ZPY components)^6,20,39,40^ and the observation that all vertebrate egg coats contain at least one subunit of each type^1,41^. Moreover, it agrees with the relative ∼1:4:5 stoichiometry of mouse ZP1-3 subunits^38^. Most importantly, because the protein-protein interactions within one protofilament are cemented by the IDLs of the other (Figure 5B, C), the filament architecture straightforwardly explains the results of *ZP* gene ablation/replacement experiments in mouse, rat, rabbit and medaka^23,28, 42–49^. For example, a mouse egg coat filament model where a type I mZP3 protofilament pairs with a type II protofilament containing 80% mZP2 and 20% mZP1 rationalizes why the ZP is completely absent and approximately half the normal width in *Zp3^-/-^* and *Zp3^+/-^* mice, respectively^42,43,50^; why the ZP is clearly present in *Zp1^-/-^* animals, despite being abnormally organized due to lack of crosslinks^44^; and why the oocytes of *Zp2^-/-^* mice have a transient, very thin ZP made up of mZP3 and mZP1^28^. At the same time, the existence of type I and type II protofilaments is compatible with the observation that, when incubated *in vitro* at relatively high concentrations, individual subunits purified from fish or mouse egg coats can form homooligomers^51,52^. Finally, the apparent interchangeability of type II subunits suggests that their incorporation into growing egg coat filaments is not only stochastic, but also reflects their relative localization and expression levels; this explains how restricted areas of the egg coat, such as the germinal disc-facing part of the avian VE^53^, may in some cases become specialized to present a specific population of polymeric subunits implicated in sperm binding.

### Mechanism of the ZP2 block to polyspermy

The structural organization of the egg coat filament implies that, even when two copies of ZP2 immediately follow each other within a type II protofilament, their ZP-N2 domains are too far to interact between themselves (Figure 7A). Considering that ZP filaments are relatively straight and largely parallel to adjacent ones within ZP bundles^54^, this strongly suggests that post-fertilization cleavage triggers interaction between the processed NTRs of ZP2 subunits belonging to different filaments. Based on the relative abundance of ZP2^3^ and our structural information (Figure 2J, K), this will extensively bridge ZP filaments and bring them within less than 110 Å from each other, approximately 7-fold closer than the maximum distance allowed by their intermolecular ZP1 cross-links^26,37^ (Figure 7B, C). Consistent with the phenomena associated with hardening^55^ and the phenotype of the ZP2 ΔN1/ΔN1 mice (Figure 4), this will in turn rigidify the overall structure of the matrix and prevent polyspermy by hindering sperm penetration. This mechanism provides a molecular basis for the observed coalescence of hZP filaments after fertilization^54^, as well as the polyspermy associated with a female infertility mutation in the hOvst gene^56^. Moreover, it conceptually resembles how the fish VE is hardened by transglutaminase-mediated cross-linking of its filaments (which lack ZP2)^57^, and is compatible with the recent suggestion that cross-linking of ZP3 may also contribute to the block to polyspermy in the mouse^58^. On the other hand, in line with the observation that ZP2 cleavage is evolutionary conserved^4,59–61^ whereas ZP2-N1 is post-translationally removed from amphibian ZP2 before egg-sperm interaction and fertilization^4,62^, the mechanism does not support the currently accepted model whereby ZP2 cleavage abolishes gamete interaction through destruction or masking of a sperm-binding site on ZP2-N1^23,63^. Instead, our findings reformulate the supramolecular model of fertilization^10,61^ by suggesting that the key role of ZP2 processing — which in mammals can be crucially controlled by ZP2-N1 itself (Figures 3 and 4) — is to alter the ZP architecture so that it physically prevents the penetration of sperm. Notably, as implied by the phenotype of the ZP2 ΔN1/ΔN1 mice (Figure 4), this mechanism is compatible with data suggesting that, depending on the species, one or more binding sites other than ZP2-N1 may significantly contribute the initial attachment of sperm to the egg coat^4,6,34^. At the same time, again consistent with the evolutionary conservation of ZP2 cleavage, the mechanism does not intrinsically depend on the nature of such interaction(s), or how gamete recognition is abolished following fusion. The basis of the ZP2 block to polyspermy is thus reminiscent of the extracellular matrix changes involved in the formation of the fertilization envelope of invertebrates^64^ or the cell wall block to polyspermy of plants^65^.

### Significance for human infertility

Because of the dramatic conformational changes that the ZP module undergoes during polymerization (Figure 5D), the previously available structures of isolated ZP subunits or domains thereof^6,8,20^ could give only limited information about their mature filamentous form. The availability of detailed structural knowledge on the egg coat filament not only provides a solid molecular framework for informing future studies of sperm-ZP interaction, but also significantly improves our ability to interpret the growing number of human *ZP* gene mutations linked with female infertility^66^.

For example, hZP1 R366Q, associated with an oocyte maturation defect that leads to primary infertility due to atresia or empty follicles^67,68^, affects the key Arg residue engaged in the invariant Asp/Arg salt bridge of homotypic interfaces (fZP1-C D487/fZP1-N R359; Figures 6E and 7D). This suggests that the mutation impairs the assembly of human ZP type II protofilaments, by disrupting the interaction between hZP1 R366 and the invariant ZP-C aspartic acid of hZP1 (D483), hZP2 (D571) or hZP4 (D393). Notably, mutation of the near-invariant Asp residue that immediately follows R366 has also been reported to cause infertility, when in compound heterozygosis with a frameshift mutation that produces a secretion-impaired truncated hZP1 variant^69^. The structure of the egg coat filament rationalizes the lack of a ZP associated with the hZP1 D367G mutation by showing that the corresponding fZP1-N residue (D360) makes an intersubunit hydrogen bond with another highly conserved residue, fZP1-C R437 (hZP1 R433) (Figures 6C, E and 7D). This does not lend support to a previous suggestion that the mutation affects the ability of D367 to interact intramolecularly with non-conserved residues R299, H306 and S368^69^.

On the other hand, the suggestion that hZP2 mutation L474P causes ZP abnormality and infertility not only by reducing protein secretion but also by affecting hZP2/hZP3 interaction^70^ is supported by the insertion of the corresponding fZP1 IDL residue (L379) between two conserved Leu residues of ZP3 ZP-C (L211 and L213 in fZP3; Figure 7E). Similar considerations apply to hZP3 E168del, an IDL residue deletion found in patients with empty follicle syndrome^71^ (Figure 7F), as well as hZP4 V444L, a mutation that is associated with thin and irregular ZPs^72^ and affects a ZP-C residue (corresponding to fZP1 T537) that interfaces with the ZP3 IDL amino acid preceding the conserved N-glycosylation site (Figure 7G).

### Conclusions

Considering the phylogenetic relationship and domain architecture similarity between ZP4 and ZP1 and between ZPAX/ZPY and ZP2^12,41^, the present data provides comprehensive structural knowledge on the vertebrate egg coat. By revealing how ZP proteins heteropolymerize into a filamentous matrix that is remodeled upon cleavage of ZP2 NTR, our studies also show an example of how the mosaic architecture of ZP module-containing proteins allows them to orchestrate a complex supramolecular rearrangement in response to a defined molecular signal. Combined with the knowledge on UMOD homopolymerization, these findings open the way to a general mechanistic understanding of how other members of the ZP module superfamily evolved to carry out their remarkably diverse biological functions.

## STAR★METHODS

### KEY RESOURCE TABLE

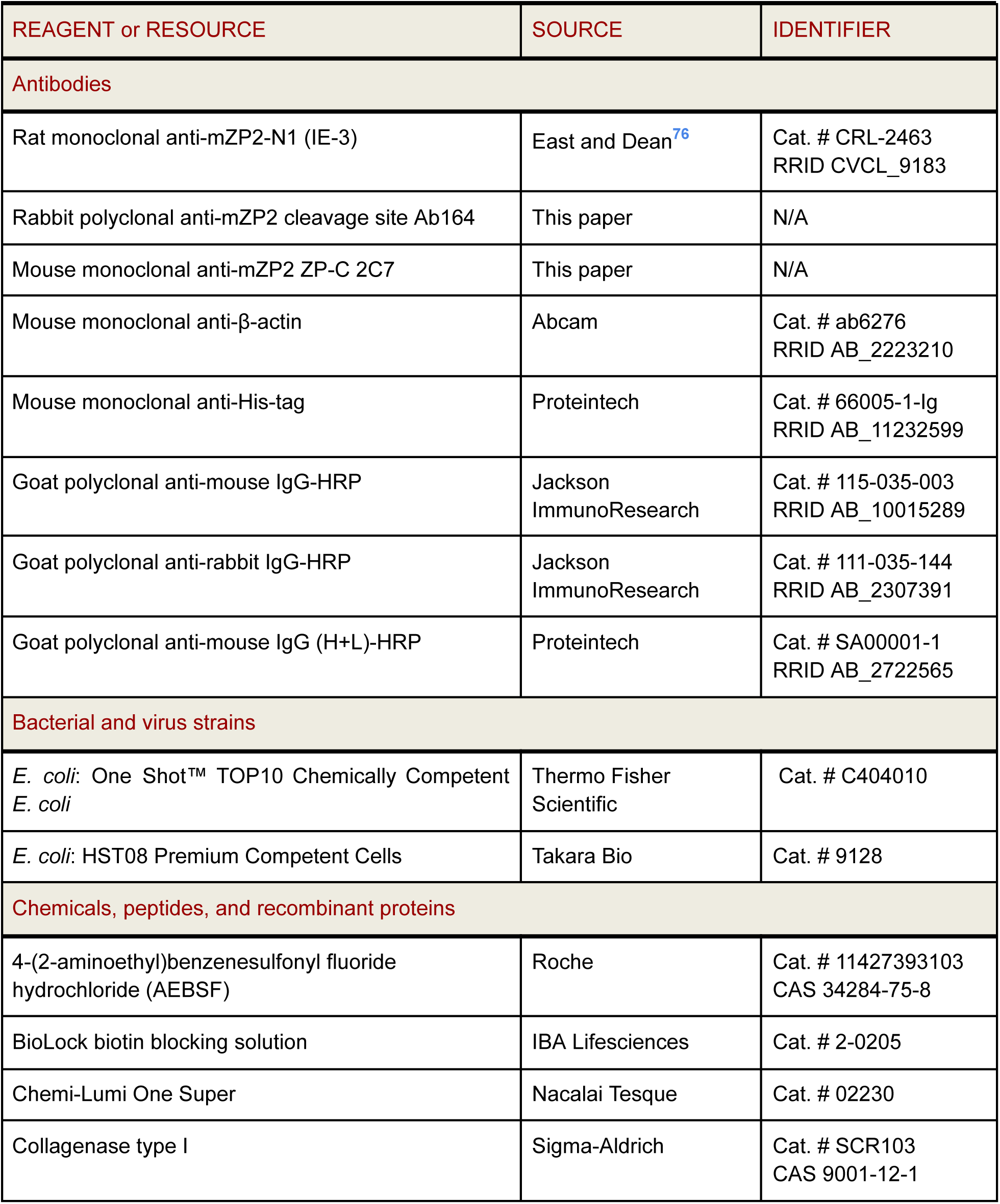
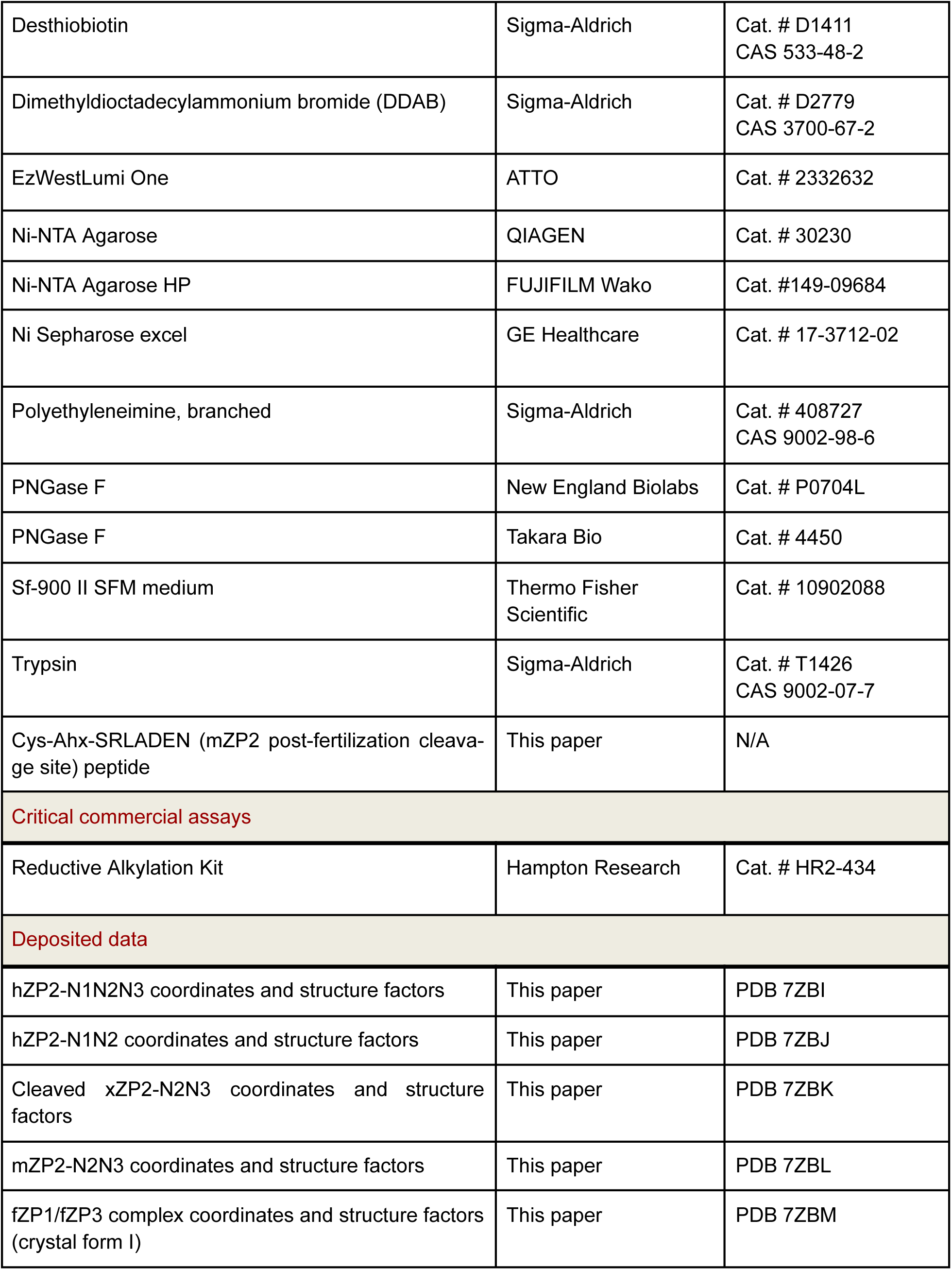
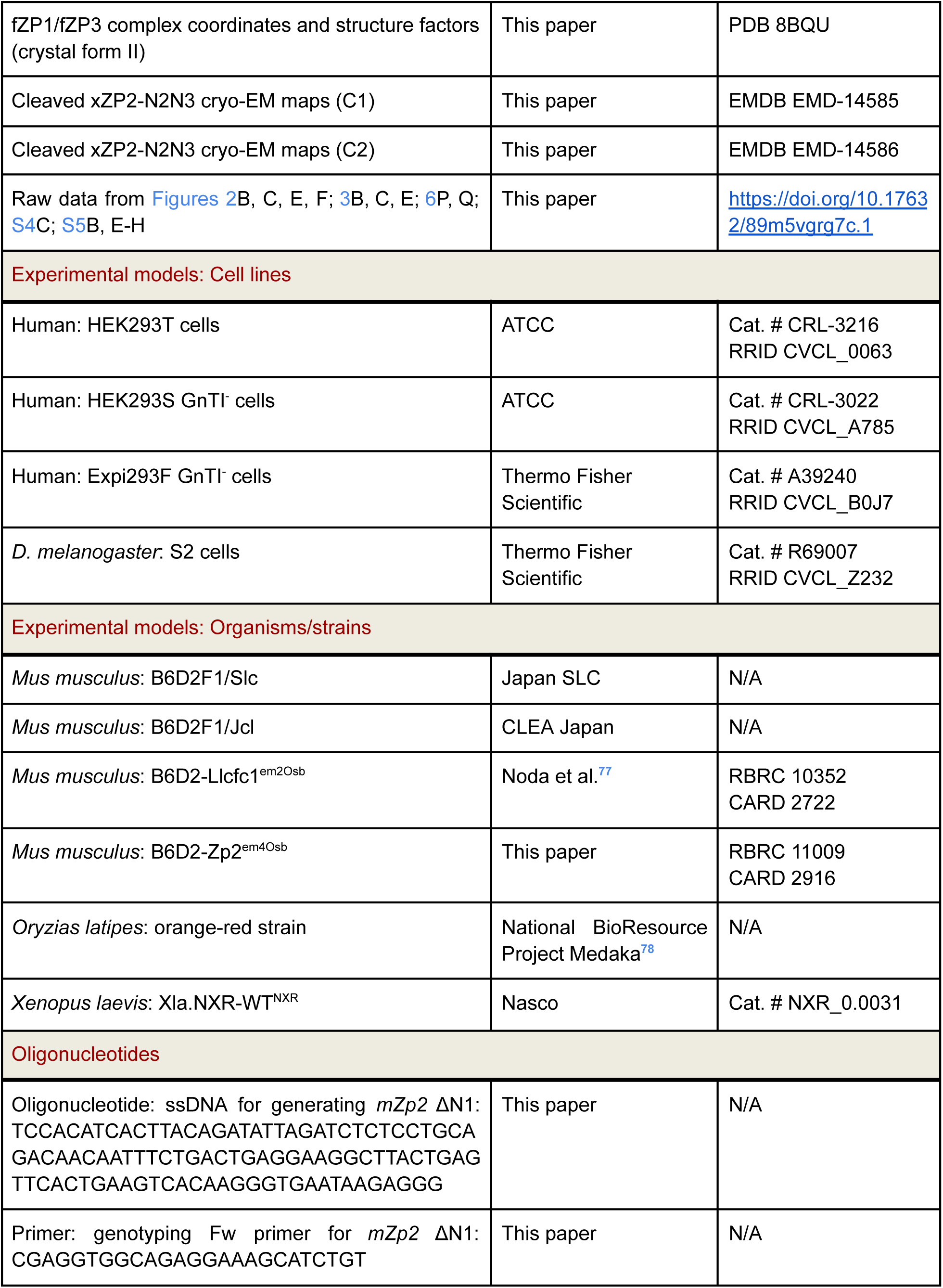
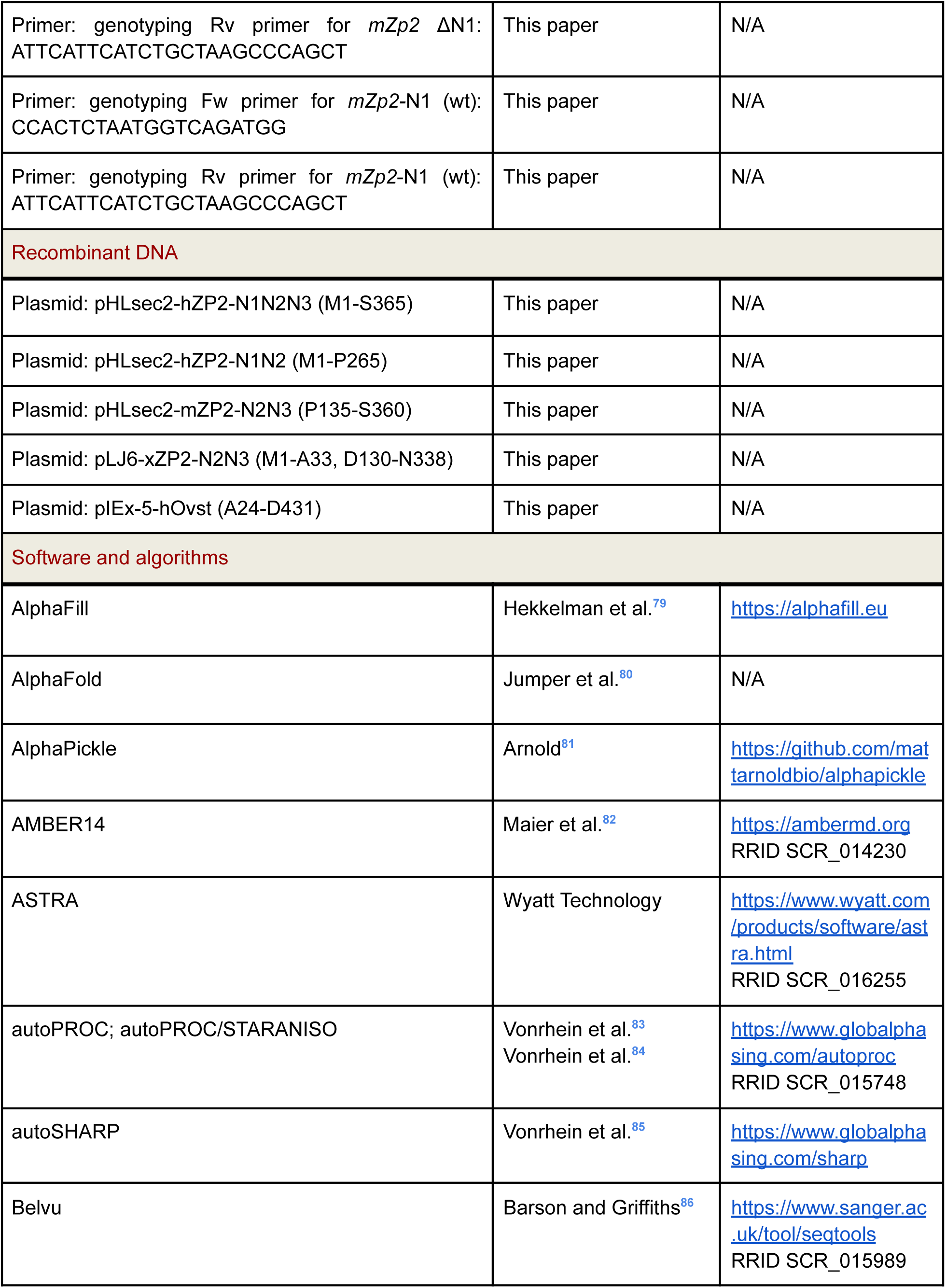
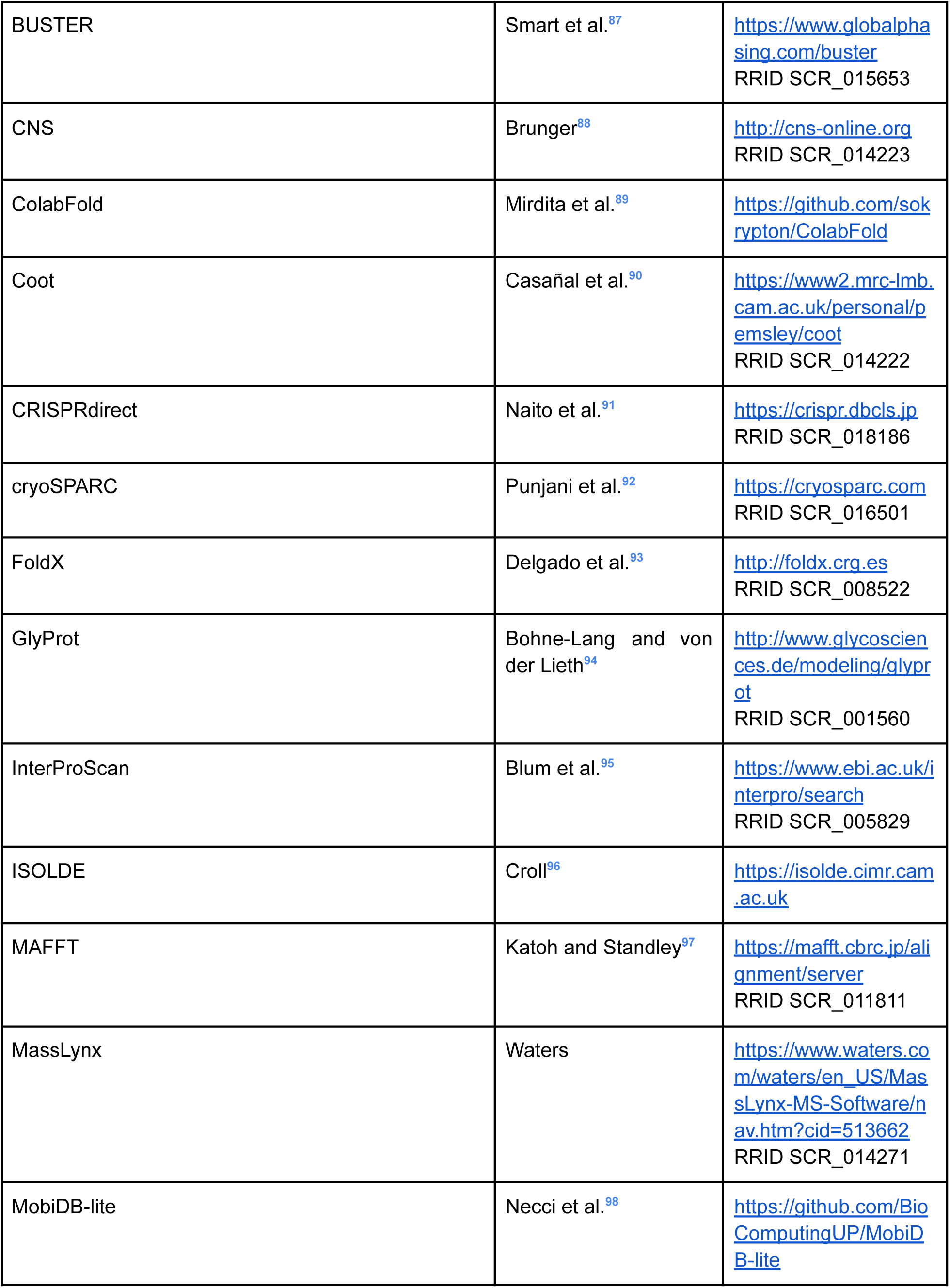
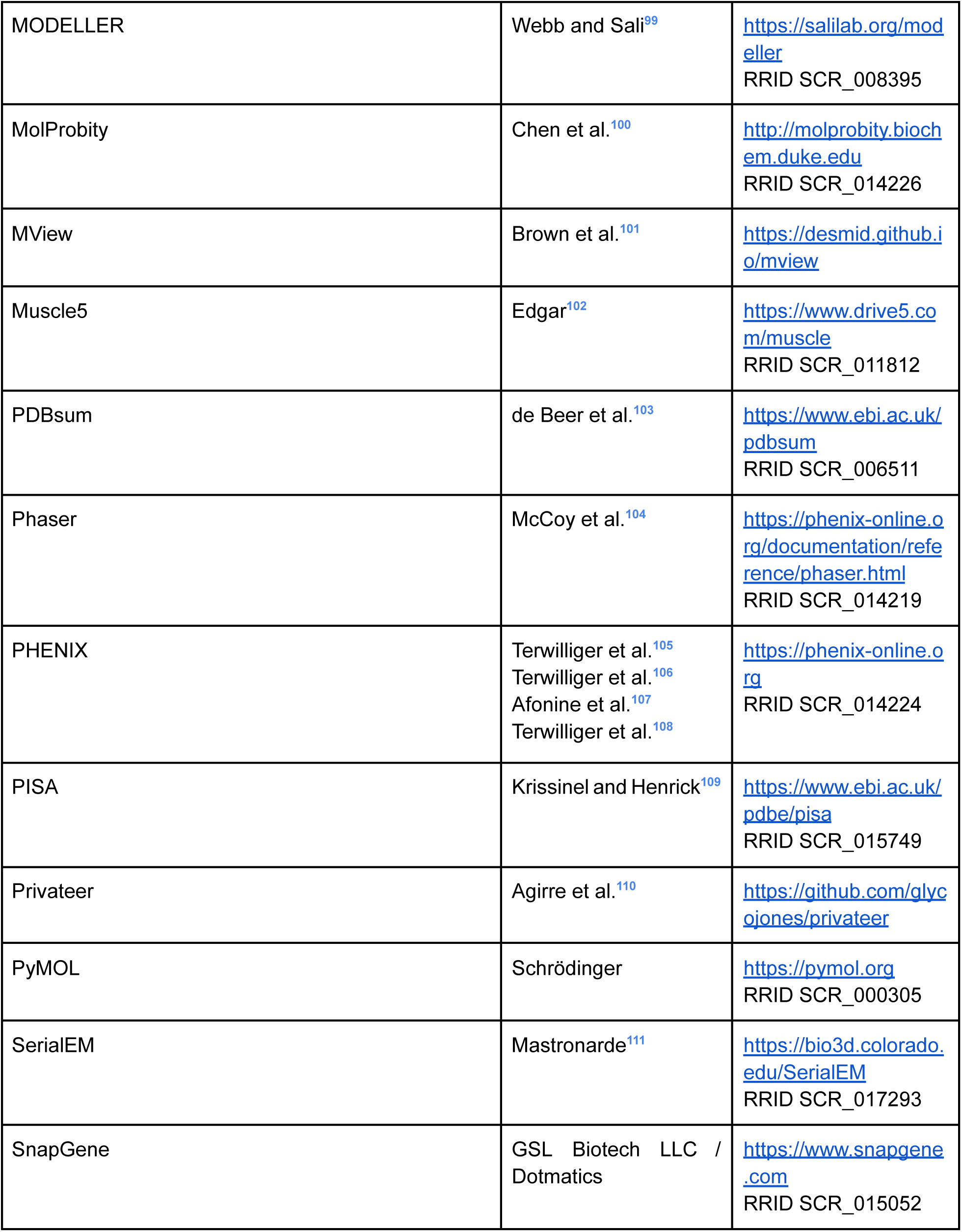
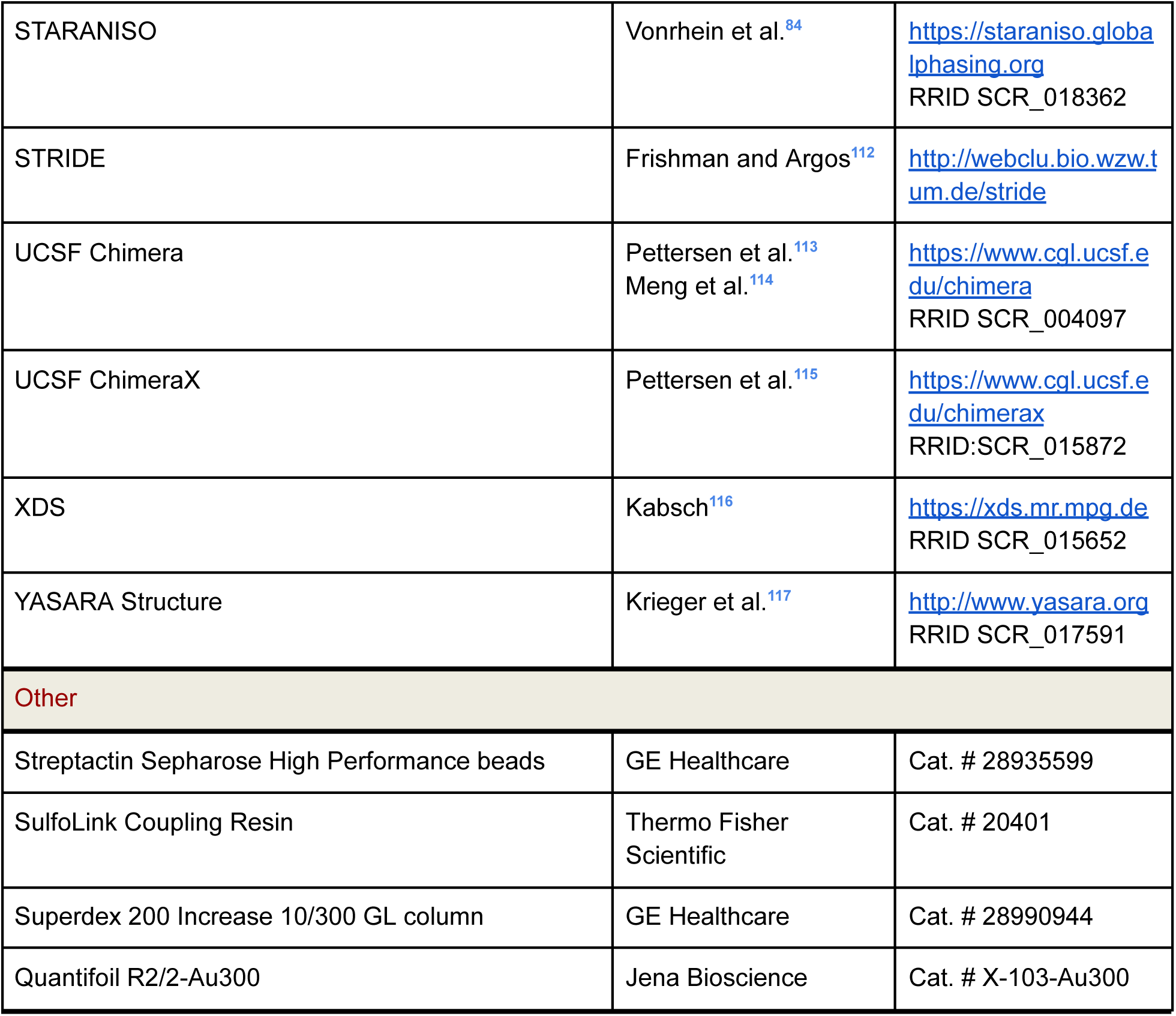

### RESOURCE AVAILABILITY

#### Lead contact

Further information and requests for resources and reagents should be directed to and will be fulfilled by the lead contact, Luca Jovine.

#### Materials availability

All unique/stable reagents generated in this study are available from the lead contact.

#### Data and code availability

- The structural models and density maps have been deposited at the Protein Data Bank and are publicly available as of the date of publication. Accession numbers are listed in the key resources table. The Cryo-EM maps have been deposited at the Electron Microscopy Data Bank and are publicly available as of the date of publication. Accession numbers are listed in the key resources table. Raw data from Figures 2B, C, E, F; 3B, C, E; 6P, Q; S4C; S5B, E-H were deposited on Mendeley at https://doi.org/10.17632/89m5vgrg7c.1.
- This paper does not report original code.
- Any additional information required to reanalyze the data reported in this paper is available from the lead contact upon request.

### EXPERIMENTAL MODEL AND SUBJECT DETAILS

*Escherichia coli* cells were cultured at 37°C in 2xYT medium containing 50 µg/mL ampicillin or 12.5 µg/mL kanamycin.

*Xenopus laevis* females were housed at 18°C with a 12 h/12 h light/dark cycle, following procedures conducted using accepted standards of humane animal care and approved by the Animal Care and Use Committee at the University of Pittsburgh (protocol 20026806).

All mice were housed with a 12 h/12 h light/dark cycle and food/water available *ad libitum*. All experiments involving mice were approved by the Institutional Animal Care and Use Committees of Research Institute for Microbial Diseases of Osaka University (approval ID Biken-AP-H30-01) and were conducted in compliance with the guidelines and regulations for animal experimentation. For mouse fertility test, sexually mature ZP2ΔN1 female mice (5-16-week-old) and wt B6D2F1 mice were caged with 8-week-old wt male mice for 2 months.

HEK293T cells (a kind gift of R. Aricescu and Y. Zhao, University of Oxford) and HEK293S GnTI- cells (ATCC) were cultivated at 37°C, 5% CO_2_ in DMEM (Thermo Fisher Scientific) supplemented with 10% FBS (Biological Industries) and 4 mM L-glutamine (Thermo Fisher Scientific). Expi293F GnTI- cells (Thermo Fisher Scientific) were grown at 37°C, 8% CO_2_ in FreeStyle™ 293 Expression Medium (Thermo Fisher Scientific). *Drosophila melanogaster* S2 cells (Thermo Fisher Scientific) were maintained at 27°C and ambient CO_2_ in Sf-900 II SFM medium (Thermo Fisher Scientific).

### METHOD DETAILS

#### DNA constructs

cDNAs encoding hZP2-N1N2N3 (M1-S365) and hZP2-N1N2 (M1-P265) were amplified by PCR from I.M.A.G.E. Consortium clone 40007473 (ATCC) and subcloned into mammalian expression vector pHLsec2, in frame with a C-terminal Leu+Glu+6His (LEH_6_)-tag-encoding sequence. A cDNA fragment encoding mZP2-N2N3 (P135-S360) cDNA, amplified from a full-length *Zp2* cDNA^118^, was subcloned into the same vector, in frame with 5’ and 3’ sequences encoding an N-terminal chicken Crypα signal peptide and the C-terminal LEH_6_-tag, respectively.

Synthetic genes encoding the ectodomain (D130-D650), N1N2N3 (M34-N338), N2N3 (D130-N338) and N2N3ZP-N (D130-G447) regions of xZP2 (ATUM) were subcloned into mammalian expression vector pLJ6, in frame with sequences that encoded the native xZP2 signal peptide (M1-A33) and a C-terminal 8His-tag.

A cDNA fragment encoding hOvst without signal peptide (A24-D431) was amplified from ASTL(NM_001002036) cDNA (GenScript) and subcloned into insect cell expression vector pIEx-5, in frame with 5’ and 3’ sequences encoding an AKH signal peptide and a C-terminal Strep-tag II (PGWSHPQFEK), respectively.

For evaluating the contribution of the conserved ZP3 N-glycan to protein secretion, synthetic genes encoding full-length wild-type fZP3 and its N184A counterpart (ATUM) were cloned into pLJ6, in frame with the sequence encoding the C-terminal LEH_8_ tag.

Oligonucleotides were purchased from Sigma-Aldrich and all constructs were verified by DNA sequencing (Eurofins Genomics).

#### Protein expression and purification

For expression construct evaluation, biochemical studies and cryo-EM, proteins were expressed in HEK293T cells (ATCC) transiently transfected with 25 kDa branched polyethyleneimine (PEI; Sigma-Aldrich)^119,120^. The same method was used to transfect HEK293S GnTI^-^ cells (ATCC) in order to produce hZP2-N1N2N3, hZP2-N1N2 and mZP2-N2N3 proteins for X-ray crystallography. After capture from the conditioned media of transfected cells by immobilized metal affinity chromatography (IMAC) with Ni-NTA Agarose (QIAGEN) or Ni Sepharose excel (GE Healthcare) beads, proteins were subjected to size-exclusion chromatography (SEC), concentrated and stored at -80°C until use. For crystallographic studies, a deglycosylation step with Endoglycosidase H (Endo H) was also included before SEC. Reductive methylation of hZP2-N1N2N3 was carried out using a Reductive Alkylation Kit (Hampton Research); production of SeMet-hZP2-N1N2 was performed following an established protocol^119^.

xZP2-N2N3 was expressed in Expi293F GnTI^-^ cells as described^121,122^, and purified and deglycosylated as detailed above.

For expression of hOvst, *Drosophila* Schneider 2 (S2) cells were cultured in Sf-900 II SFM medium (Thermo Fisher Scientific) and transiently transfected using dimethyldioctadecylammonium bromide (DDAB; Sigma-Aldrich)^123^. 90-96 h after transfection, the conditioned medium was harvested and cell debris was removed by centrifugation at 4,000 x g. The supernatant was adjusted to 0.1 M Tris-HCl pH 8.1, 150 mM NaCl (buffer A) and 240 µL BioLock (IBA Lifesciences) was added to 100 mL medium. After 20 min incubation at room temperature, the medium was further centrifuged at 4,000 x g for 15 min. The cleared medium was incubated with 1/100 volume of Streptactin Sepharose High Performance beads (GE Healthcare) pre-equilibrated with buffer A for 1 h at room temperature. Streptactin beads were collected to an open column by gravity flow and washed with 20 volumes buffer A. Bound protein was batch-eluted 5 times with 1 mL buffer A supplemented with 5 mM desthiobiotin. Fractions containing hOvst were concentrated with 10 kDa-cutoff centrifugal filtration devices and stored at -80°C until use.

The native fZP1/fZP3 complex, corresponding to fraction F2 of the medaka VE hatching enzyme digest, was prepared from unfertilized eggs isolated from spawning female Japanese rice fish, essentially as described^21^. All procedures were approved by the Ethics Committee of Sophia University (approval ID 2016-006).

C-terminally His-tagged recombinant fZP3 and its N184 mutant counterpart were expressed in HEK293T cells transiently transfected by 25 kDa branched PEI in 6-well plates. 72 h after transfection, conditioned media were collected and secreted proteins were captured using Ni-NTA Agarose HP (FUJIFILM Wako).

#### *In vitro* cleavage of xZP2 with collagenase

Purified xZP2-N2N3 was adjusted to 2 mg mL^-1^ in 20 mM Na-HEPES pH 7.8, 150 mM NaCl, 5 mM CaCl_2_ and incubated for 2 h at room temperature with 200 µg mL^-1^ collagenase type I (Sigma-Aldrich). The cleaved protein was then further purified by SEC, using a Superdex 200 Increase 10/300 GL column (GE Healthcare) pre-equilibrated with 20 mM Na-HEPES pH 7.8, 150 mM NaCl.

#### *In vitro* cleavage of xZP2 with *Xenopus* oocyte exudate

To collect eggs, laying was stimulated by injection of 1,000 IU human chorionic gonadotropin into the dorsal lymph sac of sexually mature *X. laevis* females obtained commercially (NASCO). Following injection, animals were housed overnight for 12-16 h at 14-16°C. Typically, egg laying began within 2 h of moving to room temperature, and batches of 12-155 eggs from 8 frogs were collected on dry petri dishes and used within 10 min of being laid. Exudate collection was performed in two steps. First, the jelly coat was removed by agitating the eggs in 45 mM β-mercaptoethanol in dilute modified Ringer’s solution (33.3 mM NaCl, 0.6 mM KCl, 0.67 mM CaCl_2_, 0.33 mM MgCl_2_, 1.33 mM Na-HEPES) pH 8.5 for 2-3 min; once de-jellying was confirmed by visual observation, eggs were neutralized in dilute modified Ringer’s solution pH 6.5, following by three washes in dilute modified Ringer’s pH 7.8. Second, de-jellied eggs were incubated in a 35 mm plastic petri dish containing 2 µM ionomycin in dilute modified Ringer’s solution pH 7.8 for 60 min on a flat rotating platform at 27 rpm. Following incubation, dish solution was collected into 1.7 mL Eppendorf tubes while carefully avoiding cell lysis. Samples were held on ice, spun at 6000 rcf for 30 min at 4°C using 0.5 mL Pierce protein concentrators with a polyethersulfone (PES) membrane and a 3 kDa-cutoff (Thermo Fisher Scientific), and finally transferred to clean Eppendorf tubes. After evaluating total protein concentration with a Bradford solution (1:4 dilution), samples were frozen in liquid nitrogen and stored at -80°C or on dry ice.

For *in vitro* cleavage experiments, thawed exudate samples were further concentrated 10 times using 10 kDa-cutoff centrifugal filtration devices (Amicon) and then cleared from precipitate using 0.2 µm centrifugal filtration devices (Millipore). 10 µL concentrated exudate was added to 90 µL purified xZP2-N2N3 (2 mg mL^-1^) and, after 10 days of incubation at room temperature, the sample was subjected to SEC-MALS.

#### *In vitro* cleavage of hZP2 with recombinant hOvst

For cleavage of hZP2-N1N2N3, hOvst (0.2 mg mL^-1^) was activated using 1/100 (v/v) volume of trypsin (Sigma-Aldrich) for 1 h at 37°C. Trypsin activity was quenched with 10 mM 4-(2-aminoethyl)benzenesulfonyl fluoride hydrochloride (AEBSF, Roche)^124^. hZP2-N1N2N3 (1 mg mL^-1^) was treated for 36-40 h at room temperature with trypsinized hOvst (hOvst-T) or untreated hOvst (100:1 substrate:protease ratio) and subjected to SEC-MALS. To confirm the effect of metal ions — and thus the identity of the protease responsible for cleaving hZP2 —, 5 mM EDTA was added to the reaction mixture. hZP2-N1N2N3 or hZP2-N2N3 (1 mg mL^-1^) were incubated with hOvst (100:1 substrate:protease ratio) and reaction mixture aliquots harvested at different times (0-40 h) were subjected to SDS-PAGE under reducing and non-reducing conditions to analyze the cleavage products.

#### Protein crystallization and derivatization

Crystallization trials by hanging or sitting drop vapor diffusion were set up at 20°C or 4°C using a mosquito crystallization robot (TTP Labtech).

Thick plates of methylated hZP2-N1N2N3 (6.4 mg mL^-1^) grew at 4°C in 10% (w/v) PEG 4000, 0.1 M MES/imidazole pH 6.5, 0.1 M carboxylic acids (MD2-100-76, Molecular Dimensions), 20% (v/v) glycerol. For experimental phasing, 0.1 µL drops containing crystals of the protein were mixed with 0.5 µL 15% (w/v) PEG 4000, 0.1 M MES/imidazole pH 6.5, 0.1 M carboxylic acids, 30% (v/v) glycerol, 1 mM (NH_4_)PtCl_2_ for 4 h at 4°C.

hZP2-N1N2 and SeMet-hZP2-N1N2 (5.0 mg mL^-1^) formed crystals with an hexagonal bipyramid shape at 20°C in 10-30% (v/v) MPD, 0.1 M MES pH 6.0 or 0.1 M Na-HEPES pH 7.5.

Thin plate crystals of collagenase-cleaved xZP2-N2N3 (6.9 mg mL^-1^) were obtained at 20°C by mixing 0.1 µL protein solution with 0.2 µL reservoir solution containing 12.5% (v/v) MPD, 12.5% (v/v) PEG 1000, 12.5% (w/v) PEG 3350, 0.1 M Tris base/Bicine pH 8.5, 0.09 M NPS (nitrate, phosphate, sulfate; MD2-100-72, Molecular Dimensions).

mZP2-N2N3 (7.9 mg mL^-1^) formed prismatic crystals at 20°C in 20% (w/v) PEG 3350, 0.2 M Na-nitrate.

The medaka ZP1/ZP3 complex (6 mg mL^-1^) yielded two distinct trigonal crystal forms. The first had an hexagonal plate morphology and grew at 20°C in 5% (w/v) PEG 20000, 25% (w/v) trimethylpropane, 0.1 M N,N-bis(2-hydroxyethyl)-2-amino ethanesulfonic acid (BES)/triethanolamine pH 7.5, 1% (w/v) non-detergent sulfobetaine (NDSB) 195, 0.5 mM YCl_3_, 0.5 mM ErCl_3_, 0.5 mM TbCl_3_, 0.5 mM YbCl_3_ (condition E8 of the MORPHEUS II crystallization screen^125^ (Molecular Dimensions)). The second crystal form, with a bipyramidal prismatic morphology, grew at 4°C in 22% (w/v) PEG 3350, 0.2 M sodium/potassium phosphate, 0.1 M Tris-HCl pH 8.0 (a condition obtained by optimizing an initial hit in condition 94 of the PACT suite^126^ (QIAGEN)).

#### X-ray data collection and processing

For data collection at synchrotron, single crystals were harvested using MicroMounts, MicroLoops or MicroMeshes (MiTeGen) and flash frozen in liquid nitrogen.

X-ray diffraction data was collected at Helmholtz-Zentrum Berlin (HZB)/Bessy II MX beamline BL14.1^127^ (hZP2-N1N2N3); European Synchrotron Radiation Facility (ESRF) beamlines ID23-1^128^ (hZP2-N1N2), ID29^129^ (mZP2-N2N3 and fZP1/fZP3 complex crystal form I) and ID30B^130^ (cleaved xZP2-N2N3); and Diamond Light Source beamlines I04-1^131^ (SeMet-hZP2-N1N2) and I02^131^ (fZP1/fZP3 complex crystal form II).

All datasets were processed using XDS^116^, except for the fZP1/fZP3 complex crystal form II data which — as further detailed below — was processed using autoPROC/STARANISO^84^ (Tables S1 and S3).

#### X-ray structure determination of hZP2-N1N2N3

Initial attempts to solve the structure of hZP2-N1N2N3 by MR using search models derived from the crystal structure of mZP2-N1^9^ failed. Data collected from Pt-soaked crystals could however be phased by single isomorphous replacement with anomalous scattering using autoSHARP^85^. This identified 8 Pt sites that resulted in a phasing power for isomorphous differences of 0.688/0.759 (acentric/centric reflections) and a phasing power for anomalous differences of 0.524. Despite the very low figure of merit (FOM) of the initial phases (0.087/0.108), density modification^132^ showed a clear difference between the two hands that had scores of 0.1576 and 0.0855, respectively; moreover, alternated cycles of density modification and automated model building^133,134^ produced an initial model (R_free_ 0.51) that showed some evidence for the presence of β-sheets. When this set of coordinates was used as a search model for MR with Phaser^104^, it produced a single solution (LLG 468, TFZ 18.7) that could be improved to R_free_ 0.40 by PHENIX AutoBuild^105^. The structure, whose asymmetric unit consists of three molecules arranged in ring-like fashion, was completed by alternating cycles of manual rebuilding in Coot^90^, refinement with non-crystallographic symmetry (NCS) restraints using phenix.refine^107^ and validation with MolProbity^100^ and Privateer^110^ (Table S1). Notably, chain C of the model has a significantly higher average B factor (160 Å^2^) than the other two chains (chain A: 98 Å^2^; chain B: 104 Å^2^), largely due to local disorder at the level of its ZP-N3 domain (208 Å^2^); however, because omission of the latter consistently increased R_free_ by ∼0.5%, its coordinates were kept in the model but only subjected to rigid-body refinement.

#### X-ray structure determination of hZP2-N1N2

As in the case of hZP2-N1N2N3, the structure of hZP2-N1N2 could not be phased by MR using mZP2-N1^9^ as a search model. However, MR was successful using an ensemble consisting of the latter (residues N42-Q138 of PDB ID 5II6) and the three copies of the ZP-N1 domain from a partially refined model of hZP2-N1N2N3 (residues N46-V143), trimmed using a 1.5 Å threshold. This produced a single solution with two molecules of hZP2-N1N2 per asymmetric unit (LLG 282, TFZ 18.9), whose correctness was verified by its ability to locate 9 Se sites (chain A M160, M166, M203, M239; chain B M85, M160, M166, M203, M239) by cross-Fourier analysis in PHENIX AutoSol^106^ (FOM 0.694, Bayesian correlation coefficient (BAYES-CC) 75.49), using a 3.5 Å-resolution dataset collected at 13.477 keV from crystals of SeMet-hZP2-N1N2 (Figure S2A). The structure was completed, refined and validated as described for hZP2-N1N2N3, except that refinement was carried out also using BUSTER^87^ (Table S1).

Notably, the ZP-N2 domains from mZP2-N2N3 and cleaved xZP2-N2N3 can be superimposed on the ZP-N2 domain of hZP2-N1N2N3 with average RMSDs of 2.5 Å (over 88 Cα) and 1.9 Å (over 72 Cα), respectively (Figure S2B, C). On the contrary, the ZP-N2 domain of hZP2-N1N2 shows significant differences in the orientation of the edges of its C/E’/F/G β-sheet relative to the A/B/E/D β-sheet, so that its average RMSD from the ZP-N2 domain of hZP2-N1N2N3 is 5.0 Å (over 96 Cα). Despite these differences in the hZP2-N1N2 structure, which are likely to at least partially reflect the lack of an interacting ZP-N3 domain as well as different crystal packing, the ZP-N1/ZP-N2 domain interface of hZP2-N1N2 is essentially the same as that of hZP2-N1N2N3 (Figure 1C and Figure S2D).

#### X-ray structure determination of cleaved xZP2-N2N3

The structure of cleaved xZP2-N2N3 was solved by MR, using as search model the top ranked AlphaFold prediction generated by ColabFold^80,89^ for the sequence D130-N338, trimmed to exclude two low-confidence regions corresponding to the N-terminus of mature xZP2 (D130-V137) and the cleavage loop of ZP-N2 (P149-A163). Phaser found a single unambiguous solution (LLG=427, TFZ=18.4) that consisted of 4 copies of xZP2-N2N3, corresponding to the biologically relevant tetramer, in the asymmetric unit of a P2_1_2_1_2_1_ cell. After autobuilding with PHENIX AutoBuild, several cycles of manual rebuilding and refinement could not improve R_free_ beyond ∼0.31; however, twin refinement of two copies of the xZP2-N2N3 tetramer against data reprocessed in space group P2_1_, using operator h,-k,-l, allowed structure determination to proceed to a final R_free_∼0.26 (Table S1).

#### X-ray structure determination of mZP2-N2N3

The structure of mZP2-N2N3 was solved by native single-wavelength anomalous diffraction (SAD) using a 2.0 Å-resolution dataset collected at 7 keV. PHENIX AutoSol identified a 23-atom substructure (FOM 0.232, BAYES-CC 27.69) whose phases resulted in an initial model of 299 residues (R=0.33, R_free_=0.34). After further autobuilding using PHENIX AutoBuild, which significantly improved the solution (383 residues, R=0.26, R_free_=0.29), manual rebuilding, refinement and validation were performed as detailed above (Table S1).

#### X-ray structure determination of the fZP1/fZP3 complex (crystal form I)

Initial attempts to phase the data collected from crystal form I of the native fZP1/fZP3 complex, using AlphaFold^80^ models of individual egg coat subunits or domains thereof as MR search models (either by themselves or together in the context of sequential searches), failed. However, it was possible to solve the structure by using a combination of two AlphaFold-generated subcomplex models (whose fZP1 coordinates matched the sequence of the most abundant medaka VE fZP1 isoform, derived from choriogenin H (Chg H)^135^). The first AlphaFold prediction was generated by submitting to ColabFold^89^ two sequences: the first was a hybrid sequence consisting of (a) the ZP-N domain and IDL of fZP3 (residues Y74-L207, with Y74 being the first residue of the fZP3 37 kDa low choriolytic enzyme (LCE) digestion product^29^), (b) a 10 glycine-spacer and (c) the trefoil and ZP-N domains of fZP1, plus part of the protein’s IDL (residues T221-D387, corresponding to the fZP1 17 kDa LCE digestion product^29^); the second was the sequence of fZP1 ZP-C and part of the preceding IDL (residues S388-G557, corresponding to the fZP1 16 kDa LCE digestion fragment^29^). The top 3 ranked models of this subcomplex were then aligned into an ensemble (ensemble I) and trimmed to remove low confidence regions. The second prediction was also carried out by submitting two sequences to ColabFold: the first corresponded to the IDL and ZP-C domain of fZP3 (residues H183-T393), and the second was the same fZP1 trefoil/ZP-N/IDL fragment described above (residues T221-D387). A corresponding ensemble (ensemble II) was then generated by aligning 4 of the top 5 ranked models of this subcomplex and subsequently trimming them to exclude the trefoil and ZP-N domains of ZP3 as well as remove low confidence regions.

Considering that the Matthews coefficient suggested the presence of 1 or 2 complexes per asymmetric unit and taking into account the relatively weak diffraction of the fZP3/fZP1 crystals, ensemble I was first used to search for a single molecule in the asymmetric unit using Phaser (∼79% solvent content). Using RMSD=1.5 Å as search model variance from the expected structure, this resulted in a single MR solution (LLG=95, TFZ=11.5) that was refined to R_free_=0.49. MR was then repeated with the resulting coordinates (using 100% sequence identity as search model variance), yielding the same solution but with increased LLG=203 and TFZ=17.0. Finally, a search for a single copy of ensemble II (with RMSD=1.5 Å as search model variance) was successfully carried out using the placed and refined ensemble I coordinates as a partial solution (notably, a parallel search with placed but not refined ensemble I coordinates failed). This resulted in a single solution with LLG=558 and TFZ=22.9 that was automatically refined to R_free_=0.44, producing a difference map that showed clear density for the N-glycan attached to fZP3 N184 (a post-translational modification that was not included in the search model) (Figure S6A) but no evidence for any additional copy of the complex within the asymmetric unit of the crystal. Although the fZP3/fZP1 complex data was collected at 11.563 keV, we could still take advantage of the fact that — as detailed above — the protein was crystallized in the presence of Y/Er/Tb/Yb to further confirm the correctness of the solution by its ability to locate heavy atom sites by MR-SAD in PHENIX AutoSol (FOM 0.419, overall score 75.49±3.41). Based on this result, we added to the model coordinates four sites with occupancy >0.5 (all of which also matched large positive difference density peaks), tentatively assigned as Yb by AutoSol (Figure S6A). The model was improved through iterative rounds of manual model building, taking advantage of maps generated by deformable elastic network (DEN) refinement^136^ in CNS^88^ (Figure S6B) as well as map sharpening^137^ in PHENIX^108^, and refinement and validation as detailed above (Table S3).

Notably, inspection of the map region between the ZP-C sequences FKMFTFV (identical in both the Chg H- and Chg Hminor-derived fZP1 subunits of the VE^135,138^) and PL^R^/_K_E^K^/_H_VYIHC (Chg H/Chg Hminor) showed that these are connected by a relatively short loop, consistent with the sequence of Chg H (6 intervening residues) but not that of Chg Hminor (21 intervening residues). This suggested that the crystallized VE fragments contained only the higher abundant Chg H-derived fZP1 subunit, an hypothesis that was later confirmed by the higher resolution map of crystal form II.

#### X-ray structure determination of the fZP1/fZP3 complex (crystal form II) and analysis of its packing

The second crystal form of the medaka VE filament fragment was characterized by a very long unit cell *c* axis (>470 Å) and extremely anisotropic diffraction, combined with variable diffracting power and significant anisomorphism. Although some specimens showed reflections beyond 3 Å resolution, these features precluded early attempts at structure determination by either experimental phasing or MR. The subsequent availability of a refined model for crystal form I of the complex allowed Phaser to identify a convincing single solution (LLG=222, TFZ=16.3), using a crystal form II dataset that had been conservatively processed to 3.5 Å resolution with autoPROC^83^ (Table S3, column “isotropic, standard (TRUNCATE)”). Initial refinement of the placed model, using two rigid body groups ((1) fZP3 ZP-N + IDL residues K182-L193, fZP1 IDL residues P389-L398 + ZP-C; (2) fZP1 trefoil + ZP-N + IDL residues T374-P386, fZP3 IDL residues W194-L207 + ZP-C) as well as four TLS groups (matching fZP3 ZP-N, fZP1 ZP-C, fZP1 trefoil + ZP-N and fZP3 ZP-C, plus interacting IDL moieties from the other subunits), resulted in R=0.36, R_free_=0.42. As in the case of crystal form I, the corresponding *2mFo-DFc* and *mFo-DFc* maps also confirmed the correctness of the MR solution by showing prominent density for the fZP3 N184 N-glycan, which had been omitted from the search model coordinates; moreover, the maps also showed density for fZP3 C-terminal residues G359-S374, which were disordered in crystal form I (whose packing prevented the interaction between the ZP-C and ZP-N domains of different ZP3 molecules). Consistent with visual inspection of crystal form II’s diffraction, analysis of the corresponding data with STARANISO^84^ identified an anisotropic cut-off surface that could be fit with an ellipsoid having approximate diffraction limits of 4.0 Å, 4.0 Å and 2.3 Å along its principal axes. Whereas conventional isotropic processing could not cope with such strong anisotropy beyond ∼3.0 Å resolution (Table S3, column “Isotropic, extended (AIMLESS)”), we were able to include a significant amount of higher resolution information by carrying out anisotropic processing with autoPROC^84^ (Table S3, column “Anisotropic (STARANISO)”). Refinement against anisotropically processed data lowered R_free_ by >5% and yielded overall improved maps with clear density for protein side chains (with the exception of fZP1 Q410, whose position was however constrained by well resolved adjacent residues). The model was completed by alternating cycles of rebuilding using both Coot and ISOLDE^96^, refinement with phenix.refine and validation with MolProbity and Privateer; based on visual inspection of maps calculated using different cutoffs during the course of this process, a high-resolution limit of 2.7 Å was ultimately chosen as optimal for the crystal form II dataset (Table S3).

To assess whether the packing of crystal form II of the fZP1/fZP3 complex was likely to recapitulate the conformation of intact fZP1 within the uncleaved native VE filament (and thus mirror the structure of the latter), the refined coordinates of the complex and those of its coaxially stacked -X+Y, -X, Z+⅓ + (1 0 0) & {0 0 0} and -Y, X-Y, Z+⅔ + (0 -1 -1) & {0 0 0} symmetry-related copies were imported into YASARA Structure^117^, which was then used to introduce covalent bonds between the D387 and S388 residues of adjacent fZP1 moieties. This did not introduce any clash with spatially close non-linker residues and was followed by AMBER14^82^ energy minimization of residues P385-P389 (to relax the linker region immediately around 387|388) and then of all residues within 10 Å of 387|388. Notably, this allowed S388 and P389 to make main chain hydrogen bonds with ZP3 V172 and thereby extend the β2 strand of ZP1’s IDL. Together with the formation of additional intersubunit hydrogen bonds (hydroxyl group of fZP1 S388/carbonyl oxygen of ZP3 Q170; hydroxyl group of fZP3 S169/carbonyl oxygen of fZP1 P386), this suggests that the packing of proteolytic fragments observed in crystal form II results in a supramolecular arrangement that closely resembles the structure of the intact VE filament.

#### Cryo-EM data collection and structure determination of cleaved xZP2-N2N3

Purified cleaved xZP2-N2N3 protein at 0.9 mg mL^-1^ (3 µL) was applied to cryo-EM grids (Quantifoil R2/2-Au300) previously glow-discharged under vacuum for 120 s at 20 mA (PELCO easiGLOW), incubated for 30 s, blotted for 3 s, and plunge frozen in liquid ethane using a Vitrobot Mark4 grid freezing device (FEI) with the chamber maintained at 4°C and 100% relative humidity. An optimized grid was imaged with a Thermo Scientific Talos Arctica electron microscope equipped with a K2 camera, operating at 200 kV. 1347 movies of 40 frames each were acquired in electron counting mode at 0.9459 Å/pixel, with a total dose of 52 electrons/Å^2^ and stacked into a single TIFF stack using SerialEM^111^ control software with defocus values ranging from 1.4 to 3.0 µm (Table S2).

Contrast transfer function (CTF) parameters were estimated from averaged movies using Patch CTF and beam-induced particle motion between fractions was corrected using Patch Motion within cryoSPARC^92^. Automatic particle selection was performed with templates from the initial 2D classification. The number of particle images were reduced by further 2D/3D classification and heterogeneous refinement (Figure S3). Initial maps were calculated *ab initio* and the final maps were refined using cryoSPARC’s non-uniform refinement feature^139^, with and without the application of C2 symmetry. The final raw maps have an overall nominal resolution of 4.4 Å (C2)/4.6 Å (C1) according to cryoSPARC (Table S2), or 4.9 Å (C2)/5.3 Å (C1) according to the EMDB FSC Validation Server. Rigid-body fitting of the crystallographic model of cleaved xZP2-N2N3 into the maps using ChimeraX^115^ yielded correlation values of 0.75 (C1) and 0.74 (C2), which could be improved to 0.81 (C1) and 0.82 (C2) using flexible fitting in Namdinator^140^.

#### Sequence-structure analysis

Alignments were generated using MAFFT^97^ as implemented in SnapGene (GSL Biotech), MUSCLE5^102^ or UCSF Chimera^113^ MatchMaker/Match→Align^114^; edited and analyzed with Belvu from the SeqTools package^86^; and rendered using ESPript^141^ or MView^101^. Disorder predictions were carried out using MobiDB-lite^98^ and AlphaFold^142^. Maps and models were visualized and analyzed using Coot, PyMOL (Schrödinger) and UCSF Chimera/ChimeraX. Secondary structure was assigned using STRIDE^112^. Interfaces with analyzed with PISA^109^, PDBsum^103^ and FoldX^93^. Electrostatic surface potential calculations were carried out using the APBS Tools plugin of PyMOL^143^. Structural mapping of the evolutionary conservation of amino acid positions was performed with ConSurf^144^, using manually curated sequence alignments derived from NCBI RefSeq^145^. Structural figure panels were generated using PyMOL, except for those of Figure S3 which were created with UCSF ChimeraX. AlphaFold Predicted Alignment Error (PAE) plots were generated with AlphaPickle^81^.

#### Size exclusion chromatography-multi angle light scattering (SEC-MALS)

Uncleaved or cleaved xZP2-N2N3 or hZP2-N1N2N3 (100-150 µg) were measured using an Ettan LC high-performance liquid chromatography system with a UV-900 detector (Amersham Pharmacia Biotech; λ = 280 nm), coupled with a miniDawn Treos MALS detector (Wyatt Technology; λ = 658 nm) and an Optilab T-rEX dRI detector (Wyatt Technology; λ = 660 nm). Separation was performed at 20°C using a Superdex 200 Increase 10/300 GL column (GE Healthcare), with a flow rate of 0.5 mL min^−1^ and a mobile phase consisting of 20 mM Na-HEPES pH 7.8, 150 mM NaCl. Data processing and weight-averaged molecular mass calculation were performed using the ASTRA software (Wyatt Technology). The percentages of tetrameric and dimeric species reported in Figure 2A, D were calculated using the formula (4xMW_monomer_)X+ (2xMW_monomer_)(1-X) = MW_peak_.

#### Native mass spectrometry (MS)

Protein samples (0.50-0.75 mg mL^-1^) were buffer-exchanged to 100 mM ammonium acetate buffer using ZebaSpin columns (Thermo Scientific) and immediately subjected to native MS analysis. Mass spectra were acquired on a Waters Synapt G1 TWIMS MS modified for analysis of intact protein complexes (MS Vision) and equipped with an offline nanospray source. The capillary voltage was 1.5 kV. The cone voltage was set to 50 V, the trap voltage was 20 V and the source temperature was maintained at 20°C. The source pressure was adjusted to 8 mbar. Data were analyzed using MassLynx (Waters).

#### Mouse experiments

Wt mice were purchased from CLEA Japan, Inc. or Japan SLC, Inc. The mutant mice used in this study are available through the Riken BioResource Center (RBRC; https://web.brc.riken.jp/en) under ID *in-frame del*,11009 or via the Center for Animal Resources and Development (CARD; http://card.medic.kumamoto-u.ac.jp/card/english) under ID *in-frame del*,2916.

#### Design and generation of ZP2 ΔN1 mutant mice

The ΔN1 deletion, corresponding to mZP2 residues S40-I144, was designed based on (a) the crystal structure of the mZP2-N1 domain^9^; (b) the predicted (and subsequently experimentally confirmed, Figure S1B) N-terminus of the mZP2-N2 domain, based on secondary structure predictions and homology modeling; and (c) the observation that the two residues immediately before the predicted beginning of mZP2-N2 match the sequence 40-SE-41 that closely follows the signal peptide cleavage site of mZP2 (S34|V35). As a result, the mZP2-N1 deletion of the mice described in this manuscript is different from that of previously reported animals expressing a mutant ZP2 protein that lacked C51-V149 (which replaced the first β-strand of mZP2-N2 with the first strand of mZP2-N1)^23^.

Mutant lines were generated by genetically modifying wt zygotes *in vitro*, using CRISPR/Cas9 genome editing technology. Single guide RNAs (sgRNAs) were designed with CRISPRdirect^91^. Zygotes were electroporated using a NEPA21 Super Electroporator (Nepa Gene) with crRNA/tracrRNA/Cas9 ribonucleoprotein complexes and single strand DNA (ssDNA) with homologous sequence targeting sites^146,147^. The eggs were incubated in KSOM media overnight and two-cell stage embryos were transferred into the ampulla of pseudo-pregnant ICR females. 19 days after egg implantation, offspring (F0) were obtained by natural birth or caesarean section. Pups carrying a mutant genome were screened by PCR and Sanger sequencing. PCR was performed using KOD Fx Neo (KFX-201). The sequence of the ssDNA used for ZP2 ΔN1 is: 5’-TCCACATCACTTACAGATATTAGATCTCTCCTGCAGACAACAATTTCTGACTGAGGAAGGCTTACTGAGTTCACTGAAGTCACAAGGGTGAATAAGAGGG-3’. The sequences of the primers used for genotyping are: ZP2 ΔN1: Fw 5’-CGAGGTGGCAGA GGAAAGCATCTGT-3’, Rv 5’-ATTCATTCATCTGCTAAGCCCAGCT -3’; ZP2-N1 (wt): Fw 5’-CCACTCTAATGGTCAGATGG-3’, Rv 5’- ATTCATTCATCTGCTAAGCCCAGCT-3’.

#### Mouse fertility test

Sexually mature ZP2 ΔN1 female mice (5-16-week-old) and wt B6D2F1 mice were caged with 8-week-old wt male mice for 2 months. During the fertility test, the vaginal plugs and the number of pups at birth were counted every day. The average number of pups/plugs for each mouse line was calculated by dividing the total number of pups by the number of plugs.

#### *In vitro* fertilization (IVF)

Spermatozoa were obtained from the cauda epididymis of adult B6D2F1 males and cultured in Toyoda, Yokoyama, Hoshi (TYH) medium^148^ for 2 h at 37 °C under 5% CO_2_. Oocytes at the second metaphase (MII) stage were collected from the oviducts of females 16 h after injection with pregnant mare’s serum gonadotropin (PMSG) (5 units), followed 48 h later by human chorionic gonadotropin (hCG) (5 units). Wt sperm incubated for 2 h in TYH medium were added to a drop containing cumulus-intact oocytes at a final density of 2 × 10^5^ spermatozoa mL^-1^. 8 h after adding the spermatozoa, two-pronuclear (2-PN) eggs were counted.

#### Sperm-ZP binding assays

Spermatozoa and MII stage oocytes were obtained as described above. Cumulus cells were removed from oocytes by treatment with 330 µg/ml hyaluronidase (Sigma-Aldrich) for 5 min and pipetting. Spermatozoa were incubated for 2 h at 37°C, 5% CO_2_ and added to a drop of TYH medium containing cumulus-free oocytes at a final density of 2 × 10^5^ spermatozoa mL^-1^. 30 min after mixing the spermatozoa, oocytes were fixed with 0.25% glutaraldehyde. Sperm-bound eggs were observed with an IX73 microscope (Olympus).

#### Experiments with *Sof1* knockout mouse sperm

To distinguish whether the infertile phenotype of ZP2 ΔN1 mice was due to impairment of sperm binding or penetration through the ZP, we have taken advantage of recently described *Sof1*^-/-^ mice^77^. These knockout (KO) animals produce sperm that can bind to the ZP and penetrate it, but cannot fuse with the plasma membrane of the oocyte. As a result, sperm accumulates into the perivitelline space and can thus be used to follow and quantify ZP penetration.

For ZP binding experiments, *Sof1* KO sperm were preincubated in TYH medium for 2 h at 37°C, 5% CO_2_. The capacitated sperm were incubated with anti-IZUMO1 antibody and anti-rat IgG Alexa Fluor 488-conjugated antibody (both at a dilution of 1:100) for another 30 min. The eggs harvested from wt and ZP2 ΔN1 mutant female mice were denuded by treating with 0.3 mg mL^-1^ hyaluronidase at 37°C for 5 min. The antibody-probed sperm were added to medium drops containing cumulus-free eggs at a sperm density of 2 × 10^5^ spermatozoa mL^-1^ and incubated for 30 min at 37°C, 5% CO_2_. Sperm-egg complexes were fixed in 0.25% glutaraldehyde for 15 min on ice, rinsed in fresh FHM medium drops for three times, and examined under a Keyence microscope. The total number of sperm and the number of acrosome-reacted sperm bound to the ZP of each egg were recorded. Z-stack images were captured by a Nikon Eclipse Ti microscope equipped with a Nikon C2 confocal module.

For ZP penetration assays, *Sof1* KO sperm were preincubated in TYH medium for 2 h at 37°C, 5% CO_2_. Capacitated sperm were then incubated with cumulus-intact eggs harvested from wt and ZP2 ΔN1 mutant female mice at a sperm concentration of 2 × 10^5^ spermatozoa mL^-1^ for 6 h at 37°C, 5% CO_2_. Sperm-egg complexes were fixed in 0.25% glutaraldehyde for 15 min on ice, rinsed in fresh FHM medium drops for three times, and observed under an Olympus inverted microscope. The number of sperm in the perivitelline space of each egg was counted. Images and videos were captured using a Keyence microscope.

#### ZP hardening assay

Oocytes at the second metaphase (MII) stage were collected as described above. Oocytes and embryos were treated with 10 mg ml^-1^ collagenase (type I; Sigma-Aldrich) to remove the ZP. 10 min after the addition of the collagenase, eggs were repeatedly pipetted and their morphology was observed as described above.

#### Protein deglycosylation

The native fZP1/fZP3 complex and recombinant fZP3 proteins were treated with PNGase F (New England Biolabs and Takara Bio, respectively) for 1 hr at 37°C (native fZP1/fZP3 complex) or overnight at 37°C (recombinant fZP3 proteins), in three different conditions: native (New England Biolabs: 50 mM sodium phosphate pH 7.5; Takara Bio: 100 mM Tris-HCl pH 8.6), semi-denaturing (New England Biolabs: 50 mM sodium phosphate pH 7.5, 0.5% SDS) and fully denaturing (New England Biolabs: 50 mM sodium phosphate pH 7.5, 0.5% SDS, 40 mM DTT; Takara Bio: 100 mM Tris-HCl pH 8.6, 0.1% SDS, 0.15% β-mercaptoethanol).

#### Immunoblot analysis

Protein lysates from MII or fertilized eggs from wt or ZP2 ΔN1 mice were resolved by SDS-PAGE and transferred to PVDF membranes. After blocking, blots were incubated with primary antibodies overnight at 4°C and then incubated with secondary antibodies conjugated to horseradish peroxidase (Jackson ImmunoResearch; 1:10000) for 1 h at RT. Antibodies used: anti-mZP2-N1 monoclonal IE-3^76^ (ATCC CRL-2463; 1:1000); anti-mZP2 cleavage site polyclonal Ab164 (1:1000); anti-mZP2 ZP-C monoclonal 2C7 (1:1000), anti-β-actin (ACTB) (Abcam ab6276; 1:1000). Detection was performed using Chemi-Lumi One Super (Nacalai Tesque). The rabbit polyclonal Ab164 antibody was produced by immunization with a synthetic peptide corresponding to the mZP2 post-fertilization cleavage site (Cys-Ahx-SRLADEN) and purified using the same peptide coupled to SulfoLink resin (Thermo Fisher Scientific). Recombinant fZP3 proteins were detected using anti-His-tag monoclonal (Proteintech) and HRP-conjugated goat anti-mouse IgG (H+L) secondary antibody (Proteintech); EzWestLumi One (ATTO) was used as chemiluminescent substrate.

#### Calculation of the relative stoichiometry of mouse ZP subunits

After digitizing a previously reported high performance liquid chromatograph SEC elution profile of a preparation of ∼30,000 mZPs^38^, A_280 nm_ peak areas corresponding to mZP1-3 were integrated and converted to a relative subunit ratio by taking into account extinction coefficients calculated from the sequences of the respective mature polypeptides.

#### Modeling of the human ZP2 NTR/ovastacin complex

The structure of the complex between hZP2 NTR (residues I39-I366) and the mature form of hOvst (L86-D431) was predicted with AlphaFold-Multimer^149^, using version 3 of its model parameters (released on 13 December 2022) and standard run settings. AlphaFill^79^ was then used to transplant the catalytic Zn^2+^ ion from the crystallographic model of zebrafish hatching enzyme ZHE1 (another member of the astacin-like metalloprotease family; PDB 3LQB)^150^ to the AlphaFold Protein Structure Database^151^ entry for hOvst (https://www.alphafold.ebi.ac.uk/entry/Q6HA08) (optimized transplant clash score 0.14 Å). This was then superimposed on the corresponding chain of the top-ranked complex prediction (ipTM+pTM 0.69; ipTM 0.70), with an RMSD of 0.64 Å over 170 Cα atoms. The resulting hZP2 NTR/hOvst+Zn^2+^ model was finally energy minimized in YASARA Structure, using the YASARA2 force field.

#### Modeling of human ZP filaments

As described above, we initially determined the structure of the fZP1/fZP3 complex by MR using two subcomplex models generated by a ColabFold implementation of AlphaFold. When AlphaFold-Multimer became subsequently available, we found that we could use a single prediction job to generate models of the whole VE fragment that phased the crystallographic data with improved statistics relative to the original two-step procedure. Specifically, a 5 model-ensemble of the fZP1/fZP3 complex produced a single Phaser solution (LLG=274, TFZ=20.2) that could be automatically refined to R_free_=0.40. Based on this result and the experimental evidence that homomeric and heteromeric filaments made by ZP module-containing proteins share the same basic architecture (Figure 5F), we used the AlphaFold-Multimer model parameters to carry out a comprehensive computational analysis of possible subunit-subunit interactions between the four components of the human ZP (hZP1-4), in three steps (Figure S7).

In the first step, we ran a matrix of predictions for ternary complexes consisting of (I-1) a complete hZP subunit; (I-2) a fragment corresponding to the second half of the ZP-N/ZP-C linker and ZP-C domain of any hZP subunit (including the one in (1)), up to its C-terminal consensus furin-cleavage site; and (I-3) a fragment encompassing the full N-terminal region, ZP-N domain and first half of the ZP-N/ZP-C linker of the same subunit (Figure S7A, step I). For type II subunits (hZP1/2/4), this showed that complexes consistent with the filament architecture shared by the VE fragment and the UMOD polymer were obtained only in combination with type I subunit hZP3 (green boxes); moreover, all these filament predictions had significantly higher ranking scores (average ipTM+pTM 0.75) than the non-filamentous arrangements obtained when type II subunits were combined with each other (gray boxes; average ipTM+pTM 0.30), as well as much more convincing Predicted Aligned Error (PAE) heat maps (compare Figure S7B with Figure S7C). At the same time, it is interesting to notice that cross-hairpin interactions like that observed in the fZP1/fZP3 crystal structures were also found in low-scoring, non-filamentous complex predictions (Figure S7C). In the case of the hZP3-only combination (orange box), 3 models out of 5 predicted a filamentous arrangement, whereas the remaining 2 yielded inferences that matched the homodimeric structure of the cZP3 precursor^6^.

In the second step, we generated an additional matrix of predictions for ternary complexes consisting of (II-1) a complete hZP3 subunit; (II-2) a fragment corresponding to the second half of the ZP-N/ZP-C linker and ZP-C domain of any subunit; and (II-3) a fragment encompassing the full N-terminal region, ZP-N domain and first half of the ZP-N/ZP-C linker of any subunit other than the one in (II-2) (Figure S7A, step II). This second analysis extended the previous by indicating that hZP1, hZP2 and hZP4 are in principle able to form all possible mixed ZP-C/ZP-N interfaces between themselves, whereas the majority of combinations involving the domains of the hZP3 ZP module resulted in a lower fraction of filamentous complex predictions.

In the third step, we predicted larger quaternary complexes consisting of 2 copies of hZP3 and: (III-1) 1 copy of the hZP1 trefoil domain+ZP module, 1 copy of the hZP2 ZP module; (III-2) 1 copy of the hZP1 trefoil domain+ZP module, 1 copy of the hZP4 trefoil domain+ZP module; or (III-3) 1 copy of the hZP2 ZP module, 1 copy of the hZP4 trefoil domain+ZP module. Even when using 5 seeds per deep learning model, all predictions resulting from these combinations showed filamentous arrangements that were either circular (reminiscent of early prediction studies of UMOD^30^) or straight (by disconnecting the ZP-N and ZP-C moieties of one subunit) (Figure S7A, step III). Moreover, in both cases, all the predicted filaments (amounting to a total of 75 models) consisted of a hZP3 protofilament combined with a hZP1/2/4 protofilament. In other words, in complete agreement with the crystal structure of the fZP1/fZP3 complex (Figures 5 and 6), homomeric ZP-N/ZP-C interfaces were never observed. Accordingly, no filament-like arrangement were observed in parallel predictions of (III-4) 4 copies of the hZP1 trefoil domain+ZP module; (III-5) 4 copies of the hZP2 module; (III-6) 4 copies of the hZP3 module; (III-7) 4 copies of the hZP4 trefoil domain+ZP module; and (III-8) 2 copies of the hZP2 module, 1 copy of the hZP1 trefoil domain+ZP module and 1 copy of the hZP4 trefoil domain+ZP module (for a total of 125 models).

The information obtained from steps I-III was used for assembling filament models like the ones shown in Figure 7B. To do so, fragments corresponding to ternary complexes generated as described above were superimposed over common protein domains, after which all half-protein copies with truncated ZP-N/ZP-C IDLs were deleted (except the ones at either end of the filament). Despite its simplicity, this procedure did not introduce any significant clash between hZP subunits; as a result, relieving minor unfavorable contacts by standard energy minimization in YASARA Structure was sufficient to achieve very good geometry (for the individual filaments in Figure 7B: MolProbity score 1.06, clashscore 0.83, Ramachandran favored 96.3%, Rama distribution Z-score 0.01±0.10). Moreover, inspection of the polymeric models in PyMOL and analysis with GlyProt^94^ showed that all their possible N-glycosylation sites (corresponding to a total of 116 sites in the case of the filaments shown in Figure 7B) are, as expected, exposed to the solvent.

Like in the mouse and the chicken, human egg coat filaments are covalently cross-linked into a 3D matrix by intermolecular disulfides involving the ZP-N1 domains of hZP1 subunits from different filaments^26,37^. Each of these N-terminal hZP1-N1 domains is spaced from the corresponding trefoil domain and C-terminal ZP module by a ∼90-residue linker, which is recognized as unstructured by both dedicated predictors^152^ and an analysis of AlphaFold model per-residue confidence (pLDDT) values^73,80^. Based on this information, we modeled hZP1 cross-links between hZP filaments by (1) generating a random coil model of the hZP1 disordered spacer region; (2) connecting the N-termini of two copies of this spacer to the C-termini of a model of a disulfide-bonded dimer of hZP1-N1, generated using a combination of MODELLER^99^ and AlphaFold; (3) connecting the C-termini of the resulting 2 x (hZP1-N1 + spacer) assembly to the N-termini of the trefoil domains of hZP1 subunits belonging to two hZP filaments. Clearly, given the intrinsic flexibility of the hZP1 spacer, the semi-extended cross-link of the model depicted in Figure 7B is only one of the many possible conformations that this region of the protein adopts while cross-linking hZP filaments with variable relative orientations. For this reason, the model is strictly meant to provide a chemically realistic example that can be used to draw general conclusions about hZP architecture, rather than recapitulate the variety of filament cross-link conformations that are likely to exist in nature.

To model the non-covalent filament cross-links resulting from post-fertilization processing of hZP2 (Figure 7C), we first used AlphaFold-Multimer to model cleaved hZP2-N2N3. The top ranked prediction had a clearly superior PAE heat map and showed the same relative subunit arrangement observed in our experimental structures of cleaved xZP2-N2N3 (Figure 2J, K) and uncleaved mZP2-N2N3 (Figure 4I), with the βG strand of the hZP2-N2 domain mediating homodimer formation. The two moieties of this model were then superimposed on the ZP-N2 and -N3 domains from two hZP2 subunits belonging to different filaments. To complete the assembly, two copies of the hZP2-N1 domain from the hZP2-N1N2N3 crystal structure were finally linked to the hZP2-N2 N-termini in random relative orientation. Although AlphaFold can accurately predict the structure of the NTR of hZP2, analysis of hZP filament fragment models that include this subunit suggests that the NTR does not appreciably interact with the rest of the protein and can adopt multiple orientations relative to the core of the filament, due to flexibility of the linker between ZP-N3 and the ZP module (Figure S7D). One possible set of orientations was chosen to generate the model depicted in Figure 7C but, as in the case of the hZP1 cross-link, the presence of such a hinge within hZP2 is likely to be important in order to establish non-covalent cross-links between variably oriented filaments. However, even in the extreme case where the longest axis of a cleaved hZP2 NTR homodimer is oriented perpendicular to the filaments, the latter could not be separated by more than the longest dimension of the NTR dimer itself (∼110 Å).

### QUANTIFICATION AND STATISTICAL ANALYSIS

Biochemical and mouse experiments were repeated at least two and three times, respectively; in all cases, results were reproducible. Statistical analyses were performed using Student’s t-test (two-tailed). Data represent the means ± standard deviation (SD) and error bars indicate SD.

### DATA AVAILABILITY

Structure factors and coordinates are deposited in the Protein Data Bank under accession numbers 7ZBI (hZP2-N1N2N3), 7ZBJ (hZP2-N1N2), 7ZBK (cleaved xZP2-N2N3), 7ZBL (mZP2-N2N3), 7ZBM and 8BQU (fZP1/fZP3 complex crystal forms I and II); EM maps of cleaved xZP2-N2N3 are deposited in the Electron Microscopy Data Bank under accession code EMD-14585 (C1) and EMD-14586 (C2). Other relevant data and materials are available from the corresponding author upon reasonable request.

## ACKNOWLEDGEMENTS

This work was supported by the Knut and Alice Wallenberg Foundation (grant 2018.0042 to L.J.); the Swedish Research Council (grants 2016-03999 and 2020-04936 to L.J.; grant 2019–01961 to M.L.); the Center for Innovative Medicine (grant 2-537/2014 to L.J.); the German Research Foundation (grant 401763100 to D.F.); NIH (grant R01 GM125638 to A.E.C.) and JSPS (grant 20K06774 to S.Y; grants 19H05750 and 21H05033 to M.I.). We thank the staff of the European Synchrotron Radiation Facility (ESRF), Diamond Light Source (DLS) and the Swedish National Cryo-EM Facility in Stockholm for help with X-ray and cryo-electron microscopy data collection; Blanca Algarra (Karolinska Institutet) for initial work on ZP2 cleavage; Kathryn Tunyasuvunakool (DeepMind) for initial models of monomeric fish VE subunits; Gérard Bricogne and Clemens Vonrhein (Global Phasing Ltd.) for help with deposition of STARANISO-derived diffraction data; Paul Wassarman (Icahn School of Medicine at Mount Sinai) for the hybridoma line producing monoclonal 2C7 and discussion; Franco Cotelli (University of Milano) and Tsukasa Matsuda (Fukushima University) for comments.

## AUTHOR CONTRIBUTIONS

Protein expression and purification: S.N., D.F., E.D., L.H.; X-ray crystallography: S.N., B.W., D.F., E.D., S.Z.-C., M.B., A.S., D.dS., L.J.; cryo-EM: B.W.; AlphaFold predictions: S.N., B.W., L.J.; native MS: M.L.; *Xenopus* egg exudate collection: R.E.B, K.M.K., A.E.C.; mouse experiments: C.E., Y.L., M.K., M.I.; fish VE complex preparation: S.Y. The project was conceived and directed by L.J., who wrote the manuscript together with S.N. and A.S., with contributions from the other authors.

## ADDITIONAL INFORMATION

Correspondence and requests for materials should be addressed to Luca Jovine.

## SUPPLEMENTAL FIGURE LEGENDS

**Figure S1.**
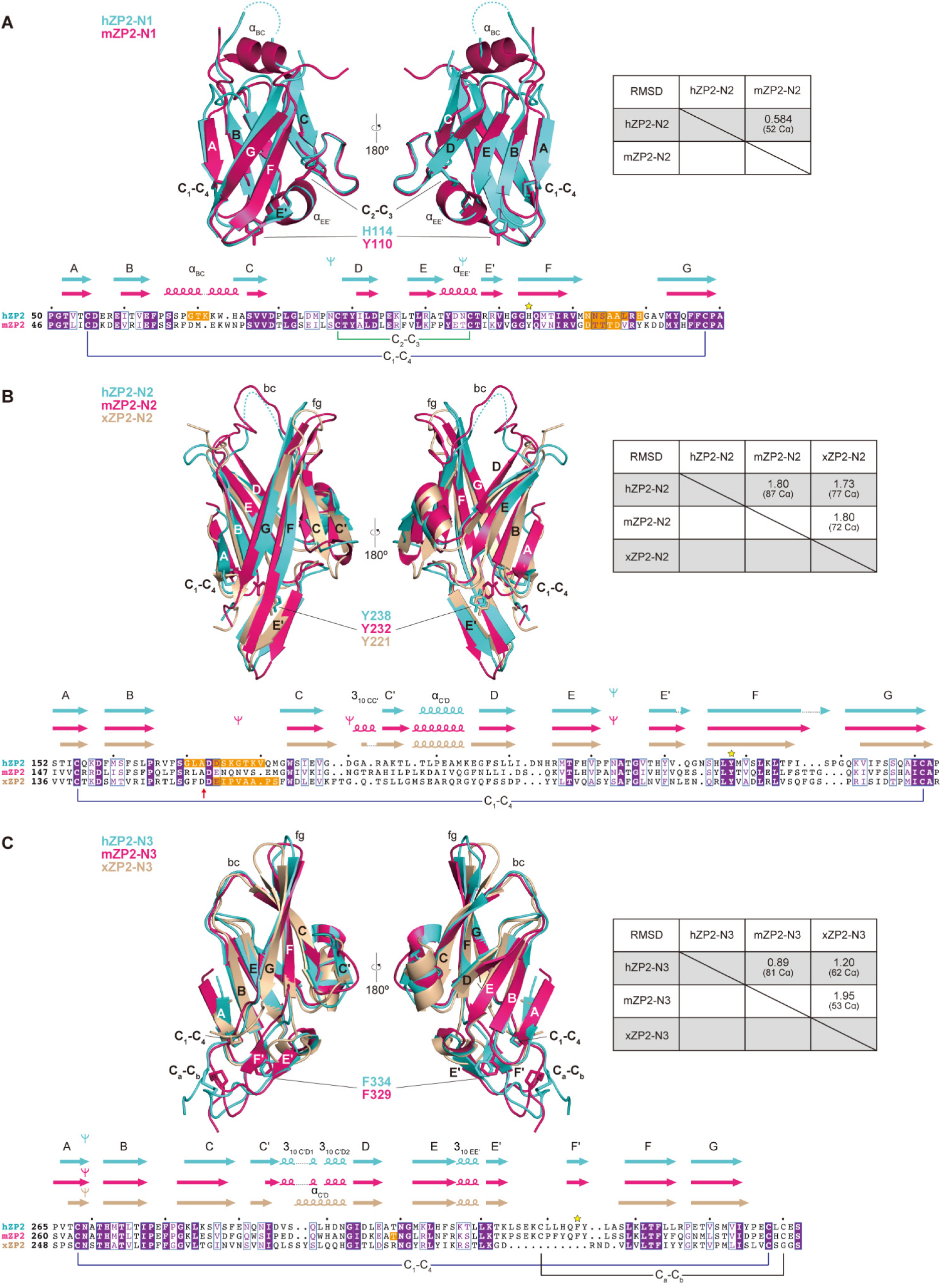
Structural comparisons of the ZP-N domains of ZP2 NTR, related to Figure 1. (A) The ZP-N1 domains of hZP2 and mZP2 (PDB ID 5II6) adopt a typical ZP-N fold, characterized by 8 β-strands and two intramolecular disulfides with 1-4, 2-3 connectivity. Inverted tripods, N-glycan sites; yellow stars, positions corresponding to the signature Tyr of the ZP-N domain. Amino acids highlighted in orange are disordered in the crystals structures and were thus aligned based on sequence only. (B) The ZP-N2 domains of hZP2, mZP2 and xZP2 lack the C_2_-C_3_ disulfide. The red vertical arrow below the sequence alignment indicates the position of the post-fertilization cleavage site. (C) ZP-N3 domains of hZP2, mZP2 and xZP2. Like ZP-N2, ZP-N3 also lacks the second classic ZP-N disulfide. Note that the mammalian ZP2 ZP-N3 domain has a lower secondary structure content at the level of its E’FG extension but is stabilized by an additional C_a_-C_b_ disulfide bond.

**Figure S2.**
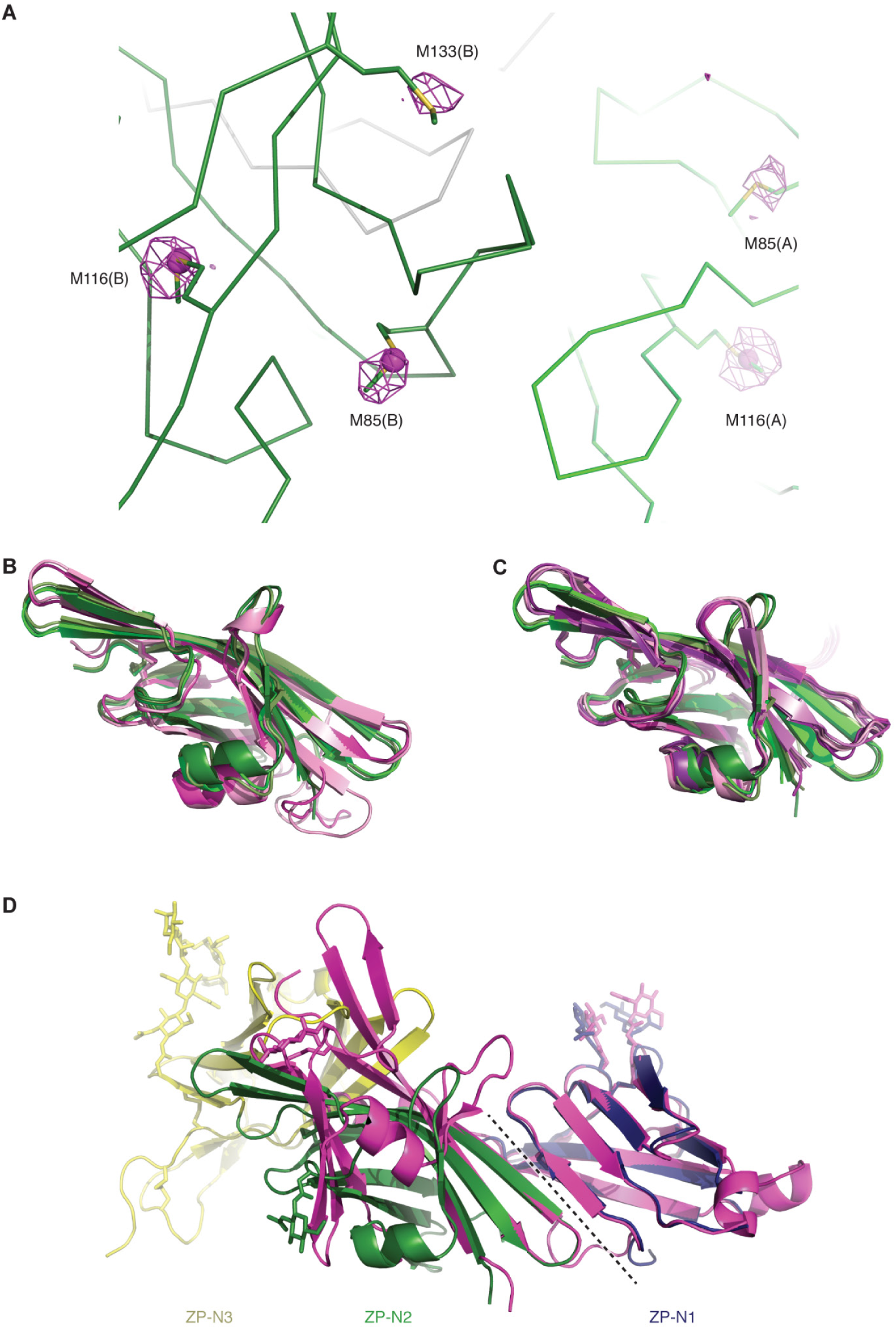
Structure determination and analysis of hZP2-N1N2, related to Figure 1. (A) Validation of the MR solutions for hZP2-N1N2. A section of the hZP2-N1N2 model is shown in Cα trace representation, with the side chains of Met residues shown as sticks. Chain A and B are light and dark green, respectively, while symmetry-related molecules are gray. The violet mesh is an anomalous difference map contoured at 3.5 σ, calculated from SeMet-hZP2-N1N2 data collected at λ=0.92 Å. Note how the map, calculated with phases from the experiment and the refined model, also shows clear peaks for Se atoms that were not included among the sites initially identified by PHENIX AutoSol (shown as magenta spheres). (B) Superposition of the ZP-N2 domains of hZP2-N1N2N3 and mZP2-N2N3 (average RMSD 2.5 Å). The three chains in the asymmetric unit of the hZP2-N1N2N3 are shown in cartoon representation and colored in shades of green, whereas the two chains in the asymmetric unit of the mZP2-N2N3 are colored in shades of magenta. Disulfide bonds are shown as sticks. (C) Superposition of the ZP-N2 domains of hZP2-N1N2N3 and cleaved xZP2-N2N3 (average RMSD 1.9 Å). Molecules are represented as in panel B, with the hZP2-N1N2N3 chains colored in shades of green and the eight chains in the asymmetric unit of the cleaved xZP2-N2N3 crystal colored in shades of magenta. (D) Superposition of the structures of hZP2-N1N2N3 (chain A, colored by domain as in Figure 1A, B) and hZP2-N1N2 (chain A, colored magenta) over ZP-N1 and part of β-strands F and G of ZP-N2 (RMSD 0.4 Å over 75 Cα). N-glycans and disulfides are depicted in stick representation, and the conserved interface between ZP-N1 and ZP-N2 domains is indicated by a dashed black line.

**Figure S3.**
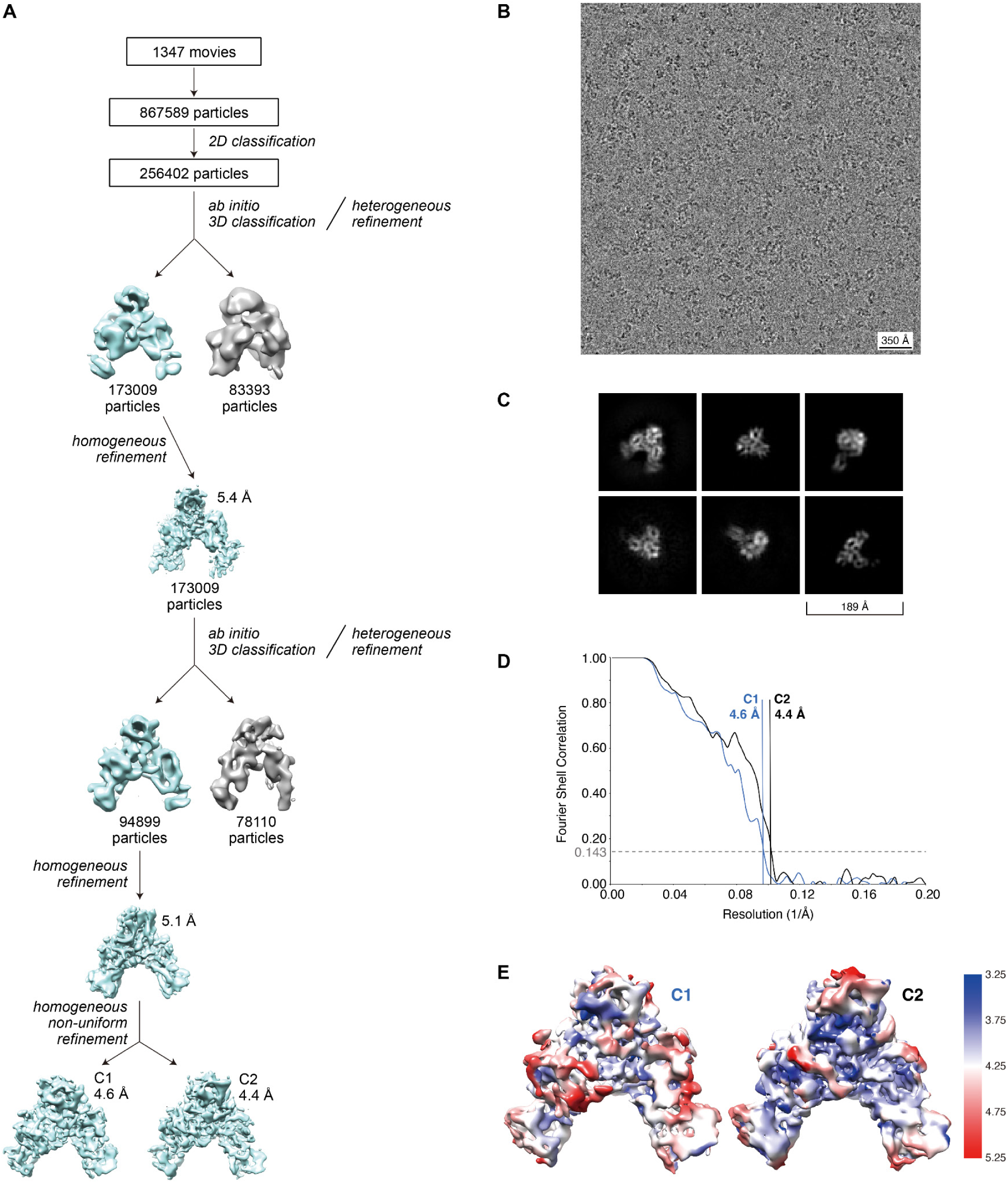
Workflow and data assessment for the cryo-EM structure of cleaved xZP2-N2N3, related to Figure 2. (A) Cryo-EM particle processing workflow. Auto-picked particles were filtered through one round of 2D classification followed by two rounds of *ab initio* 3D classification. The use of homogeneous non-uniform refinement as well as the application of C2 symmetry to the final reconstruction resulted in a modest improvement in resolution. (B) Representative micrograph from a single data collection of 1347 micrographs used for automated particle picking for 2D classification. (C) 2D class averages of cleaved xZP2-N2N3. (D) Fourier Shell Correlation (FSC) of the final volumes. (E) Local-resolution estimation of the C1 and C2 density maps.

**Figure S4.**
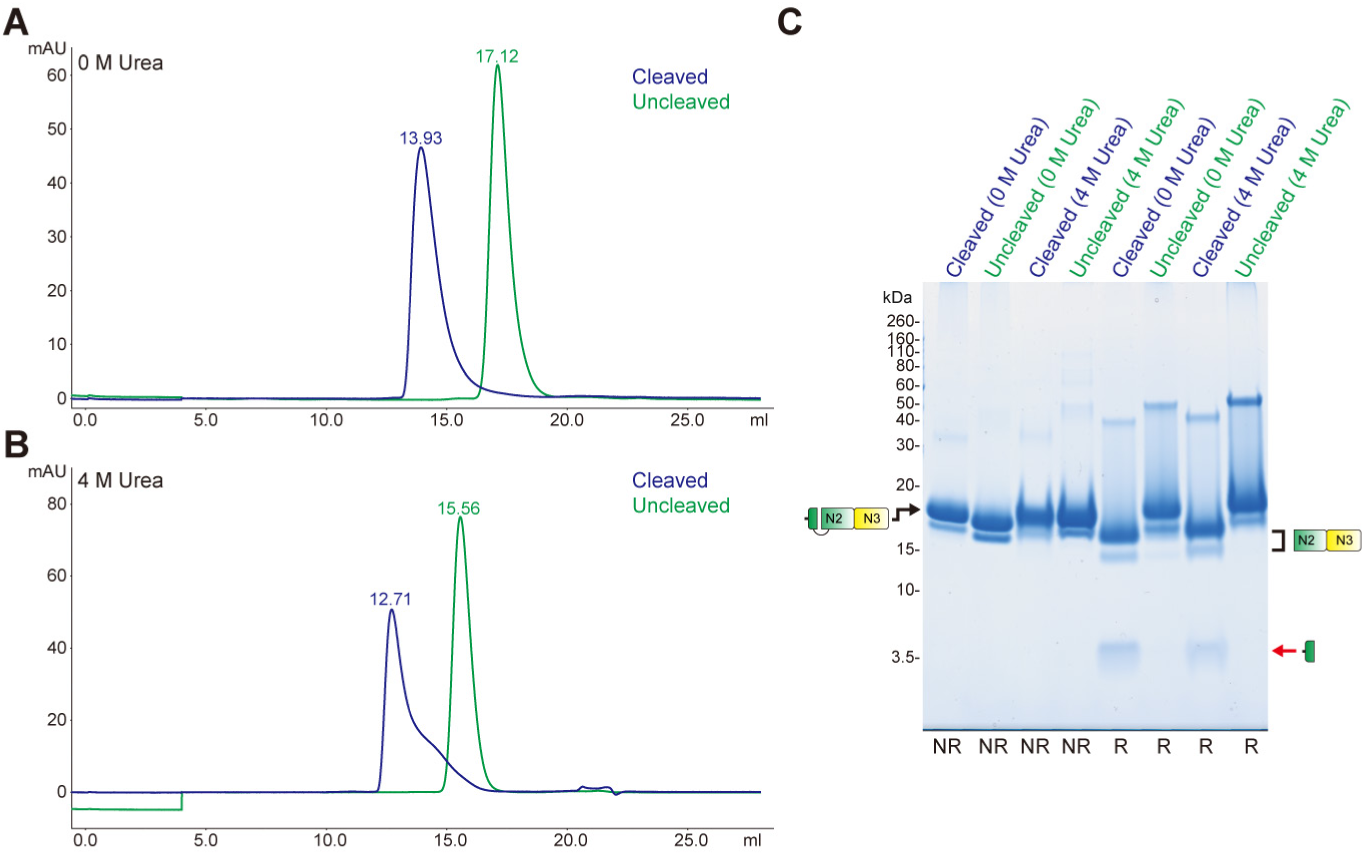
Stability of the xZP2 dimer of dimers, related to Figure 2. (A, B) SEC profiles of uncleaved and cleaved xZP2-N2N3 in the absence (A) and presence (B) of 4 M urea. (C) SDS-PAGE analysis of peaks from the SEC runs shown in panels A and B. A red arrow indicates the N-terminal fragment of xZP2.

**Figure S5.**
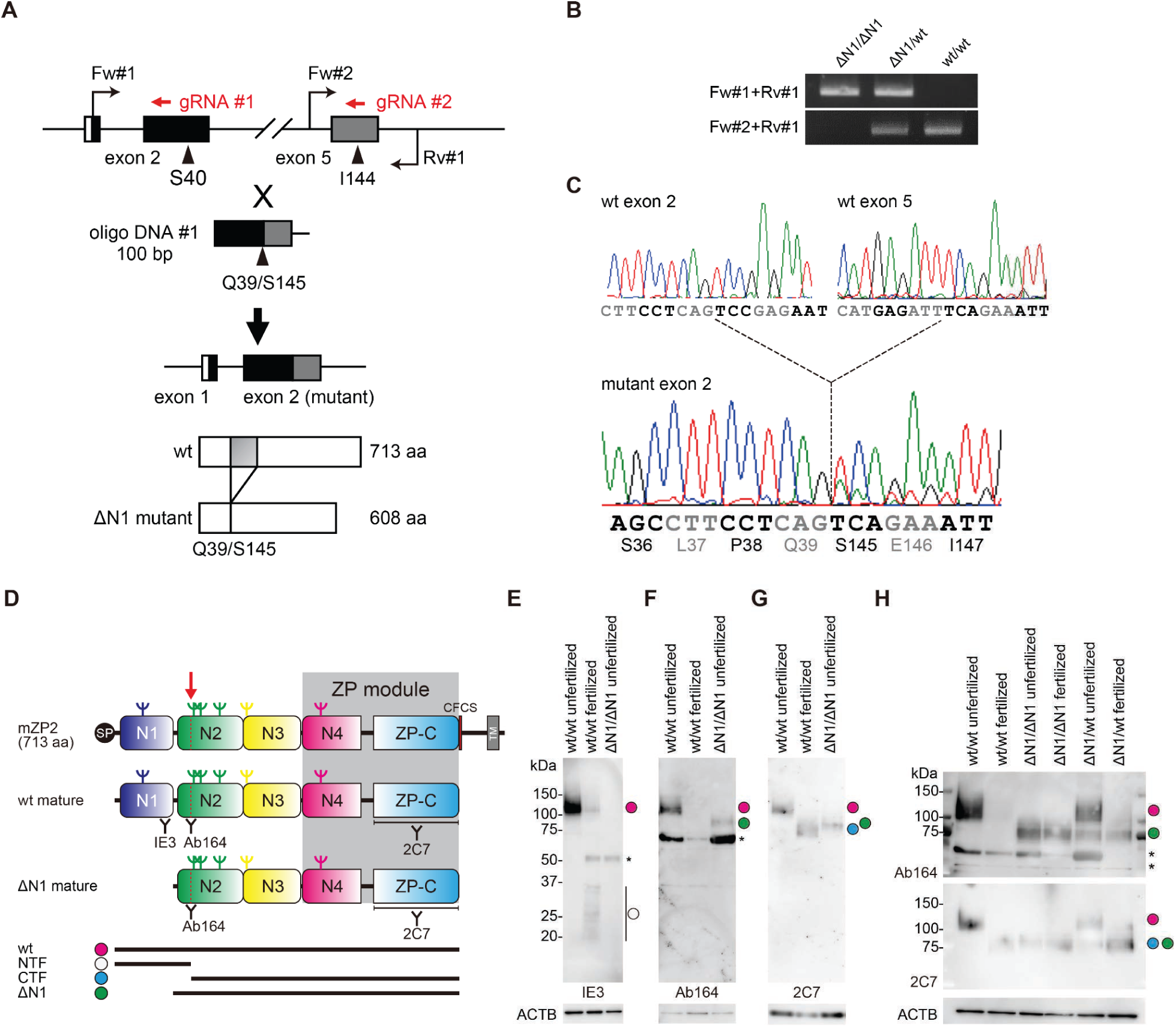
Generation of ZP2 ΔN1 oocytes and analysis of their ZP2 cleavage status, related to Figure 4. (A-C) Generation of ZP2 ΔN1 mice: strategy for generating the ZP2 ΔN1 mutant mice (A), PCR amplification of the mutated genomic region (B) and DNA sequencing of ZP2 ΔN1 mice (C). (D-H) ZP2 immunoblotting of wt and ΔN1 oocytes. (D) Schematic diagram of the mZP2 precursor and the mature forms of the wt protein and its ΔN1 mutant. Red arrow, post-fertilization cleavage site of ZP-N2. Inverted tripods, N-glycans. SP, signal peptide; CFCS, consensus furin cleavage site; TM, transmembrane domain; NTF and CTF, N- and C-terminal fragments generated by the post-fertilization cleavage of mZP2. Monoclonal antibodies IE-3 and 2C7 recognize the ZP-N1 and ZP-C domains of mZP2, respectively. Rabbit polyclonal antibody Ab164 recognizes the uncleaved bc loop of mZP2-N2 (S164-N170). (E-G) Reducing immunoblot analysis of oocytes from wt and ΔN1 homozygous mice shows that the mZP2-N2 cleavage loop of mutant animals is intact before fertilization. mZP2 is detected by IE-3 (E), Ab164 (F) and 2C7 (G). ACTB, β-actin internal control. Asterisks indicate non-specific bands. (H) Reducing immunoblot analysis of wt and ΔN1 homozygous or heterozygous oocytes with Ab164 and 2C7. Note how the mZP2-N2 cleavage loop remains intact even after ΔN1/ΔN1 animals are fertilized by IVF (by making holes in the ZP) and how, in heterozygous mice that express both wt and ΔN1 mutant protein, mOvst only cleaves the former. These observations agree with the fact that hZP2-N1 is required for efficient cleavage of hZP2 by hOvst *in vitro* (Figure 3E), and are also consistent with the idea that the loop/loop interaction observed in the structure of homodimeric mZP2-N2N3 (Figure 4I) makes the mZP2 cleavage site less accessible to the protease.

**Figure S6.**
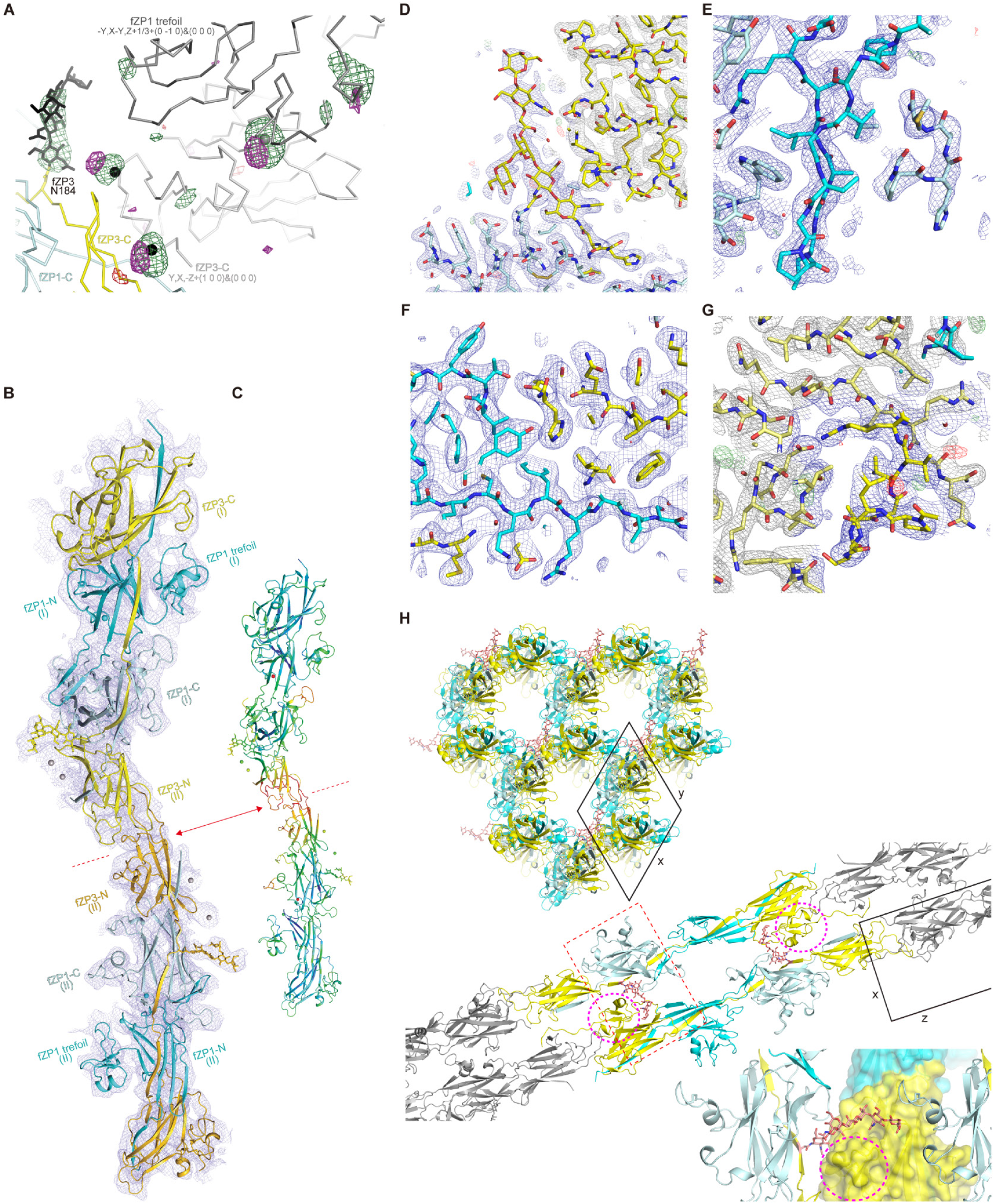
Structure determination and analysis of the native fish VE fragment, related to Figures 5 and 6. (A, B) Validation of the single Phaser MR solution for crystal form I of the fZP1/fZP3 complex. (A) Inspection of the sigma-A-weighted *F*_obs_*−F*_calc_ map obtained upon initial refinement of the solution (green mesh, 3 σ; red mesh, -3 σ) revealed strong positive difference density for the glycan chain attached to fZP3 N184, which is shown here (black sticks) but was not included in the search model used for MR and initial refinement. Four additional large positive difference density peaks were also observed, which have been tentatively assigned to Yb^3+^ ions from the mother liquor (spheres) based on the fact that they match peaks in a phased anomalous difference map calculated from the same data (magenta mesh, 3.8 σ). Molecules are shown as Cα traces, with the copy of the fZP1/fZP3 complex contained in the reference asymmetric unit of the crystal (XYZ) colored as in Figure 5A and two symmetry-related molecules colored light and dark gray. (B) 4.2 Å resolution sigma-A-weighted *2F*_obs_*−F*_calc_ electron density map after DEN refinement, contoured at 1 σ and carved at 4 Å around the complex model in the reference asymmetric unit (complex copy (I), above the red dotted line) as well as a copy related by symmetry operation -X,-X+Y,-Z+1/3+(2 1 0)&(0 0 0) (complex copy (II), below the red dotted line). Note how the relative arrangement of the two copies of the complex mimics a filament, although their relative orientation is inverted compared to the one that these fragments must necessarily adopt in the native VE filament (due to the covalent continuity of the fZP1 IDL that connects them before hatching enzyme cleavage). (C) Cartoon model of the same complexes depicted in panel B, shown at a smaller scale and colored by B factor from blue (low) to red (high). Consistent with the above considerations and reflecting the poor quality of this part of the map, high model B factors at the level of the fZP3 ZP-N/ZP-N interaction that connects the two complexes (red double-headed arrow) clearly identify the latter as a non-native crystal contact. (D-G) Representative sections of the refined sigma-A-weighted *2F*_obs_*−F*_calc_ and *F*_obs_*−F*_calc_ electron density maps at 2.7 Å resolution of crystal form II of the fZP1/fZP3 complex. The *2F*_obs_*−F*_calc_ map, contoured at 1.0 σ and carved at 4 Å around the molecular model (in stick representation and colored as in Figure 5A), is colored blue and dark gray for the complex in the reference asymmetric unit and symmetry-related ones, respectively; the *F*_obs_*−F*_calc_ map is also carved at 4 Å around the model and colored green (3.5 σ) and red (-3.5 σ). (D) The fZP3 IDL N-glycan (center) packs against the ZP-C subdomain of a symmetry-related fZP3 molecule from a different filament (top right). (E) View of the ZP-C/ZP-N interface of two fZP1 subunits, centered on the ZP-N βF’ and βF” strands packing against ZP-C W475 and P521. (F) Detail of the fZP1 ZP-N/fZP3 ZP-C interface, involving ZP-N Tyr 339 (βF) and 371 (βG). (G) The fZP3 ZP-C/fZP3 ZP-N interface, centered on the ZP-C D279/ZP-N R167 salt bridge. (H) Cross section (XY plane; top) and longitudinal (XZ plane; middle) views of the packing of crystal form II, highlighting how the fZP3 glycans of one pseudo-infinite filament (pink sticks) interact with the fZP3 ZP-C subdomains of another (dashed magenta circles). fZP1/fZP3 complex subunits are colored as in Figure 5A, except in the middle panel where this is only done for two neighboring asymmetric units. The black box represents the trigonal unit cell and the dashed red rectangle indicates the fZP3 N-glycan/ZP-C subdomain contact region magnified in the bottom panel. The latter depicts one of the filaments in surface representation, with the ZP-C subdomain cartoon in red, and shows how the tip of the fZP3 N-glycan also packs against the fZP1 ZP-C loop that contains the C_x_-C_y_ disulfide.

**Figure S7.**
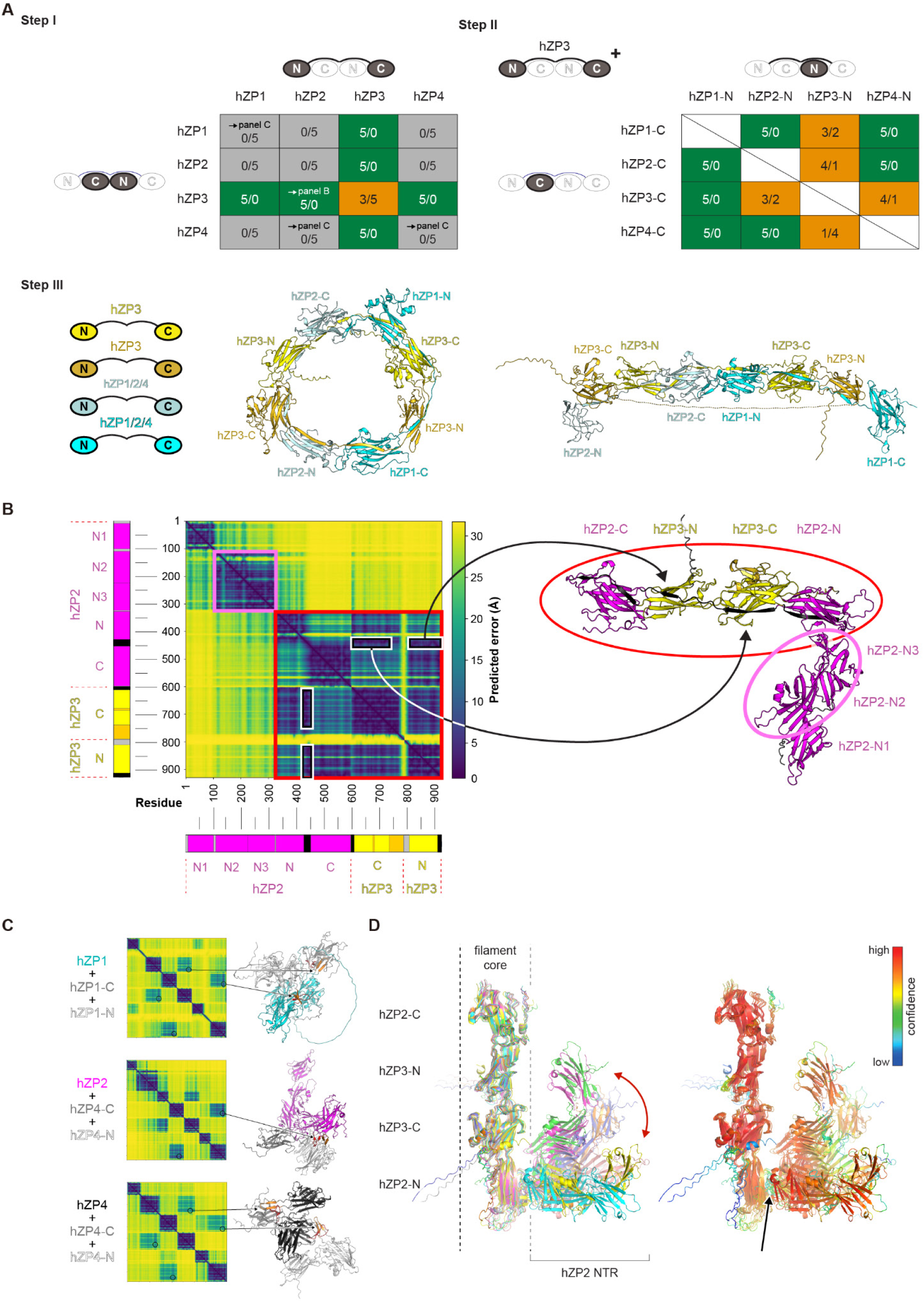
AlphaFold interaction matrix of human ZP subunits, related to Figure 7. (A-C) Schematic summary of the three sets of AlphaFold prediction runs used to probe possible subunit interactions within hZP filaments. (A) Based on the experimental architecture of the fish VE fragment (Figure 5), step I tested all possible combinations between any full-length hZP component and the two moieties that are generated by dividing every subunit (including itself) in the center of its ZP-N/ZP-C linker (dark ovals). Each prediction was performed using 5 distinct deep learning models; combinations that resulted in a filament conformation in all cases, in some cases only and in no cases are indicated by green, orange and gray boxes, respectively. Step II tested all possible combinations between a full-length hZP3 subunit and mixed moieties from all subunits, whereas step III predicted quaternary complexes consisting of two type I (hZP3) and two type II (hZP1/2/4) components. Notably, the latter adopted a circular filament arrangement (middle panel) or a straight filament conformation (right panel) resulting from the non-physical stretching of the ZP-N/ZP-C linker of one subunit (dotted line). For more details, please refer to the corresponding section of the Method details. (B) Step I AlphaFold prediction for a type I/type II hZP subunit combination, the ternary complex of a full-length hZP3 subunit and two moieties from hZP2. The left half of the panel shows the Predicted Aligned Error (PAE) heat map of the top ranked prediction of the complex, whose model is shown on the right, in relation to the domain structure of its components (depicted as bars with bands color-matched to the model). By reporting the predicted error (in Å) between all pairs of residues, AlphaFold’s PAE heat maps can be used to identify domain interactions as areas of low score (colored dark blue in this panel’s maps)^80^. The white boxes and corresponding arrows indicate the clearly recognizable interactions between the N- and C- terminal halves of the IDL of hZP3 and the ZP-C and ZP-N domains of hZP2, respectively; the light magenta square and oval show how the hZP2 ZP-N2 and -N3 domains extensively interact with each other (as observed in the crystal structures of hZP2 (Figure 1B), xZP2 (Figure 2J, K) and mZP2 (Figure 4I) but not with the rest of the complex; the red box and oval encompass the large number of interactions at the level of the filament core. (C) Top ranking step I predictions for different combinations of type II hZP components (hZP1/2/4). Despite having low ipTM+pTM scores and lacking a filament conformation, these models include cross-hairpin ZP-N/ZP-C interactions between subunits (indicated by arrows and orange/red colors for the ZP-N and ZP-C hairpins, respectively). (D) Superposition of multiple models of the complex between full-length hZP2 and two hZP3 moieties, colored by model (left) or by pLDDT (right). As also shown in panel B, the NTR of hZP2 is predicted to make little interaction with the protein’s ZP module or the rest of the complex; instead, the NTR adopts a range of different orientations relative to the core of the filament (red double-headed arrow) due to flexibility (and thus relatively low prediction confidence) of the short linker between hZP2-N3 and the ZP module (black arrow).

**Table S1.** ZP2 X-ray data collection and refinement statistics, related to Figures 1, 2, 4, S1 and S2.

|  | hZP2-N1N2N3<br>(PDB 7ZBI) | hZP2-N1N2<br>(PDB 7ZBJ) | Cleaved<br>xZP2-N2N3<br>(PDB 7ZBK) | mZP2-N2N3<br>(PDB 7ZBL) |
| --- | --- | --- | --- | --- |
| **Data collection** |  |  |  |  |
| Space group | C2 (5) | *P*3<sub>1</sub>21 (152) | *P*2<sub>1</sub> (4) | C2 (5) |
| Cell dimensions |  |  |  |  |
| *a*, *b*, *c* (Å) | 147.68, 97.77, 104.33 | 86.93, 86.93, 178.68 | 88.21, 70.31, 149.86 | 163.01, 54.17, 68.66 |
| $\alpha$, $\beta$, $\gamma$ (°) | 90, 94.803, 90 | 90, 90, 120 | 90, 90.037, 90 | 90, 107.984, 90 |
| Wavelength (Å) | 0.9184 | 0.9538 | 1.7712 | 0.9194 |
| Resolution range (Å) | 21.9 - 2.7 (2.80 - 2.70)<sup>§</sup> | 19.9 - 3.2 (3.31 - 3.20) | 38.0 - 2.7 (2.80 - 2.70) | 43.7 - 1.9 (1.98 - 1.90) |
| Unique reflections | 40494 (3976) | 13385 (1296) | 48609 (3655) | 44849 (4926) |
| Multiplicity | 6.9 (6.4) | 9.7 (9.8) | 3.1 (2.2) | 5.2 (5.4) |
| Completeness (%) | 99.2 (98.0) | 99.3 (99.2) | 95.3 (71.1) | 99.2 (98.5) |
| Mean *I* / σ*I* | 15.7 (0.9) | 11.8 (0.6) | 4.2 (0.6) | 12.6 (0.8) |
| Wilson B-factor (Å<sup>2</sup>) | 79.7 | 130.3 | - | 40.7 |
| *R*<sub>merge</sub> | 0.082 (2.152) | 0.154 (3.808) | 0.253 (1.306) | 0.065 (2.330) |
| *R*<sub>meas</sub> | 0.089 (2.345) | 0.163 (4.018) | 0.306 (1.683) | 0.072 (2.572) |
| *R*<sub>pim</sub> | 0.034 (0.916) | 0.052 (1.275) | 0.170 (1.045) | 0.031 (1.074) |
| CC<sub>1/2</sub> | 1.00 (0.51) | 1.00 (0.34) | 0.96 (0.17) | 1.00 (0.49) |
| CC* | 1.00 (0.82) | 1.00 (0.71) | 0.99 (0.54) | 1.00 (0.81) |
| **Refinement** |  |  |  |  |
| Reflections | 40494 (3952) | 13385 (1296) | 48609 (3591) | 44849 (4915) |
| *R*<sub>free</sub> reflections | 1575 (159) | 1320 (128) | 1988 (148) | 2223 (240) |
| *R*<sub>work</sub> | 0.236 (0.382) | 0.269 (0.412) | 0.217 (0.270) | 0.214 (0.388) |
| *R*<sub>free</sub> | 0.284 (0.419) | 0.312 (0.418) | 0.255 (0.294) | 0.248 (0.394) |
| Number of non-H atoms | 6863 | 3472 | 12412 | 3824 |
| Macromolecules | 6591 | 3402 | 12058 | 3467 |
| Ligand | 239 | 70 | 154 | 170 |
| Solvent | 33 | 0 | 200 | 187 |
| Protein residues | 842 | 438 | 1540 | 438 |
| R.m.s. deviations |  |  |  |  |
| Bond lengths (Å) | 0.006 | 0.004 | 0.004 | 0.005 |
| Bond angles (°) | 0.86 | 0.70 | 0.69 | 0.79 |
| Ramachandran |  |  |  |  |
| Favored (%) | 97.4 | 97.2 | 96.4 | 98.4 |
| Allowed (%) | 2.6 | 2.8 | 3.6 | 1.6 |
| Outliers (%) | 0.0 | 0.0 | 0.0 | 0.0 |
| Rotamer outliers (%) | 0.1 | 2.4 | 0.5 | 0.0 |
| MolProbity score | 1.42 | 1.77 | 1.66 | 0.94 |
| Clashscore | 5.54 | 6.13 | 7.65 | 1.81 |
| Average B-factor (Å<sup>2</sup>) | 118.4 | 157.0 | 53.4 | 59.6 |
| Macromolecules | 117.3 | 156.8 | 53.5 | 58.2 |
| Ligand | 155.3 | 166.9 | 78.1 | 99.1 |
| Solvent | 71.7 | - | 29.6 | 50.7 |

<sup>§</sup> Values in parentheses are for the highest resolution shell

**Table S2.** Cleaved xZP2-N2N3 cryo-EM data collection statistics, related to Figures 2 and S3.

|  | **Cleaved**<br>**xZP2-N2N3**<br>**(EMD-14585)** | **Cleaved**<br>**xZP2-N2N3**<br>**(EMD-14586)** |
| --- | --- | --- |
| Magnification | 15000x | 15000x |
| Voltage (kV) | 200 | 200 |
| Electron exposure (e-/Å<sup>2</sup>) | 52 | 52 |
| Exposures (no.) | 1347 | 1347 |
| Defocus range (µm) | 1.0-3.2 | 1.0-3.2 |
| Pixel size (Å) | 0.9459 | 0.9459 |
| Initial particle images (no.) | 867589 | 867589 |
| Final particle images (no.) | 94899 | 94899 |
| Symmetry imposed | C2 | C1 |
| Map resolution (Å) | 4.4 | 4.6 |
| FSC threshold | 0.143 | 0.143 |
| Map resolution range (Å) | 9.3-2.3 | 13-3.3 |

**Table S3.** fZP1/fZP3 complex X-ray data collection and refinement statistics, related to Figures 5, 6 and S6.

|  | Crystal form I<br>(PDB 7ZBM) |  | Crystal form II<br>(PDB 8BQU) |  |
| --- | --- | --- | --- | --- |
|  |  | Isotropic, standard<br>(TRUNCATE)<sup>§</sup> | Isotropic, extended<br>(AIMLESS)<sup>§§</sup> | Anisotropic<br>(STARANISO)<sup>§§§</sup> |
| Data collection |  |  |  |  |
| Space group | *P*3<sub>1</sub>21 (152) | *P*3<sub>2</sub>21 (154) | *P*3<sub>2</sub>21 (154) | *P*3<sub>2</sub>21 (154) |
| Cell dimensions |  |  |  |  |
| *a*, *b*, *c* (Å) | 108.35, 108.35, 255.07 | 59.06, 59.06, 473.92 | 59.06, 59.06, 473.92 | 59.06, 59.06, 473.92 |
| α, β, γ (°) | 90, 90, 120 | 90.0 90.0 120.0 | 90.0 90.0 120.0 | 90.0 90.0 120.0 |
| Wavelength (Å) | 1.0723 | 0.9794 | 0.9794 | 0.9794 |
| Resolution range (Å) | 88.1-4.2 (4.52-4.20)<sup>§</sup> | 50.8-3.47 (3.53-3.47) | 51.1-2.42 (2.46-2.42) | 51.1-2.42 (2.46-2.42) |
| Unique reflections | 13155 (2600) | 13278 (611) | 38183 (1886) | 14348 (720) |
| Multiplicity | 6.4 (6.8) | 4.7 (4.7) | 4.8 (4.4) | 4.4 (3.1) |
| Completeness, spherical (%) | 98.5 (99.2) | 99.1 (99.5) | 99.1 (97.9) | 37.3 (6.1) |
| Completeness, ellipsoidal (%) | - | - | 98.1 (72.8) | 89.9 (70.0) |
| Mean *I* / σ*I* | 4.4 (0.8) | 5.8 (2.4) | 2.4 (0.1) | 6.2 (2.2) |
| Wilson B-factor (Å²) | 203.2 | 66.0 | 27.7 | 19.7 |
| R<sub>merge</sub> | 0.207 (2.457) | 0.230 (0.753) | 0.675 (187) | 0.199 (0.615) |
| R<sub>meas</sub> | 0.224 (2.649) | 0.259 (0.844) | 0.756 (212) | 0.225 (0.734) |
| R<sub>pim</sub> | 0.085 (0.965) | 0.115 (0.375) | 0.333 (97) | 0.102 (0.389) |
| CC<sub>1/2</sub> | 0.99 (0.71) | 0.99 (0.95) | 0.98 (0.03) | 0.99 (0.72) |
| CC* | 1.00 (0.91) | - | - | - |
| Refinement |  |  |  |  |
| Resolution range (Å) | 88.1-4.2 (4.52-4.20) |  |  | 51.1-2.70 (2.91-2.70) |
| Reflections | 13155 (2583) |  |  | 13792 (747) |
| R<sub>free</sub> reflections | 652 (127) |  |  | 646 (35) |
| R<sub>work</sub> | 0.329 (0.434) |  |  | 0.247 (0.351) |
| R<sub>free</sub> | 0.368 (0.436) |  |  | 0.307 (0.480) |
| Number of non-H atoms | 4656 |  |  | 4852 |
| Macromolecules | 4613 |  |  | 4766 |
| Ligand | 77 |  |  | 162 |
| Solvent | 0 |  |  | 0 |
| Protein residues | 599 |  |  | 622 |
| R.m.s. deviations |  |  |  |  |
| Bond lengths (Å) | 0.003 |  |  | 0.003 |
| Bond angles (°) | 0.63 |  |  | 0.60 |
| Ramachandran |  |  |  |  |
| Favored (%) | 95.6 |  |  | 97.4 |
| Allowed (%) | 4.1 |  |  | 2.6 |
| Outliers (%) | 0.3 |  |  | 0.0 |
| Rotamer outliers (%) | 0.0 |  |  | 0.0 |
| MolProbity score | 1.46 |  |  | 0.94 |
| Clashscore | 3.62 |  |  | 1.16 |
| Average B-factor (Å²) | 286.5 |  |  | 25.5 |
| Macromolecules | 286.5 |  |  | 25.4 |
| Ligand | 281.9 |  |  | 33.8 |

<sup>§</sup> Values in parentheses are for the highest resolution shell
<sup>§</sup> Criteria used in determination of resolution cutoff: *R*<sub>pim</sub> ≤ 0.6000, *I*/ $\sigma$ *I* ≥ 2.00, CC(1/2) ≥ 0.3000 (autoPROC default)
<sup>§§, §§§</sup> Criteria used in determination of diffraction limits: local(*I*/ $\sigma$ *I*) ≥ 1.20 (autoPROC default)

## REFERENCES

1. Wong, J.L., and Wessel, G.M. (2006). Defending the zygote: search for the ancestral animal block to polyspermy. Curr. Top. Dev. Biol. 72, 1–151. https://doi.org/10.1016/S0070-2153(05)72001-9.

2. Evans, J.P. (2020). Preventing polyspermy in mammalian eggs—Contributions of the membrane block and other mechanisms. Mol. Reprod. Dev. 87, 341–349. https://doi.org/10.1002/mrd.23331.

3. Bleil, J.D., Beall, C.F., and Wassarman, P.M. (1981). Mammalian sperm-egg interaction: Fertilization of mouse eggs triggers modification of the major zona pellucida glycoprotein, ZP2. Dev. Biol. 86, 189–197. https://doi.org/10.1016/0012-1606(81)90329-8.

4. Tian, J., Gong, H., and Lennarz, W.J. (1999). *Xenopus laevis* sperm receptor gp69/64 glycoprotein is a homolog of the mammalian sperm receptor ZP2. Proc. Natl. Acad. Sci. U. S. A. 96, 829–834. https://doi.org/10.1073/pnas.96.3.829.

5. Burkart, A.D., Xiong, B., Baibakov, B., Jiménez-Movilla, M., and Dean, J. (2012). Ovastacin, a cortical granule protease, cleaves ZP2 in the zona pellucida to prevent polyspermy. J. Cell Biol. 197, 37–44. https://doi.org/10.1083/jcb.201112094.

6. Han, L., Monné, M., Okumura, H., Schwend, T., Cherry, A.L., Flot, D., Matsuda, T., and Jovine, L. (2010). Insights into egg coat assembly and egg-sperm interaction from the X-ray structure of full-length ZP3. Cell 143, 404–415. https://doi.org/10.1016/j.cell.2010.09.041.

7. Callebaut, I., Mornon, J.-P., and Monget, P. (2007). Isolated ZP-N domains constitute the N-terminal extensions of Zona Pellucida proteins. Bioinformatics 23, 1871–1874. https://doi.org/10.1093/bioinformatics/btm265.

8. Monné, M., Han, L., Schwend, T., Burendahl, S., and Jovine, L. (2008). Crystal structure of the ZP-N domain of ZP3 reveals the core fold of animal egg coats. Nature 456, 653–657. https://doi.org/10.1038/nature07599.

9. Raj, I., Sadat Al Hosseini, H., Dioguardi, E., Nishimura, K., Han, L., Villa, A., de Sanctis, D., and Jovine, L. (2017). Structural Basis of Egg Coat-Sperm Recognition at Fertilization. Cell 169, 1315–1326.e17. https://doi.org/10.1016/j.cell.2017.05.033.

10. Rankin, T.L., Coleman, J.S., Epifano, O., Hoodbhoy, T., Turner, S.G., Castle, P.E., Lee, E., Gore-Langton, R., and Dean, J. (2003). Fertility and taxon-specific sperm binding persist after replacement of mouse sperm receptors with human homologs. Dev. Cell 5, 33–43. https://doi.org/10.1016/S1534-5807(03)00195-3.

11. Lindsay, L.L., and Hedrick, J.L. (2004). Proteolysis of *Xenopus laevis* egg envelope ZPA triggers envelope hardening. Biochem. Biophys. Res. Commun. 324, 648–654. https://doi.org/10.1016/j.bbrc.2004.09.099.

12. Goudet, G., Mugnier, S., Callebaut, I., and Monget, P. (2008). Phylogenetic analysis and identification of pseudogenes reveal a progressive loss of zona pellucida genes during evolution of vertebrates. Biol. Reprod. 78, 796–806. https://doi.org/10.1095/biolreprod.107.064568.

13. Jovine, L., Qi, H., Williams, Z., Litscher, E., and Wassarman, P.M. (2002). The ZP domain is a conserved module for polymerization of extracellular proteins. Nat. Cell Biol. 4, 457–461. https://doi.org/10.1038/ncb802.

14. Jovine, L., Darie, C.C., Litscher, E.S., and Wassarman, P.M. (2005). Zona pellucida domain proteins. Annu. Rev. Biochem. 74, 83–114. https://doi.org/10.1146/annurev.biochem.74.082803.133039.

15. Aagaard, J.E., Vacquier, V.D., MacCoss, M.J., and Swanson, W.J. (2010). ZP domain proteins in the abalone egg coat include a paralog of VERL under positive selection that binds lysin and 18-kDa sperm proteins. Mol. Biol. Evol. 27, 193–203. https://doi.org/10.1093/molbev/msp221.

16. Litscher, E.S., and Wassarman, P.M. (2020). Zona Pellucida Proteins, Fibrils, and Matrix. Annu. Rev. Biochem. 89, 695–715. https://doi.org/10.1146/annurev-biochem-011520-105310.

17. Litscher, E.S., Qi, H., and Wassarman, P.M. (1999). Mouse Zona Pellucida Glycoproteins mZP2 and mZP3 Undergo Carboxy-Terminal Proteolytic Processing in Growing Oocytes. Biochemistry 38, 12280–12287. https://doi.org/10.1021/bi991154y.

18. Brunati, M., Perucca, S., Han, L., Cattaneo, A., Consolato, F., Andolfo, A., Schaeffer, C., Olinger, E., Peng, J., Santambrogio, S., et al. (2015). The serine protease hepsin mediates urinary secretion and polymerisation of Zona Pellucida domain protein uromodulin. Elife 4, e08887. https://doi.org/10.7554/eLife.08887.

19. Jovine, L., Qi, H., Williams, Z., Litscher, E.S., and Wassarman, P.M. (2004). A duplicated motif controls assembly of zona pellucida domain proteins. Proc. Natl. Acad. Sci. U. S. A. 101, 5922–5927. https://doi.org/10.1073/pnas.0401600101.

20. Bokhove, M., Nishimura, K., Brunati, M., Han, L., de Sanctis, D., Rampoldi, L., and Jovine, L. (2016). A structured interdomain linker directs self-polymerization of human uromodulin. Proc. Natl. Acad. Sci. U. S. A. 113, 1552–1557. https://doi.org/10.1073/pnas.1519803113.

21. Stsiapanava, A., Xu, C., Brunati, M., Zamora-Caballero, S., Schaeffer, C., Bokhove, M., Han, L., Hebert, H., Carroni, M., Yasumasu, S., et al. (2020). Cryo-EM structure of native human uromodulin, a zona pellucida module polymer. EMBO J. 39, e106807. https://doi.org/10.15252/embj.2020106807.

22. Stanisich, J.J., Zyla, D.S., Afanasyev, P., Xu, J., Kipp, A., Olinger, E., Devuyst, O., Pilhofer, M., Boehringer, D., and Glockshuber, R. (2020). The cryo-EM structure of the human uromodulin filament core reveals a unique assembly mechanism. Elife 9, e60265. https://doi.org/10.7554/eLife.60265.

23. Avella, M.A., Baibakov, B., and Dean, J. (2014). A single domain of the ZP2 zona pellucida protein mediates gamete recognition in mice and humans. J. Cell Biol. 205, 801–809. https://doi.org/10.1083/jcb.201404025.

24. Tokuhiro, K., and Dean, J. (2018). Glycan-Independent Gamete Recognition Triggers Egg Zinc Sparks and ZP2 Cleavage to Prevent Polyspermy. Dev. Cell 46, 627–640.e5. https://doi.org/10.1016/j.devcel.2018.07.020.

25. Schroeder, A.C., Schultz, R.M., Kopf, G.S., Taylor, F.R., Becker, R.B., and Eppig, J.J. (1990). Fetuin inhibits zona pellucida hardening and conversion of ZP2 to ZP2f during spontaneous mouse oocyte maturation in vitro in the absence of serum. Biol. Reprod. 43, 891–897. https://doi.org/10.1095/biolreprod43.5.891.

26. Nishimura, K., Dioguardi, E., Nishio, S., Villa, A., Han, L., Matsuda, T., and Jovine, L. (2019). Molecular basis of egg coat cross-linking sheds light on ZP1-associated female infertility. Nat. Commun. 10, 3086. https://doi.org/10.1038/s41467-019-10931-5.

27. Tian, J., Gong, H., Thomsen, G.H., and Lennarz, W.J. (1997). Gamete interactions in *Xenopus laevis*: identification of sperm binding glycoproteins in the egg vitelline envelope. J. Cell Biol. 136, 1099–1108. https://doi.org/10.1083/jcb.136.5.1099.

28. Rankin, T.L., O’Brien, M., Lee, E., Wigglesworth, K., Eppig, J., and Dean, J. (2001). Defective zonae pellucidae in *Zp2*-null mice disrupt folliculogenesis, fertility and development. Development 128, 1119–1126. https://doi.org/10.1242/dev.128.7.1119.

29. Yasumasu, S., Kawaguchi, M., Ouchi, S., Sano, K., Murata, K., Sugiyama, H., Akema, T., and Iuchi, I. (2010). Mechanism of egg envelope digestion by hatching enzymes, HCE and LCE in medaka, *Oryzias latipes*. J. Biochem. 148, 439–448. https://doi.org/10.1093/jb/mvq086.

30. Jovine, L. (2021). Using machine learning to study protein-protein interactions: From the uromodulin polymer to egg zona pellucida filaments. Mol. Reprod. Dev. 88, 686–693. https://doi.org/10.1002/mrd.23538.

31. Zhao, M., Boja, E.S., Hoodbhoy, T., Nawrocki, J., Kaufman, J.B., Kresge, N., Ghirlando, R., Shiloach, J., Pannell, L., Levine, R.L., et al. (2004). Mass spectrometry analysis of recombinant human ZP3 expressed in glycosylation-deficient CHO cells. Biochemistry 43, 12090–12104. https://doi.org/10.1021/bi048958k.

32. Saito, T., Bokhove, M., Croci, R., Zamora-Caballero, S., Han, L., Letarte, M., de Sanctis, D., and Jovine, L. (2017). Structural Basis of the Human Endoglin-BMP9 Interaction: Insights into BMP Signaling and HHT1. Cell Rep. 19, 1917–1928. https://doi.org/10.1016/j.celrep.2017.05.011.

33. Legan, P.K., Lukashkina, V.A., Goodyear, R.J., Lukashkin, A.N., Verhoeven, K., Van Camp, G., Russell, I.J., and Richardson, G.P. (2005). A deafness mutation isolates a second role for the tectorial membrane in hearing. Nat. Neurosci. 8, 1035–1042. https://doi.org/10.1038/nn1496.

34. Suzuki, K., Tatebe, N., Kojima, S., Hamano, A., Orita, M., and Yonezawa, N. (2015). The Hinge Region of Bovine Zona Pellucida Glycoprotein ZP3 Is Involved in the Formation of the Sperm-Binding Active ZP3/ZP4 Complex. Biomolecules 5, 3339–3353. https://doi.org/10.3390/biom5043339.

35. Chalabi, S., Panico, M., Sutton-Smith, M., Haslam, S.M., Patankar, M.S., Lattanzio, F.A., Morris, H.R., Clark, G.F., and Dell, A. (2006). Differential O-glycosylation of a conserved domain expressed in murine and human ZP3. Biochemistry 45, 637–647. https://doi.org/10.1021/bi0512804.

36. Terwilliger, T.C., Afonine, P.V., Liebschner, D., Croll, T.I., McCoy, A.J., Oeffner, R.D., Williams, C.J., Poon, B.K., Richardson, J.S., Read, R.J., et al. (2023). Accelerating crystal structure determination with iterative AlphaFold prediction. Acta Crystallogr. D Struct. Biol. 79, 234–244. https://doi.org/10.1107/S205979832300102X.

37. Greve, J.M., and Wassarman, P.M. (1985). Mouse egg extracellular coat is a matrix of interconnected filaments possessing a structural repeat. J. Mol. Biol. 181, 253–264. https://doi.org/10.1016/0022-2836(85)90089-0.

38. Wassarman, P.M., and Mortillo, S. (1991). Structure of the mouse egg extracellular coat, the zona pellucida. Int. Rev. Cytol. 130, 85–110. https://doi.org/10.1016/s0074-7696(08)61502-8.

39. Boja, E.S., Hoodbhoy, T., Fales, H.M., and Dean, J. (2003). Structural characterization of native mouse zona pellucida proteins using mass spectrometry. J. Biol. Chem. 278, 34189–34202. https://doi.org/10.1074/jbc.M304026200.

40. Bokhove, M., and Jovine, L. (2018). Structure of Zona Pellucida Module Proteins. Curr. Top. Dev. Biol. 130, 413–442. https://doi.org/10.1016/bs.ctdb.2018.02.007.

41. Feng, J.-M., Tian, H.-F., Hu, Q.-M., Meng, Y., and Xiao, H.-B. (2018). Evolution and multiple origins of zona pellucida genes in vertebrates. Biol. Open 7, bio036137. https://doi.org/10.1242/bio.036137.

42. Liu, C., Litscher, E.S., Mortillo, S., Sakai, Y., Kinloch, R.A., Stewart, C.L., and Wassarman, P.M. (1996). Targeted disruption of the *mZP3* gene results in production of eggs lacking a zona pellucida and infertility in female mice. Proc. Natl. Acad. Sci. U. S. A. 93, 5431–5436. https://doi.org/10.1073/pnas.93.11.5431.

43. Rankin, T., Familari, M., Lee, E., Ginsberg, A., Dwyer, N., Blanchette-Mackie, J., Drago, J., Westphal, H., and Dean, J. (1996). Mice homozygous for an insertional mutation in the *Zp3* gene lack a zona pellucida and are infertile. Development 122, 2903–2910. https://doi.org/10.1242/dev.122.9.2903.

44. Rankin, T., Talbot, P., Lee, E., and Dean, J. (1999). Abnormal zonae pellucidae in mice lacking ZP1 result in early embryonic loss. Development 126, 3847–3855. https://doi.org/10.1242/dev.126.17.3847.

45. Zeng, M.-H., Wang, Y., Huang, H.-L., Quan, R.-P., Yang, J.-T., Guo, D., Sun, Y., Lv, C., Li, T.-Y., Wang, L., et al. (2021). *Zp4* is completely dispensable for fertility in female rats. Biol. Reprod. 104, 1282–1291. https://doi.org/10.1093/biolre/ioab047.

46. Lamas-Toranzo, I., Fonseca Balvís, N., Querejeta-Fernández, A., Izquierdo-Rico, M.J., González-Brusi, L., Lorenzo, P.L., García-Rebollar, P., Avilés, M., and Bermejo-Álvarez, P. (2019). ZP4 confers structural properties to the zona pellucida essential for embryo development. Elife 8, e48904. https://doi.org/10.7554/eLife.48904.

47. Murata, K., and Kinoshita, M. (2022). Targeted deletion of liver-expressed Choriogenin L results in the production of soft eggs and infertility in medaka, *Oryzias latipes*. Zoological Lett 8, 1. https://doi.org/10.1186/s40851-021-00185-9.

48. Birk, D.S., Onose, S., Kinoshita, M., and Murata, K. (2022). Medaka, Oryzias latipes, egg envelopes are created by ovarian-expressed ZP proteins and liver-expressed choriogenins. Zoological Lett 8, 11. https://doi.org/10.1186/s40851-022-00194-2.

49. Yokokawa, R., Watanabe, K., Kanda, S., Nishino, Y., Yasumasu, S., and Sano, K. (2023). Egg envelope formation of medaka *Oryzias latipes* requires ZP proteins originating from both the liver and ovary. J. Biol. Chem. 299, 104600. https://doi.org/10.1016/j.jbc.2023.104600.

50. Wassarman, P.M., Qi, H., and Litscher, E.S. (1997). Mutant female mice carrying a single *mZP3* allele produce eggs with a thin zona pellucida, but reproduce normally. Proc. Biol. Sci. 264, 323–328. https://doi.org/10.1098/rspb.1997.0046.

51. Darie, C.C., Janssen, W.G., Litscher, E.S., and Wassarman, P.M. (2008). Purified trout egg vitelline envelope proteins VEβ and VEγ polymerize into homomeric fibrils from dimers *in vitro*. Biochim. Biophys. Acta 1784, 385–392. https://doi.org/10.1016/j.bbapap.2007.10.011.

52. Litscher, E.S., Janssen, W.G., Darie, C.C., and Wassarman, P.M. (2008). Purified mouse egg zona pellucida glycoproteins polymerize into homomeric fibrils under non-denaturing conditions. J. Cell. Physiol. 214, 153–157. https://doi.org/10.1002/jcp.21174.

53. Nishio, S., Kohno, Y., Iwata, Y., Arai, M., Okumura, H., Oshima, K., Nadano, D., and Matsuda, T. (2014). Glycosylated chicken ZP2 accumulates in the egg coat of immature oocytes and remains localized to the germinal disc region of mature eggs. Biol. Reprod. 91, 107, 1–10. https://doi.org/10.1095/biolreprod.114.119826.

54. Familiari, G., Nottola, S.A., Macchiarelli, G., Micara, G., Aragona, C., and Motta, P.M. (1992). Human zona pellucida during in vitro fertilization: an ultrastructural study using saponin, ruthenium red, and osmium-thiocarbohydrazide. Mol. Reprod. Dev. 32, 51–61. https://doi.org/10.1002/mrd.1080320109.

55. Fahrenkamp, E., Algarra, B., and Jovine, L. (2020). Mammalian egg coat modifications and the block to polyspermy. Mol. Reprod. Dev. 87, 326–340. https://doi.org/10.1002/mrd.23320.

56. Maddirevula, S., Coskun, S., Al-Qahtani, M., Aboyousef, O., Alhassan, S., Aldeery, M., and Alkuraya, F.S. (2022). *ASTL* is mutated in female infertility. Hum. Genet. 141, 49–54. https://doi.org/10.1007/s00439-021-02388-8.

57. Iuchi, I., Ha, C.-R., and Matsuda, K. (1995). Chorion hardening in medaka (*Oryzias latipes*) egg. The Fish Biology Journal MEDAKA. 7, 15–20. https://doi.org/10.18999/fisbjm.7.15.

58. Cui, Z., Lu, Y., Miao, Y., Dai, X., Zhang, Y., and Xiong, B. (2022). Transglutaminase 2 crosslinks zona pellucida glycoprotein 3 to prevent polyspermy. Cell Death Differ. 29, 1466–1473. https://doi.org/10.1038/s41418-022-00933-0.

59. Hasegawa, A., Koyama, K., Okazaki, Y., Sugimoto, M., and Isojima, S. (1994). Amino acid sequence of a porcine zona pellucida glycoprotein ZP4 determined by peptide mapping and cDNA cloning. J. Reprod. Fertil. 100, 245–255. https://doi.org/10.1530/jrf.0.1000245.

60. Kwamoto, K., Ikeda, K., Yonezawa, N., Noguchi, S., Kudo, K., Hamano, S., Kuwayama, M., and Nakano, M. (1999). Disulfide formation in bovine zona pellucida glycoproteins during fertilization: evidence for the involvement of cystine cross-linkages in hardening of the zona pellucida. Reproduction 117, 395–402. https://doi.org/10.1530/jrf.0.1170395.

61. Gahlay, G., Gauthier, L., Baibakov, B., Epifano, O., and Dean, J. (2010). Gamete recognition in mice depends on the cleavage status of an egg’s zona pellucida protein. Science 329, 216–219. https://doi.org/10.1126/science.1188178.

62. Scarpeci, S.L., Sanchez, M.L., and Cabada, M.O. (2008). Cellular origin of the *Bufo arenarum* sperm receptor gp75, a ZP2 family member: its proteolysis after fertilization. Biol. Cell 100, 219–230. https://doi.org/10.1042/BC20070052.

63. Baibakov, B., Boggs, N.A., Yauger, B., Baibakov, G., and Dean, J. (2012). Human sperm bind to the N-terminal domain of ZP2 in humanized zonae pellucidae in transgenic mice. J. Cell Biol. 197, 897–905. https://doi.org/10.1083/jcb.201203062.

64. Wessel, G.M., and Wong, J.L. (2009). Cell surface changes in the egg at fertilization. Mol. Reprod. Dev. 76, 942–953. https://doi.org/10.1002/mrd.21090.

65. Tekleyohans, D.G., Mao, Y., Kägi, C., Stierhof, Y.-D., and Groß-Hardt, R. (2017). Polyspermy barriers: a plant perspective. Curr. Opin. Plant Biol. 35, 131–137. https://doi.org/10.1016/j.pbi.2016.11.012.

66. Litscher, E.S., and Wassarman, P.M. (2020). Zona pellucida genes and proteins and human fertility. Trends Dev. Biol. 13, 21–33.

67. Loeuillet, C., Dhellemmes, M., Cazin, C., Kherraf, Z.-E., Fourati Ben Mustapha, S., Zouari, R., Thierry-Mieg, N., Arnoult, C., and Ray, P.F. (2022). A recurrent *ZP1* variant is responsible for oocyte maturation defect with degenerated oocytes in infertile females. Clin. Genet. 102, 22–29. https://doi.org/10.1111/cge.14144.

68. Pujalte, M., Camo, M., Celton, N., Attencourt, C., Lefranc, E., Jedraszak, G., and Scheffler, F. (2023). A *ZP1* gene mutation in a patient with empty follicle syndrome: A case report and literature review. Eur. J. Obstet. Gynecol. Reprod. Biol. 280, 193–197. https://doi.org/10.1016/j.ejogrb.2022.12.011.

69. Chu, K., He, Y., Wang, L., Ji, Y., Hao, M., Pang, W., Liu, Y., Sun, N., Yang, F., and Li, W. (2020). Novel ZP1 pathogenic variants identified in an infertile patient and a successful live birth following ICSI treatment. Clin. Genet. 97, 787–788. https://doi.org/10.1111/cge.13693.

70. Hou, M., Zhu, L., Jiang, J., Liu, Z., Li, Z., Jia, W., Hu, J., Zhou, X., Zhang, D., Luo, Y., et al. (2022). Novel Heterozygous Mutations in *ZP2* Cause Abnormal Zona Pellucida and Female Infertility. Reprod. Sci. 29, 3047–3054. https://doi.org/10.1007/s43032-022-00958-3.

71. Zhang, Z., Guo, Q., Jia, L., Zhou, C., He, S., Fang, C., Zhang, M., Sun, P., Zeng, Z., Wang, M., et al. (2022). A novel gene mutation in *ZP3* loop region identified in patients with empty follicle syndrome. Hum. Mutat. 43, 180–188. https://doi.org/10.1002/humu.24297.

72. Wei, X., Li, Y., Liu, Q., Liu, W., Yan, X., Zhu, X., Zhou, D., Tian, Y., Zhang, F., Li, N., et al. (2022). Mutations in *ZP4* are associated with abnormal zona pellucida and female infertility. J. Clin. Pathol. 75, 201–204. https://doi.org/10.1136/jclinpath-2020-207170.

73. Tunyasuvunakool, K., Adler, J., Wu, Z., Green, T., Zielinski, M., Žídek, A., Bridgland, A., Cowie, A., Meyer, C., Laydon, A., et al. (2021). Highly accurate protein structure prediction for the human proteome. Nature 596, 590–596. https://doi.org/10.1038/s41586-021-03828-1.

74. Sugiyama, H., Murata, K., Iuchi, I., Nomura, K., and Yamagami, K. (1999). Formation of mature egg envelope subunit proteins from their precursors (choriogenins) in the fish, *Oryzias latipes*: loss of partial C-terminal sequences of the choriogenins. J. Biochem. 125, 469–475. https://doi.org/10.1093/oxfordjournals.jbchem.a022310.

75. Järvå, M.A., Lingford, J.P., John, A., Soler, N.M., Scott, N.E., and Goddard-Borger, E.D. (2020). Trefoil factors share a lectin activity that defines their role in mucus. Nat. Commun. 11, 2265. https://doi.org/10.1038/s41467-020-16223-7.

76. East, I.J., and Dean, J. (1984). Monoclonal antibodies as probes of the distribution of ZP-2, the major sulfated glycoprotein of the murine zona pellucida. J. Cell Biol. 98, 795–800. https://doi.org/10.1083/jcb.98.3.795.

77. Noda, T., Lu, Y., Fujihara, Y., Oura, S., Koyano, T., Kobayashi, S., Matzuk, M.M., and Ikawa, M. (2020). Sperm proteins SOF1, TMEM95, and SPACA6 are required for sperm-oocyte fusion in mice. Proc. Natl. Acad. Sci. U. S. A. 117, 11493–11502. https://doi.org/10.1073/pnas.1922650117.

78. Sasado, T., Tanaka, M., Kobayashi, K., Sato, T., Sakaizumi, M., and Naruse, K. (2010). The National BioResource Project Medaka (NBRP Medaka): an integrated bioresource for biological and biomedical sciences. Exp. Anim. 59, 13–23. https://doi.org/10.1538/expanim.59.13.

79. Hekkelman, M.L., de Vries, I., Joosten, R.P., and Perrakis, A. (2022). AlphaFill: enriching AlphaFold models with ligands and cofactors. Nat. Methods, 1–9. https://doi.org/10.1038/s41592-022-01685-y.

80. Jumper, J., Evans, R., Pritzel, A., Green, T., Figurnov, M., Ronneberger, O., Tunyasuvunakool, K., Bates, R., Žídek, A., Potapenko, A., et al. (2021). Highly accurate protein structure prediction with AlphaFold. Nature 596, 583–589. https://doi.org/10.1038/s41586-021-03819-2.

81. Arnold, M.J. (2021). AlphaPickle. https://doi.org/10.5281/zenodo.5708709.

82. Maier, J.A., Martinez, C., Kasavajhala, K., Wickstrom, L., Hauser, K.E., and Simmerling, C. (2015). ff14SB: Improving the Accuracy of Protein Side Chain and Backbone Parameters from ff99SB. J. Chem. Theory Comput. 11, 3696–3713. https://doi.org/10.1021/acs.jctc.5b00255.

83. Vonrhein, C., Flensburg, C., Keller, P., Sharff, A., Smart, O., Paciorek, W., Womack, T., and Bricogne, G. (2011). Data processing and analysis with the *autoPROC* toolbox. Acta Crystallogr. D Biol. Crystallogr. 67, 293–302. https://doi.org/10.1107/S0907444911007773.

84. Vonrhein, C., Tickle, I.J., Flensburg, C., Keller, P., Paciorek, W., Sharff, A., Bricogne, G., and IUCr (2018). Advances in automated data analysis and processing within autoPROC, combined with improved characterisation, mitigation and visualisation of the anisotropy of diffraction limits using STARANISO. Acta Crystallogr. A 74, a360–a360. https://doi.org/10.1107/S010876731809640X.

85. Vonrhein, C., Blanc, E., Roversi, P., and Bricogne, G. (2007). Automated structure solution with autoSHARP. Methods Mol. Biol. 364, 215–230. https://doi.org/10.1385/1-59745-266-1:215.

86. Barson, G., and Griffiths, E. (2016). SeqTools: visual tools for manual analysis of sequence alignments. BMC Res. Notes 9, 39. https://doi.org/10.1186/s13104-016-1847-3.

87. Smart, O.S., Womack, T.O., Flensburg, C., Keller, P., Paciorek, W., Sharff, A., Vonrhein, C., and Bricogne, G. (2012). Exploiting structure similarity in refinement: automated NCS and target-structure restraints in *BUSTER*. Acta Crystallogr. D Biol. Crystallogr. 68, 368–380. https://doi.org/10.1107/S0907444911056058.

88. Brunger, A.T. (2007). Version 1.2 of the Crystallography and NMR system. Nat. Protoc. 2, 2728–2733. https://doi.org/10.1038/nprot.2007.406.

89. Mirdita, M., Schütze, K., Moriwaki, Y., Heo, L., Ovchinnikov, S., and Steinegger, M. (2022). ColabFold: making protein folding accessible to all. Nat. Methods 19, 679–682. https://doi.org/10.1038/s41592-022-01488-1.

90. Casañal, A., Lohkamp, B., and Emsley, P. (2020). Current developments in *Coot* for macromolecular model building of Electron Cryo-microscopy and Crystallographic Data. Protein Sci. 29, 1069–1078. https://doi.org/10.1002/pro.3791.

91. Naito, Y., Hino, K., Bono, H., and Ui-Tei, K. (2015). CRISPRdirect: software for designing CRISPR/Cas guide RNA with reduced off-target sites. Bioinformatics 31, 1120–1123. https://doi.org/10.1093/bioinformatics/btu743.

92. Punjani, A., Rubinstein, J.L., Fleet, D.J., and Brubaker, M.A. (2017). cryoSPARC: algorithms for rapid unsupervised cryo-EM structure determination. Nat. Methods 14, 290–296. https://doi.org/10.1038/nmeth.4169.

93. Delgado, J., Radusky, L.G., Cianferoni, D., and Serrano, L. (2019). FoldX 5.0: working with RNA, small molecules and a new graphical interface. Bioinformatics 35, 4168–4169. https://doi.org/10.1093/bioinformatics/btz184.

94. Bohne-Lang, A., and von der Lieth, C.-W. (2005). GlyProt: *in silico* glycosylation of proteins. Nucleic Acids Res. 33, W214–W219. https://doi.org/10.1093/nar/gki385.

95. Blum, M., Chang, H.-Y., Chuguransky, S., Grego, T., Kandasaamy, S., Mitchell, A., Nuka, G., Paysan-Lafosse, T., Qureshi, M., Raj, S., et al. (2021). The InterPro protein families and domains database: 20 years on. Nucleic Acids Res. 49, D344–D354. https://doi.org/10.1093/nar/gkaa977.

96. Croll, T.I. (2018). *ISOLDE*: a physically realistic environment for model building into low-resolution electron-density maps. Acta Crystallogr. D Struct. Biol. 74, 519–530. https://doi.org/10.1107/S2059798318002425.

97. Katoh, K., and Standley, D.M. (2013). MAFFT multiple sequence alignment software version 7: improvements in performance and usability. Mol. Biol. Evol. 30, 772–780. https://doi.org/10.1093/molbev/mst010.

98. Necci, M., Piovesan, D., Clementel, D., Dosztányi, Z., and Tosatto, S.C.E. (2020). MobiDB-lite 3.0: fast consensus annotation of intrinsic disorder flavors in proteins. Bioinformatics 36, 5533–5534. https://doi.org/10.1093/bioinformatics/btaa1045.

99. Webb, B., and Sali, A. (2016). Comparative Protein Structure Modeling Using MODELLER. Curr. Protoc. Protein Sci. 86, 2.9.1–2.9.37. https://doi.org/10.1002/cpps.20.

100. Chen, V.B., Arendall, W.B., 3rd, Headd, J.J., Keedy, D.A., Immormino, R.M., Kapral, G.J., Murray, L.W., Richardson, J.S., and Richardson, D.C. (2010). *MolProbity*: all-atom structure validation for macromolecular crystallography. Acta Crystallogr. D Biol. Crystallogr. 66, 12–21. https://doi.org/10.1107/S0907444909042073.

101. Brown, N.P., Leroy, C., and Sander, C. (1998). MView: a web-compatible database search or multiple alignment viewer. Bioinformatics 14, 380–381. https://doi.org/10.1093/bioinformatics/14.4.380.

102. Edgar, R.C. (2022). Muscle5: High-accuracy alignment ensembles enable unbiased assessments of sequence homology and phylogeny. Nat. Commun. 13, 6968. https://doi.org/10.1038/s41467-022-34630-w.

103. de Beer, T.A.P., Berka, K., Thornton, J.M., and Laskowski, R.A. (2014). PDBsum additions. Nucleic Acids Res. 42, D292–D296. https://doi.org/10.1093/nar/gkt940.

104. McCoy, A.J., Grosse-Kunstleve, R.W., Adams, P.D., Winn, M.D., Storoni, L.C., and Read, R.J. (2007). *Phaser* crystallographic software. J. Appl. Crystallogr. 40, 658–674. https://doi.org/10.1107/S0021889807021206.

105. Terwilliger, T.C., Grosse-Kunstleve, R.W., Afonine, P.V., Moriarty, N.W., Zwart, P.H., Hung, L.W., Read, R.J., and Adams, P.D. (2008). Iterative model building, structure refinement and density modification with the *PHENIX AutoBuild* wizard. Acta Crystallogr. D Biol. Crystallogr. 64, 61–69. https://doi.org/10.1107/S090744490705024X.

106. Terwilliger, T.C., Adams, P.D., Read, R.J., McCoy, A.J., Moriarty, N.W., Grosse-Kunstleve, R.W., Afonine, P.V., Zwart, P.H., and Hung, L.W. (2009). Decision-making in structure solution using Bayesian estimates of map quality: the *PHENIX AutoSol* wizard. Acta Crystallogr. D Biol. Crystallogr. 65, 582–601. https://doi.org/10.1107/S0907444909012098.

107. Afonine, P.V., Grosse-Kunstleve, R.W., Echols, N., Headd, J.J., Moriarty, N.W., Mustyakimov, M., Terwilliger, T.C., Urzhumtsev, A., Zwart, P.H., and Adams, P.D. (2012). Towards automated crystallographic structure refinement with *phenix.refine*. Acta Crystallogr. D Biol. Crystallogr. 68, 352–367. https://doi.org/10.1107/S0907444912001308.

108. Terwilliger, T.C., Sobolev, O.V., Afonine, P.V., and Adams, P.D. (2018). Automated map sharpening by maximization of detail and connectivity. Acta Crystallogr. D Struct. Biol. 74, 545–559. https://doi.org/10.1107/S2059798318004655.

109. Krissinel, E., and Henrick, K. (2007). Inference of macromolecular assemblies from crystalline state. J. Mol. Biol. 372, 774–797. https://doi.org/10.1016/j.jmb.2007.05.022.

110. Agirre, J., Iglesias-Fernandez, J., Rovira, C., Davies, G.J., Wilson, K.S., and Cowtan, K.D. (2015). Privateer: software for the conformational validation of carbohydrate structures. Nat. Struct. Mol. Biol. 22, 833–834. https://doi.org/10.1038/nsmb.3115.

111. Mastronarde, D.N. (2005). Automated electron microscope tomography using robust prediction of specimen movements. J. Struct. Biol. 152, 36–51. https://doi.org/10.1016/j.jsb.2005.07.007.

112. Frishman, D., and Argos, P. (1995). Knowledge-based protein secondary structure assignment. Proteins: Struct. Funct. Bioinf. 23, 566–579. https://doi.org/10.1002/prot.340230412.

113. Pettersen, E.F., Goddard, T.D., Huang, C.C., Couch, G.S., Greenblatt, D.M., Meng, E.C., and Ferrin, T.E. (2004). UCSF Chimera — a visualization system for exploratory research and analysis. J. Comput. Chem. 25, 1605–1612. https://doi.org/10.1002/jcc.20084.

114. Meng, E.C., Pettersen, E.F., Couch, G.S., Huang, C.C., and Ferrin, T.E. (2006). Tools for integrated sequence-structure analysis with UCSF Chimera. BMC Bioinformatics 7, 339. https://doi.org/10.1186/1471-2105-7-339.

115. Pettersen, E.F., Goddard, T.D., Huang, C.C., Meng, E.C., Couch, G.S., Croll, T.I., Morris, J.H., and Ferrin, T.E. (2021). UCSF ChimeraX: Structure visualization for researchers, educators, and developers. Protein Sci. 30, 70–82. https://doi.org/10.1002/pro.3943.

116. Kabsch, W. (2010). XDS. Acta Crystallogr. D Biol. Crystallogr. 66, 125–132. https://doi.org/10.1107/S0907444909047337.

117. Krieger, E., Joo, K., Lee, J., Lee, J., Raman, S., Thompson, J., Tyka, M., Baker, D., and Karplus, K. (2009). Improving physical realism, stereochemistry, and side-chain accuracy in homology modeling: Four approaches that performed well in CASP8. Proteins 77 Suppl. 9, 114–122. https://doi.org/10.1002/prot.22570.

118. Qi, H., Williams, Z., and Wassarman, P.M. (2002). Secretion and assembly of zona pellucida glycoproteins by growing mouse oocytes microinjected with epitope-tagged cDNAs for mZP2 and mZP3. Mol. Biol. Cell 13, 530–541. https://doi.org/10.1091/mbc.01-09-0440.

119. Aricescu, A.R., Lu, W., and Jones, E.Y. (2006). A time- and cost-efficient system for high-level protein production in mammalian cells. Acta Crystallogr. D Biol. Crystallogr. 62, 1243–1250. https://doi.org/10.1107/S0907444906029799.

120. Bokhove, M., Sadat Al Hosseini, H., Saito, T., Dioguardi, E., Gegenschatz-Schmid, K., Nishimura, K., Raj, I., de Sanctis, D., Han, L., and Jovine, L. (2016). Easy mammalian expression and crystallography of maltose-binding protein-fused human proteins. J. Struct. Biol. 194, 1–7. https://doi.org/10.1016/j.jsb.2016.01.016.

121. Briggs, D.C., and Hohenester, E. (2018). Structural Basis for the Initiation of Glycosaminoglycan Biosynthesis by Human Xylosyltransferase 1. Structure 26, 801–809.e3. https://doi.org/10.1016/j.str.2018.03.014.

122. Pulido, D., Hussain, S.-A., and Hohenester, E. (2017). Crystal Structure of the Heterotrimeric Integrin-Binding Region of Laminin-111. Structure 25, 530–535. https://doi.org/10.1016/j.str.2017.01.002.

123. Han, K. (1996). An efficient DDAB-mediated transfection of *Drosophila* S2 cells. Nucleic Acids Res. 24, 4362–4363. https://doi.org/10.1093/nar/24.21.4362.

124. Karmilin, K., Schmitz, C., Kuske, M., Körschgen, H., Olf, M., Meyer, K., Hildebrand, A., Felten, M., Fridrich, S., Yiallouros, I., et al. (2019). Mammalian plasma fetuin-B is a selective inhibitor of ovastacin and meprin metalloproteinases. Sci. Rep. 9, 546. https://doi.org/10.1038/s41598-018-37024-5.

125. Gorrec, F. (2015). The MORPHEUS II protein crystallization screen. Acta Crystallogr. Sect. F Struct. Biol. Cryst. Commun. 71, 831–837. https://doi.org/10.1107/S2053230X1500967X.

126. Newman, J., Egan, D., Walter, T.S., Meged, R., Berry, I., Ben Jelloul, M., Sussman, J.L., Stuart, D.I., and Perrakis, A. (2005). Towards rationalization of crystallization screening for small- to medium-sized academic laboratories: the PACT/JCSG strategy. Acta Crystallogr. D Biol. Crystallogr. 61, 1426–1431. https://doi.org/10.1107/s0907444905024984.

127. Mueller, U., Darowski, N., Fuchs, M.R., Förster, R., Hellmig, M., Paithankar, K.S., Pühringer, S., Steffien, M., Zocher, G., and Weiss, M.S. (2012). Facilities for macromolecular crystallography at the Helmholtz-Zentrum Berlin. J. Synchrotron Radiat. 19, 442–449. https://doi.org/10.1107/S0909049512006395.

128. Nurizzo, D., Mairs, T., Guijarro, M., Rey, V., Meyer, J., Fajardo, P., Chavanne, J., Biasci, J.C., McSweeney, S., and Mitchell, E. (2006). The ID23-1 structural biology beamline at the ESRF. J. Synchrotron Radiat. 13, 227–238. https://doi.org/10.1107/S0909049506004341.

129. de Sanctis, D., Beteva, A., Caserotto, H., Dobias, F., Gabadinho, J., Giraud, T., Gobbo, A., Guijarro, M., Lentini, M., Lavault, B., et al. (2012). ID29: a high-intensity highly automated ESRF beamline for macromolecular crystallography experiments exploiting anomalous scattering. J. Synchrotron Radiat. 19, 455–461. https://doi.org/10.1107/S0909049512009715.

130. McCarthy, A.A., Barrett, R., Beteva, A., Caserotto, H., Dobias, F., Felisaz, F., Giraud, T., Guijarro, M., Janocha, R., Khadrouche, A., et al. (2018). ID30B - a versatile beamline for macromolecular crystallography experiments at the ESRF. J. Synchrotron Radiat. 25, 1249–1260. https://doi.org/10.1107/S1600577518007166.

131. Allan, D.R., Collins, S.P., Evans, G., Hall, D., McAuley, K., Owen, R.L., Sorensen, T., Tang, C.C., von Delft, F., Wagner, A., et al. (2015). Status of the crystallography beamlines at Diamond Light Source. Eur. Phys. J. Plus 130, 56. https://doi.org/10.1140/epjp/i2015-15056-x.

132. Abrahams, J.P., and Leslie, A.G. (1996). Methods used in the structure determination of bovine mitochondrial F_1_ ATPase. Acta Crystallogr. D Biol. Crystallogr. 52, 30–42. https://doi.org/10.1107/S0907444995008754.

133. Cowtan, K. (2006). The Buccaneer software for automated model building. 1. Tracing protein chains. Acta Crystallogr. D Biol. Crystallogr. 62, 1002–1011. https://doi.org/10.1107/S0907444906022116.

134. Cowtan, K. (2010). Recent developments in classical density modification. Acta Crystallogr. D Biol. Crystallogr. 66, 470–478. https://doi.org/10.1107/S090744490903947X.

135. Murata, K., Sugiyama, H., Yasumasu, S., Iuchi, I., Yasumasu, I., and Yamagami, K. (1997). Cloning of cDNA and estrogen-induced hepatic gene expression for choriogenin H, a precursor protein of the fish egg envelope (chorion). Proc. Natl. Acad. Sci. U. S. A. 94, 2050–2055. https://doi.org/10.1073/pnas.94.5.2050.

136. Schröder, G.F., Levitt, M., and Brunger, A.T. (2014). Deformable elastic network refinement for low-resolution macromolecular crystallography. Acta Crystallogr. D Biol. Crystallogr. 70, 2241–2255. https://doi.org/10.1107/S1399004714016496.

137. DeLaBarre, B., and Brunger, A.T. (2006). Considerations for the refinement of low-resolution crystal structures. Acta Crystallogr. D Biol. Crystallogr. 62, 923–932. https://doi.org/10.1107/S0907444906012650.

138. Sugiyama, H., Yasumasu, S., Murata, K., Iuchi, I., and Yamagami, K. (1998). The third egg envelope subunit in fish: cDNA cloning and analysis, and gene expression. Dev. Growth Differ. 40, 35–45. https://doi.org/10.1046/j.1440-169x.1998.t01-5-00005.x.

139. Punjani, A., Zhang, H., and Fleet, D.J. (2020). Non-uniform refinement: adaptive regularization improves single-particle cryo-EM reconstruction. Nat. Methods 17, 1214–1221. https://doi.org/10.1038/s41592-020-00990-8.

140. Kidmose, R.T., Juhl, J., Nissen, P., Boesen, T., Karlsen, J.L., and Pedersen, B.P. (2019). *Namdinator* - automatic molecular dynamics flexible fitting of structural models into cryo-EM and crystallography experimental maps. IUCrJ 6, 526–531. https://doi.org/10.1107/S2052252519007619.

141. Robert, X., and Gouet, P. (2014). Deciphering key features in protein structures with the new ENDscript server. Nucleic Acids Res. 42, W320–W324. https://doi.org/10.1093/nar/gku316.

142. Wilson, C.J., Choy, W.-Y., and Karttunen, M. (2022). AlphaFold2: A Role for Disordered Protein/Region Prediction? Int. J. Mol. Sci. 23, 4591. https://doi.org/10.3390/ijms23094591.

143. Jurrus, E., Engel, D., Star, K., Monson, K., Brandi, J., Felberg, L.E., Brookes, D.H., Wilson, L., Chen, J., Liles, K., et al. (2018). Improvements to the APBS biomolecular solvation software suite. Protein Sci. 27, 112–128. https://doi.org/10.1002/pro.3280.

144. Ashkenazy, H., Abadi, S., Martz, E., Chay, O., Mayrose, I., Pupko, T., and Ben-Tal, N. (2016). ConSurf 2016: an improved methodology to estimate and visualize evolutionary conservation in macromolecules. Nucleic Acids Res. 44, W344–W350. https://doi.org/10.1093/nar/gkw408.

145. Pruitt, K.D., Tatusova, T., and Maglott, D.R. (2005). NCBI Reference Sequence (RefSeq): a curated non-redundant sequence database of genomes, transcripts and proteins. Nucleic Acids Res. 33, D501–D504. https://doi.org/10.1093/nar/gki025.

146. Abbasi, F., Miyata, H., Shimada, K., Morohoshi, A., Nozawa, K., Matsumura, T., Xu, Z., Pratiwi, P., and Ikawa, M. (2018). RSPH6A is required for sperm flagellum formation and male fertility in mice. J. Cell Sci. 131, jcs221648. https://doi.org/10.1242/jcs.221648.

147. Kaneko, T., and Mashimo, T. (2015). Simple Genome Editing of Rodent Intact Embryos by Electroporation. PLoS One 10, e0142755. https://doi.org/10.1371/journal.pone.0142755.

148. Toyoda, Y., Yokoyama, M., and Hoshi, T. (1971). Studies on the fertilization of mouse eggs in vitro. I. In vitro fertilization of mouse eggs by fresh epididymal sperm. Japan. J. Anim. Reprod. 16, 147–151. https://doi.org/10.1262/jrd1955.16.147.

149. Evans, R., O’Neill, M., Pritzel, A., Antropova, N., Senior, A., Green, T., Žídek, A., Bates, R., Blackwell, S., Yim, J., et al. (2021). Protein complex prediction with AlphaFold-Multimer. bioRxiv, 2021.10.04.463034. https://doi.org/10.1101/2021.10.04.463034.

150. Okada, A., Sano, K., Nagata, K., Yasumasu, S., Ohtsuka, J., Yamamura, A., Kubota, K., Iuchi, I., and Tanokura, M. (2010). Crystal structure of zebrafish hatching enzyme 1 from the zebrafish *Danio rerio*. J. Mol. Biol. 402, 865–878. https://doi.org/10.1016/j.jmb.2010.08.023.

151. Varadi, M., Anyango, S., Deshpande, M., Nair, S., Natassia, C., Yordanova, G., Yuan, D., Stroe, O., Wood, G., Laydon, A., et al. (2022). AlphaFold Protein Structure Database: massively expanding the structural coverage of protein-sequence space with high-accuracy models. Nucleic Acids Res. 50, D439–D444. https://doi.org/10.1093/nar/gkab1061.

152. Jones, D.T., and Cozzetto, D. (2015). DISOPRED3: precise disordered region predictions with annotated protein-binding activity. Bioinformatics 31, 857–863. https://doi.org/10.1093/bioinformatics/btu744.

